# Carbon allocation strategies across Mucoromycota and the discovery of oxalic acid secretion in *Absidia*

**DOI:** 10.64898/2026.11.16.766564

**Authors:** Cristian Bolaño Losada, Boris Zimmermann, Dana Byrtusova, Sven Andreas Högfeldt, Svein Jarle Horn, Achim Kohler, Volha Shapaval

## Abstract

In this work, nine strains representing eight species, seven genera, and five families from the phylum Mucoromycota were studied for their production of biomolecules (lipids, saccharides, phosphates, organic acids, ethanol, and glycerol) under cultivation in media with different carbon-to-nitrogen (low, medium, and high). Although lipid accumulation was a common feature across the species, the results revealed also differences in carbon allocation strategies between families, both in magnitude and nature of the biomolecules produced. In general, members of Mucoraceae allocated carbon and energy toward the production of lipids, polyphosphates, organic acids, ethanol and glycerol; Mortierellaceae toward lipids, saccharides, and to a lesser extent to organic acids; Cunninghamellaceae toward lipids, phosphates, and saccharides; Lichtheimiaceae toward lipids and saccharides; and Umbelopsidaceae toward lipids, phosphates, and slightly to organic acids. In addition, genus-specific differences were observed within some families, most notably the secretion of organic acids by species of *Rhizopus* (Mucoraceae) and *Absidia* (Cunninghamellaceae).

Remarkably, *Absidia glauca* allocated carbon to the production of organic acids independently of the medium C/N ratio. This unusual observation motivated further experiments, which resulted in the identification of oxalic acid in four different strains of *A. glauca* and other five *Absidia* species. The secretion of oxalic acid likely occurs across multiple species of *Absidia*. These findings suggest that oxalic acid secretion may be a broader metabolic trait within the genus *Absidia*. To our knowledge, this is the first report of oxalic acid secretion in *Absidia* species, revealing a previously unrecognized metabolic capability within the phylum Mucoromycota.

## 1. Introduction

The accumulation of large quantities of neutral lipids in lipid bodies is a characteristic feature of Mucoromycota fungi. Since the 30s, this characteristic promoted research of several species of the phylum for the industrial production of single cell oils (Ratledge, 2006), with *Mucor circinelloides* being the most studied species using omics technologies (Hu et al., 2024; Tang et al., 2015, 2016). The main trigger of lipid accumulation is nitrogen starvation and excess of carbon, typically caused by using imbalanced media with high carbon-to-nitrogen (C/N) ratio (Wierzchowska et al., 2021). Other parameters, such as certain metal ion and phosphate concentrations (Dzurendová et al., 2021; Dzurendova, Zimmermann, Tafintseva, Kohler, Ekeberg, et al., 2020; Dzurendova, Zimmermann, Tafintseva, Kohler, Horn, et al., 2020), morphology (Shapaval et al., 2023), and different fermentation setups (Karimi et al., 2005), have been also studied to further impact lipid yield. Under nitrogen limitation, the excess of carbon is redirected towards synthesis of neural lipids for storage until the nitrogen is available again for growth. There have been reported differences in terms of lipid composition and the maximum yield of lipids’ storage, for various Mucoromycota species (Kosa, Zimmermann, et al., 2018; Zhao et al., 2021).

In contrast with neutral lipids, the accumulation of other biomolecules in Mucoromycota species such as intracellular or cell-wall saccharides, polyphosphates, carotenoids, or secretion of short-chain organic acids has been studied far less (Dzurendová et al., 2021; Dzurendova et al., 2022; Losada et al., 2025). Among these, *Rhizopus* is the only genus well studied for organic acid production, particularly fumaric acid due to its industrial importance, while other genera have not been studied at all. Polyphosphate accumulation has also received limited attention, with only few works exploring the role and affecting factors in Mucoromycota (Do Nascimento Shari’a et al., 2002; Dzurendova et al., 2021; Werner et al., 2007), despite their potential for phosphorus removal. Regarding saccharides, research has mainly focused on structural cell-wall polysaccharides such as chitin/chitosan, whereas intracellular storage saccharides (glycogen, trehalose, etc.) have been studied far less. Published studies have addressed ecological roles (Danilova et al., 2025) and biotechnological strategies, such as redirecting carbon flux towards lipid accumulation by knocking out genes involved in intracellular saccharide accumulation (Chang et al., 2023). The co-production of biomolecules of interest is crucial for maximizing the economic feasibility of fermentation-based biorefineries (Cherubini, 2010). Beyond industrial relevance, investigating the spectrum of biomolecules in Mucoromycota provides insights into their phylogeny, physiology, and ecology (Dzurendova et al., 2022).

The aim of this study was to evaluate the carbon allocation strategies of a set of nine Mucoromycota strains, representing eight species, seven genera, and four families. In our previous study (Bolaño-Losada et al., 2025), some differences on carbon allocation were already found, while in this work, we explore and discuss these differences in more detail. Vibrational spectroscopy was chosen as the central analytical approach due to its versatility in providing comprehensive biochemical profiles of fungal biomass, and it was complemented by targeted analyte-based methods. Cultivations were performed under three C/N conditions (low, standard, and high) to assess glucose consumption, biomass production, intracellular metabolite accumulation (lipids, saccharides, phosphates), and extracellular metabolite secretion (short-chain organic acids, ethanol, and glycerol). Additionally, analysis of short-chain organic acids led to the discovery of oxalic acid production by *Absidia glauca* and other species such as *A. coerulea, A. psychrophilia, A. heterospora, A. cuneospora* and *A. pseudocylindrospora*. This is the first study reporting oxalic acid production in Mucoromycota, and genus *Absidia*.

## 2. Materials and methods

### Fungal strains

The fungal strains were obtained in lyophilized form from the American Type Culture Collection (ATCC; Manassas, United States), the Czech Collection of Microorganisms (CCM; Brno, Czech Republic), the All-Russian Collection (VKM, Pushchino, Russia), the USDA-ARS Culture Collection (NRRL, Peoria, United States), and the Norwegian School of Veterinary Science (VI; Ås, Norway). According to different aims, the strains were divided into three sets: i) Set I: representative strains of Mucoromycota for the study of carbon allocation towards different biomolecules, which have been also used in our previous studies(Bolaño Losada et al., 2025; Dzurendová et al., 2021; Dzurendova et al., 2021; Dzurendova, Zimmermann, Kohler, et al., 2020; Dzurendova, Zimmermann, Tafintseva, Kohler, Ekeberg, et al., 2020; Losada et al., 2025), ii) Set II: *Absidia glauca* and *Rhizopus stolonifer* to study and compare acid production capacity and *Lichtheimia corymbifera* for comparison of phosphate accumulation, and iii) Set III: a set of *Absidia* and *Lichtheimia* species to study and compare acid production capacity and overall biomass biochemical composition. The use of Set II and Set III was designed based on results obtained from the study performed with the Set I and some previous studies (Bolaño Losada et al., 2025; Dzurendova, Zimmermann, Tafintseva, Kohler, Ekeberg, et al., 2020). All the species of the three sets are listed in Table 1.

**Table 1.**
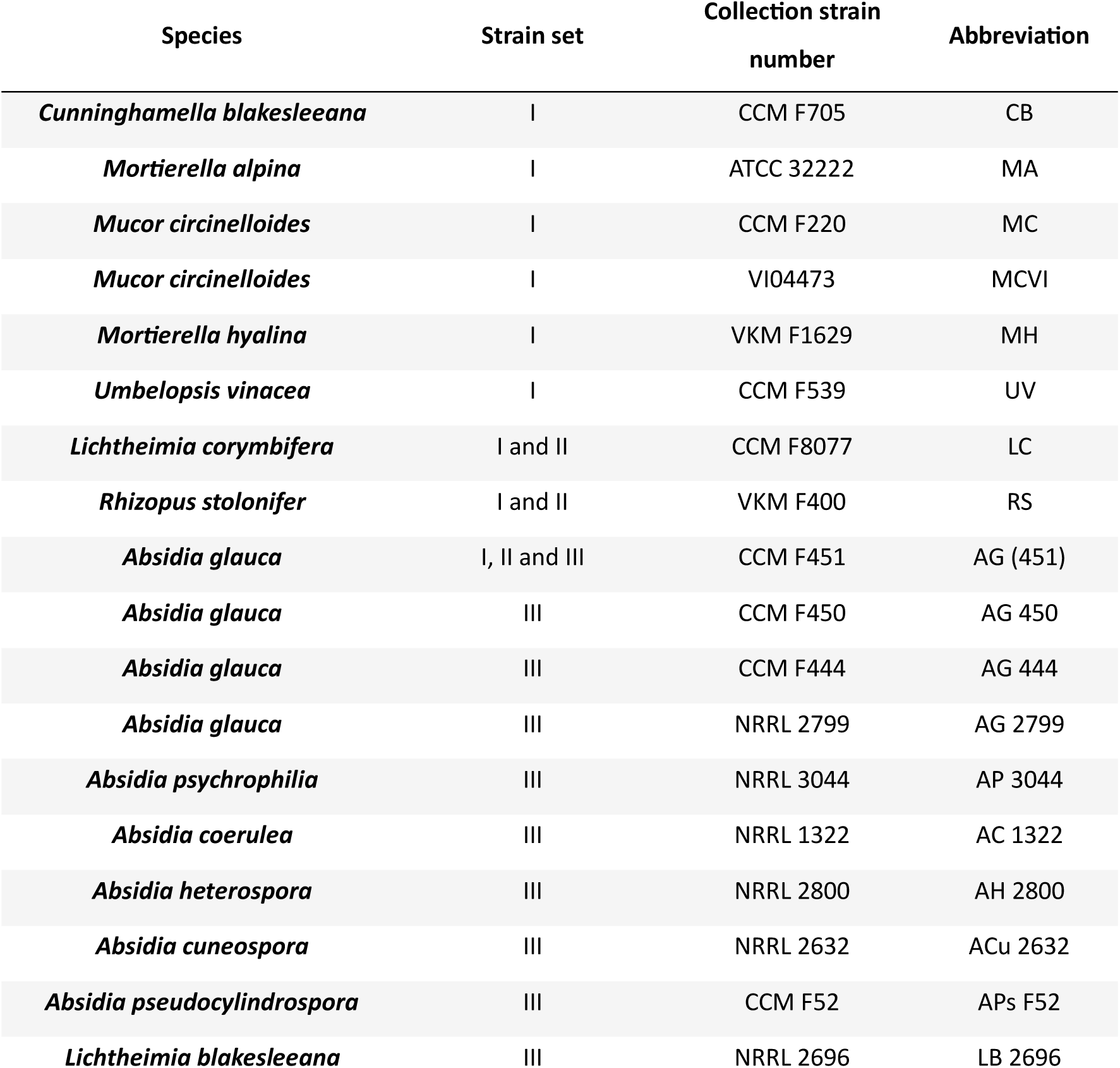
List of Mucoromycota strains with the corresponding culture collection codes and abbreviations used in this study.

**Table 2.** Biomass production, glucose consumption, pH at the end point of the cultivation, concentrations of lactic, fumaric, and oxalic acid from cultivation of *Absidia glauca* CCM F451 and *Rhizopus stolonifer* VKM F400 (Set II) in high and standard C/N medium for 5 days at 25 °C. -Δ glucose=-([glucose]_final_-[glucose]_initial_). Different letters indicate significant differences (*p* ≤ 0.05) between values of each variable and strain, while its absence indicates no significant differences.

| **Species** | **C/N** | **Biomass<br>(g/L)** | **-Δglucose<br>(g/L)** | **pH** | **Lactic acid<br>(g/L)** | **Succinic acid<br>(g/L)** | **Fumaric acid<br>(g/L)** | **Oxalic acid<br>(g/L)** | **Glycerol<br>(g/L)** | **Ethanol<br>(g/L)** |
|---|---|---|---|---|---|---|---|---|---|---|
| *A. glauca* | Standard | 4.47<sup>b</sup> ±0.08 | 18.3<sup>b</sup> ±0.00 | 3.57 ±0.10 | 0.05 ±0.00 | 0.00 ±0.00 | 0.00 ±0.00 | 2.00± 0.08 | 0.00 ±0.00 | 0.00 ±0.00 |
|  | High | 6.32<sup>a</sup> ±0.53 | 27.20<sup>a</sup> ±1.02 | 3.50 ±0.02 | 0.03 ±0.01 | 0.00 ±0.00 | 0.00 ±0.00 | 2.13 ±0.02 | 0.00 ±0.00 | 0.00 ±0.00 |
| *R. stolonifer* | Standard | 4.72<sup>b</sup> ±0.20 | 18.30<sup>b</sup> ±0.00 | 5.76<sup>a</sup> ±0.06 | 0.32 ±0.11 | 0.08<sup>b</sup> ±0.02 | 0.04<sup>b</sup> ±0.01 | 0.00± 0.00 | 0.80<sup>b</sup> ±0.54 | 0.26<sup>b</sup> ±0.21 |
|  | High | 6.60<sup>a</sup> ±1.05 | 27.36<sup>a</sup> ±0.49 | 4.48<sup>b</sup> ±0.07 | 0.17 ±0.00 | 0.22<sup>a</sup> ±0.02 | 0.18<sup>a</sup> ±0.03 | 0.00 ±0.00 | 8.62<sup>a</sup> ±0.18 | 2.05<sup>a</sup> ±0.06 |

It should be noted that the strain NRRL 2696 was originally deposited as *Absidia blakesleeana.* This designation is maintained in the NRRL culture collection catalog, while according to the last update of the strain’s history at the ATCC collection, it has been changed to *Lichtheimia blakesleeana*. The implications of this taxonomic discrepancy were addressed in the discussion section.

### Preparation of spore inoculum and stock cultures

As in our previous work (Bolaño Losada et al., 2025; Bolaño-Losada et al., 2025), malt extract agar (MEA) (Merck, Germany) was used as medium for mycelium and spore production. The fungal strains were incubated in this medium at 25 °C for 7-14 days. The spores were released from mycelium by dragging it with 10 mL of NaCl 0.9 % and a Digralsky spatula. The spores were stored in cryovials in with 33%_v/v_ glycerol at – 80 °C. As exception, *Mortierella alpina,* was regularly sub-cultured into MEA.

### Media composition

In our previous study (Bolaño-Losada et al., 2025), two media with different C/N ratio defined as “high” and “standard” were used. In the present study, we considered the introduction of a third medium with lower C/N ratio and defined as “low”. The nitrogen content of these media was kept constant, whereas the carbon content varied. The media composition was (g/L): glucose, 8.5 (for low C/N), 17 (for standard C/N) and 80 (for high C/N); peptone, 3; Na_2_HPO_4_, 2; KH_2_PO_4_, 7; MgSO_4_·7H_2_O, 1.5; CaCl_2_·2H_2_O, 0.1; FeCl_3_·6H_2_O 0.008; ZnSO_4_·7H_2_O, 0.001; CuSO_4_·5H_2_O, 0.0001; Co(NO_3_)_2_·H_2_O 0.0001; MnSO_4_·5H_2_O, 0.0001 and pH 6.2. The nitrogen and carbon content of peptone (WVR cat. no. J636-500g, Lot. 23l2556822) was 12% and 40%. The C/N ratio values for high, standard, and low medium were 89.0, 18.8, and 9.4, respectively. Considering also the importance of phosphate in the accumulation on some biomolecules, the P/N ratio value for all three media was 5.6, while C/P ratio values for high, standard, and low medium were 15.9, 3.4, and 1.7, respectively. It must be noted that both carbon and phosphorus content of peptone were not included in the C/N ratio calculation.

### Cultivation set-up to study carbon allocation strategies (strain Set I)

The cultivation of the fungal strains included in the strain Set I (Table 1) for the carbon allocation strategies experiment was performed as described in our previous study (Bolaño-Losada et al., 2025). Briefly, cultivations were performed using the Duetz-system. This consisted of a polypropylene square 96-deep well microtiter plates (MTPs) (Enzyscreen, The Netherlands, Cat #CR1496) with “sandwich covers” to ensure minimal evaporation. The plates were fixed in the shaker incubators with a Duetz clamping system. 800 µL of medium inoculated with 2 × 10^4^ spores/mL were added to each well. Separately, the inoculum of *M. alpina* was prepared by cutting four plugs of active MEA culture with the help of a cork borer no. 5. These agar plugs were smashed in a 15 mL Falcon tube, resuspended in 10 mL of 0.9 % NaCl, and used to inoculate 2-fold concentrated medium in a 1:1 (v/v) ratio. Each strain was cultivated in the different C/N media in triplicate.

The incubation was performed in a MaxQ^TM^ 4000 shaker (Thermo Scientific, USA) at 25 °C and 400 rpm for 5 days. After the incubation, the microplates were spun for 15 min at 4500 rpm and 4 °C for in a Hermle Z326K centrifuge (Hermle, Germany). Then, the supernatants were transferred and stored in deep-well 96-well plates, while biomass pellets underwent three washing cycles consisting of sequential resuspension in water, centrifugation, and removal of the supernatant. Both supernatant and biomass pellet samples were stored at −20 °C until further analysis.

### Cultivation to study production of short-chain carboxylic acids in *Absidia glauca* and *Rhizopus stolonifer* (strain Set II)

This study revealed a significant secretion of organic acids in the supernatants of *A. glauca* CCM F451 during cultivation of fungi from set I under low, standard and high C/N conditions. Considering the significant pH reductions in previous cultivations with this species (Bolaño Losada et al., 2025; Dzurendova, Zimmermann, Tafintseva, Kohler, Ekeberg, et al., 2020) and the absence of literature on its acid production, an experiment was conducted to identify and quantify the acid secreted by *A. glauca* CCM F451. In this experiment, *R. stolonifer* VKM F400 was included as a reference due to its well-documented ability to produce fumaric and lactic acid (Das et al., 2016).

To obtain enough amount of sample for the subsequent acid analysis, the cultivations were performed with standard and high C/N media using also a Duetz system with 24 extra high-deep well microtiter plates (Enzyscreen, The Netherlands, Cat #CR1424hd), low evaporation sandwich covers and clamping system. Each well contained with 7 mL of medium and inoculated with 2 × 10^4^ spores/mL. The plate was incubated for 5 days at 25 °C and 400 rpm. Each species was cultivated in triplicate for each C/N medium.

Following the incubation, the culture was collected in 15 mL Falcon tubes and centrifuged at 10 000 rpm and 10 min. Supernatants were separated and the biomass pellets washed by three successive cycles of resuspension in water – centrifugation – supernatant removal. The washed biomass was freeze-dried as described above. Both biomass and supernatants were stored at −20 °C until further analysis.

### Cultivation set up for production of short-chain carboxylic acids in *Absidia* species (strain Set III)

After identification and quantification of acids from *Absidia glauca* CCM F451, an additional experiment was conducted with *Absidia* strains. In this experiment, the strains from Set III were grown in high C/N medium using the same setting and cultivation conditions than in Set I. Briefly, Duetz system with 96-MTPs, low evaporation sandwich covers and a clamp system; 800 µL of medium inoculated with 2 × 10^4^ spores/mL; and incubation at 25 °C and 400 rpm for 5 days. The samples after fermentation were processed and stored as described previously for the strain Set I.

## Glucose determination

The glucose concentration in the supernatants from strain Set I, II, and III, was determined using two different methods, depending on the concentration range to ensure the highest precision. For the samples with glucose concentration exceeding 10 g/L, values were obtained by a predictive model method using the spectra acquired by Fourier transform infrared attenuated total reflectance (ATR-FTIR) spectroscopy and the partial least-square regression (PLSR) model developed in the work of Kosa et al. (Kosa et al., 2017) and Dzurendova et al. (Dzurendova et al., 2021). On the other hand, the samples with lower glucose concentrations were determined by the glucose oxidase/peroxidase method (GOPOD) using a commercial assay kit (Neogen, United States, cat. no. 700004297). The reagents were prepared according to the manufacturer’s instructions. The reaction was performed in triplicates in a 96-well plate and consisted of 10 µL of sample and 300 µL of GOPOD reagent. The mixture was incubated at 40 °C for 20 min and the absorbance was recorded at 510 nm. D-glucose was used as standard and distilled water with the GOPOD reagent was used as a blank. The evaporation rate during cultivation was considered to correct the glucose consumption.

### pH measurements

The pH from supernatants from experimental Set I, II and III, was determined using an Accumet XL600 pHmeter (Fisher Scientific, United States) equipped with a microtip pH electrode.

### Elemental analysis

Biomass samples from the strain Set II were subject for elemental analysis. This analysis was done with the aim to determine differences of phosphorus content between *A. glauca* (CCM F451), and *L. corymbifera* (CCM F8077). Alongside, *Rhizopus stolonifer* (VKM F400) biomass was also employed as a reference due to its high content of phosphorus. Additionally, magnesium, calcium, potassium and sodium were also determined. The sample was prepared using microwave assisted acid digestion prior to phosphorus determination using a MP-AES 4200 (Agilent Technologies, USA) microwave plasma atomic emission spectroscope.

### Ion-exclusion chromatography for extracellular biomolecules

Supernatants from the carbon allocation strategies experiment (strain Set I) were analyzed for the presence of extracellular glycerol and ethanol as in Losada et al. (Losada et al., 2025). In essence, the ultra-high performance liquid chromatography (UHPLC) equipment used consisted of a Dionex Ultimate 3000 (Dionex, Sunnyvale, CA, USA) with a Rezex ROA-Organic Acid H+ (8%) 300 × 7.8 mm column (Phenomenex, Torrance, CA, USA) coupled to a refractive index (RI) detector 101 (Shodex, Yokohama, Japan). The samples were eluted isocratically with 5 mM H_2_SO_4_ at a flow rate of 0.6 mL/min and 65 °C.

### High-throughput system Fourier transform infrared (HTS-FTIR) spectroscopy

Biomass pellet samples from the strain Set I experiment were processed directly in the 96-well microplate used for cultivation as in Bolaño-Losada et al. (Bolaño-Losada et al., 2025). In summary, mechanical disruption of washed mycelium was done by bead beating. Approximately 300 – 400 µL of distilled water and 250 mg of acid-washed beads (710–1180 μm diameter, Merck, Germany) were added to each well with the biomass pellet. The microplate was closed with a silicon mat lid and processed by bead beating using a Bead Ruptor 96 (Omni International, United States). The disruption procedure consisted of a total of three runs of 3.5 min at 25 Hz, each one separated by a resting time of 30 s.

The HTS-FTIR spectra acquisition was performed as in our previous study (Bolaño-Losada et al., 2025). In summary, a high-throughput screening module (HTS-XT) coupled to a Vertex70-FTIR spectrometer (both Bruker Optik GmbH, Germany) with a globar mid-IR source and a deuterated L-alanine doped triglycene sulfate (DLaTGS) detector was used. The spectra acquisition was done in transmission mode using a IR-transparent 384-well silicon microplate (Bruker Optik GmbH, Germany). The samples were prepared by dispensing 8 µL of homogenized biomass in each well of the plate and leaving it dry at room temperature. Each sample was dispensed in three different wells of the plate to acquire three technically independent measurements. The acquisition was performed in the spectral range of 4000 to 400 cm^-1^, with spectral resolution of 6 cm^-1^, digital spacing of 1.928 cm^-1^, 6 mm aperture, 64 scans and 3-Term Blackman-Harris apodization. Sample spectra were recorded using the spectrum of an empty well in the IR-transparent microplate as reference background. The OPUS 8.2 software (Bruker Optik GmbH, Ettlingen, Germany) was used for data acquisition and instrument control.

In total, 373 spectra were acquired: 238 spectra from 81 samples from Set I (Table 1), 45 spectra from 15 samples from Set II (*A. glauca* CCM F451 and *R. stolonifer* VKM F400 grown in high and standard C/N media, and *L. corymbifera* CCM F8077 grown only in high C/N medium), and 90 spectra from 30 samples from strain Set III (Table 1). Some technical replicates from samples of Set I were not considered due to inadequate quality (e.g. low absorbance values in characteristic regions).

### Fourier transform infrared attenuated total reflectance (FTIR-ATR) spectroscopy

The FTIR reflectance spectra (FTIR-ATR) were measured manually as in our previous works (Bolaño-Losada et al., 2025). Measurements were performed in a Vertex 70 FTIR spectrometer equipped with a single reflectance-attenuated total-reflectance (SR-ATR) High-Temperature Golden Gate ATR Mk II accessory (Specac, United Kingdom) with a horizontal SR-ATR diamond prism with a 45° angle of incidence. The spectra were acquired over the range of 4000 to 600 cm^-1^, with spectral resolution of 4 cm^-1^, digital spacing of 1.928 cm^-1^ and 32 scans, using Blackman-Harris 3-Term apodization function. The measurement of supernatants from the cultivations of Set I, II, and III required 10 µL for each spectra acquisition. Prior to each measurement, a background (reference) spectrum was recorded using the sample-free setup. Three technically independent acquisitions were performed for each sample. The OPUS 8.2 software (Bruker Optik GmbH, Germany) was used for data acquisition and instrument control.

A total of 310 spectra were acquired: 184 spectra from 81 samples (only high and std C/N conditions) from Set I strains (Table 1), 36 spectra from 12 samples from Set II (only *A. glauca* CCM F451 and *R. stolonifer* VKM F400 in high C/N medium), and 90 spectra from 30 samples from Set III (Table 1).

### Spectral preprocessing and data analysis

To analyze the carbon allocation toward different biomolecules from the strain Set I, a ratiometric analysis was carried out on the acquired FTIR spectra. The spectra were preprocessed with data truncation from 3650 to 700 cm^-1^ and rubber band baseline correction. Each metabolite-ratio was determined using the maximum intensity between 1655 and 1620 cm^-1^ of the Amide I (C=O stretching) band as the denominator, while the numerator corresponded to a particular metabolite-related peak. For the lipid-to-protein ratio (LP), the maximum intensity between 1750 and 1735 cm⁻¹, which corresponds to C=O stretching band of acylglycerols, was used. For the phosphate-to-protein ratio (PP) the maximum intensity between 1280 and 1260 cm⁻¹ of the phosphate-related P=O stretching band was selected. Lastly, for the carbohydrate-to-protein ratio (CP) the maximum intensity between 1085 and 1070 cm⁻¹ of the C-O stretching band assigned to saccharides was used.

Separately, for explanatory and comparative purposes, the spectra of *M. circinelloides* CCM F220 (MC) were preprocessed by data truncation (3650 to 700 cm^-1^), baseline rubber band correction, area normalization at 1645-1620 cm^-1^ (Amide I) and averaged (three acquisitions from three biologically independent samples) for display in Fig. 1.

**Figure 1.**
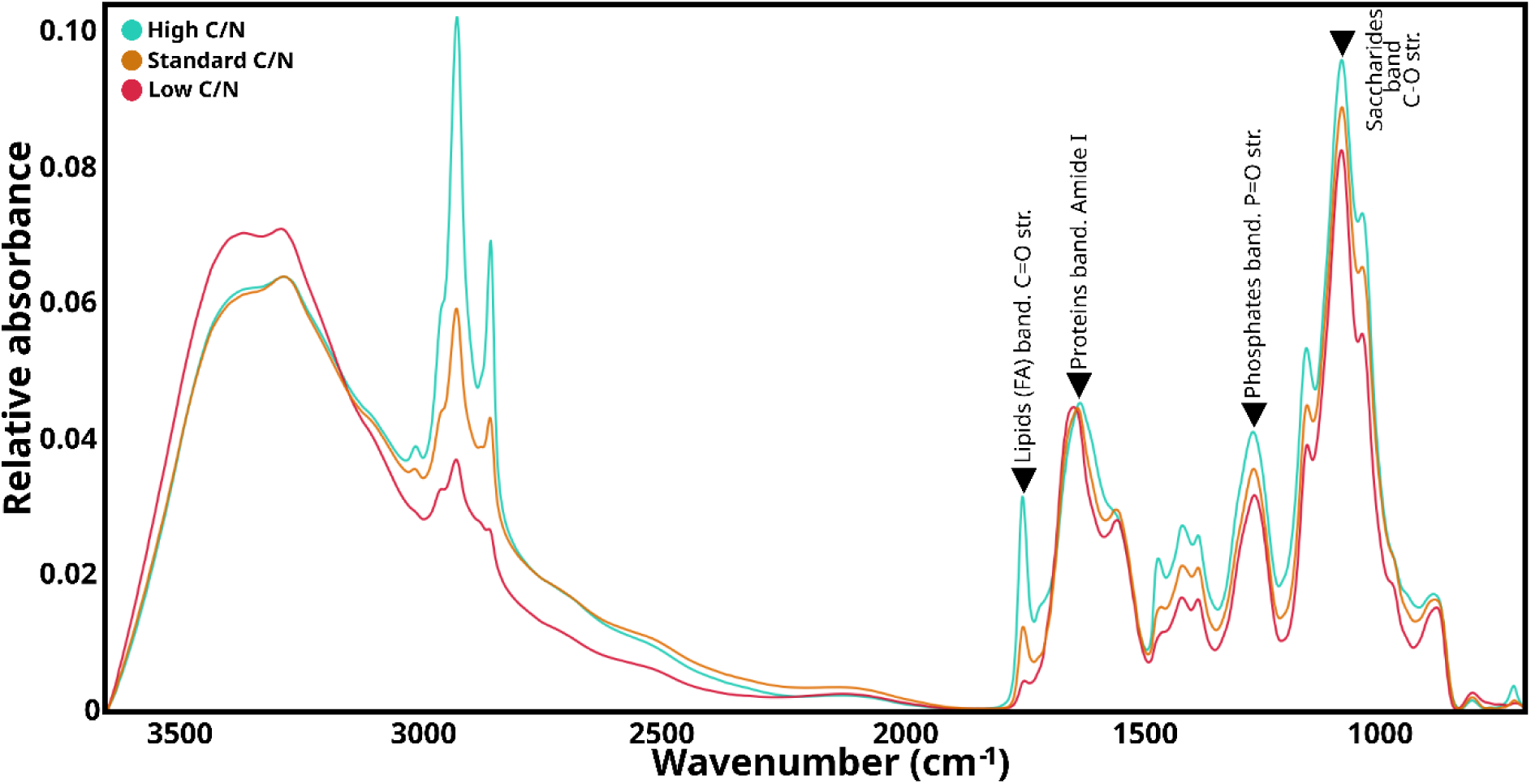
HTS-FTIR spectra from *M. circinelloides* (MC) grown in low, standard and high C/N media with the most characteristic peaks of each biomass metabolite (proteins, lipids, phosphates and saccharides) marked by arrows and labels. FA: fatty acids; str.: stretching.

The HTS-FTIR spectra from the comparative experiment of acid production between *R. stolonifer* and *A. glauca* was preprocessed by Savitzky-Golay filter (polynomial order 2, derivative order 0), data truncation between 3650 to 700 cm^-1^ and area normalization by Amide I band between 1655 to 1625 cm^-1^.

The HTS-FTIR spectra from *Absidia* species biomass (Set III) were preprocessed by data truncation from 3650 to 700 cm^-1^ and extended multiplicative signal correction (EMSC) (Guo et al., 2018; Kohler et al., 2020). For an easier identification and interpretation of differences between species and strains a principal component analysis (PCA) was applied to the spectra.

The FTIR-ATR spectra from supernatants were used for glucose determination by applying the expanded partial least square regression (PLSR) model from our previous studies was used (Dzurendova et al., 2021; Kosa et al., 2017). The model was based on FTIR spectra of supernatant samples (Dzurendova et al., 2021; Kosa et al., 2017), and a series of reference glucose and phosphate salt solutions (Dzurendova et al., 2025). Prior application of the PLSR model, the supernatant ATR-FTIR spectra were preprocessed by truncating the data from 700 to 1900 cm⁻¹, followed by baseline offset correction and peak normalization at 1635 cm^-1^ (i.e. peak corresponding to the bending vibration of water).

The preprocessing and visualization of spectra was performed using Orange Data Mining version 3.33.0 (University of Ljubljana, Slovenia) (Demšar et al., 2013; Toplak et al., 2017). The ratiometric analysis of biomolecules, glucose, biomass, and pH were visualized with Matplotlib. The glucose consumption, pH decreases, and oxalic acid values were analyzed and visualized with Excel.

### Gas chromatography-mass spectrometry (GC-MS) and ion-exchange chromatography for short-chain carboxylic acids analysis

Acidification of medium was related to secretion of organic acids. Several species of Mucoromycota used in this study lacks evidence of which short-chain carboxylic acids produce. This motivated their identification by GC-MS.

The extraction of carboxylic acids from the supernatants was performed as follows: in a Pyrex glass tube, 500 µL of supernatant was acidified with 100 µL of concentrated HCl, followed by the addition of 1 mL of diethyl ether, vortexing the obtained mixture for 2 min, and transferring the organic phase into another tube using a glass Pasteur pipette. 3 mg of sodium sulfate was added to the recovered organic phase to remove traces of water before filtration with a 0.22 µm PTFE hydrophobic syringe filter. The derivatization of the carboxylic acids was done as follows: 150 µL of N,O-Bis(trimethylsilyl)trifluoroacetamide (BSTFA) (Merck, Germany, Cat # 15222-1ML) and 150 µL of pyridine were added to 600 µL of the diethyl ether extract, followed by heating at 65 °C for 30 min.

For the GC-MS determination, an Agilent 8890 gas chromatograph system coupled with an Agilent 5977 MSD quadrupole mass spectrometer detector and a J&W DB-5HT column (Agilent, Germany) with 15 m × 320 µm × 0.1 µm dimensions (length × inner diameter × film thickness). The injection volume was 2 µL. Separation was carried out using a constant flow of H_2_ at 2.69 mL/min and temperature ramp as follows: start at 35 °C, increasing at 10 °C/min until 150 °C, then increasing at 50 °C/min until 380 °C and holding for 2 min. The total run time was 19.1 min. The detection started after 0.9 min, using electron ionization at 70 eV and 230 °C, with the quad temperature set at 150 °C and the acquisition with scan mode set at N=2 speed, range detection from 50 to 400 *m/z*, and threshold at 20. The different C/N ratio media and distilled water were analyzed in the same way and used as negative controls while a standard solution of oxalic acid was used to corroborate its retention time and mass spectrum match with the NIST standard reference database Main EI MS library (mainlib).

The quantification of acids was carried out by ion-exchange chromatography using a Dionex ICS-3000 system (Dionex Corporation, United States) equipped with a Dionex^TM^ IonPAC^TM^ AS11-HC-4µm IC column (4 × 250 mm) and with a conductivity detector. A Dionex^TM^ ASRS 4 mm suppressor was used, operating at 99 mA. Control and analysis were performed with Chromeleon 6.80 software. The sample was injected at a volume of 20 µL and eluted by gradient elution using two solutions: A) distilled water HPLC grade and B) 50 mM NaOH. Elution was performed as follows: from 0 – 5 min at 98:2 (A:B), 5 – 15 min at 70:30 (A:B), 15 – 20 min at 40:60 (A:B), 20 – 20.1 min at 98:2 (A:B), 20.1 – 35 min 98:2 (A:B).

## Statistical analysis

One-way ANOVA test was performed to determine differences among C/N conditions for each variable studied in each strain from Set I and Set II. Statistical significance was established at *p* ≤ 0.05. Subsequently, a *post hoc* pairwise comparison analysis was performed using Tukey’s Honest Significant Difference test, and the results were reported using Compact Letter Display. The statistic test results are provided in Supplementary file 1. All these tests were conducted using Python (version 3.12.1) using Jupyter Notebook.

## 3. Results and discussion Carbon allocation strategies

In our previous work, the biochemical changes induced by varying C/N ratios and H_2_O_2_ in several species of Mucoromycota fungi were discussed (Bolaño-Losada et al., 2025). The most notable changes observed in each strain were primarily related to variations of C/N ratio in the media, revealing species-specific carbon allocation strategies towards different intracellular biomolecules (Bolaño-Losada et al., 2025). Therefore, carbon allocation strategies at “low”, “standard” and “high” C/N conditions were further studied in detail in this work.

Initially, glucose consumption and biomass production were determined to investigate the growth yield of the strains under different medium C/N ratios. Then, to assess carbon allocation towards intracellular biomolecules, a ratiometric analysis of the FTIR spectra from fungal biomass was applied. FTIR spectroscopy provides comprehensive insight into fungal chemistry by simultaneously capturing broad chemical profiles of biomass via the detection of numerous functional groups (Dzurendova et al., 2025). To give an example, Fig. 1 shows the spectra of *M. circinelloides* (MC) biomass produced at the three different C/N ratios, highlighting the most characteristics peaks of the different types of biomolecules, specifically, proteins, lipids, phosphates and saccharides. In addition, a more detailed assignation of the biomass spectra can be found in our previous work (Bolaño-Losada et al., 2025; Dzurendova, Zimmermann, Kohler, et al., 2020; Kosa et al., 2017), while the averaged spectra for each strain and C/N condition are presented in Supplementary file 2. By applying the ratiometric analysis of FTIR spectra, we were able to estimate the accumulation of lipids, phosphates, and carbohydrates in the samples (Dzurendova et al., 2021). Specifically, due to the nitrogen content being equal for the three media, the Amide I band, representing proteins and chitin, was considered a stable reference across species and conditions, and thus the absorbance of this band was used as a proxy for total protein content. Accordingly, ratios of lipid-to-protein, phosphate-to-protein, and carbohydrate-to-protein were calculated using this band and C=O str. band for lipids, P=O str. band for phosphates and C-O str. band for carbohydrates to account for changes in the relative content of the respective biomolecules. Finally, the carbon allocation towards extracellular biomolecules was assessed by determination of media acidification to reflect acid production, ethanol and glycerol. To sum up, Fig. 2 shows the biomass growth and intra- and extracellular biomolecules production across the different Mucoromycota species revealing their carbon allocation strategies.

**Figure 2.**
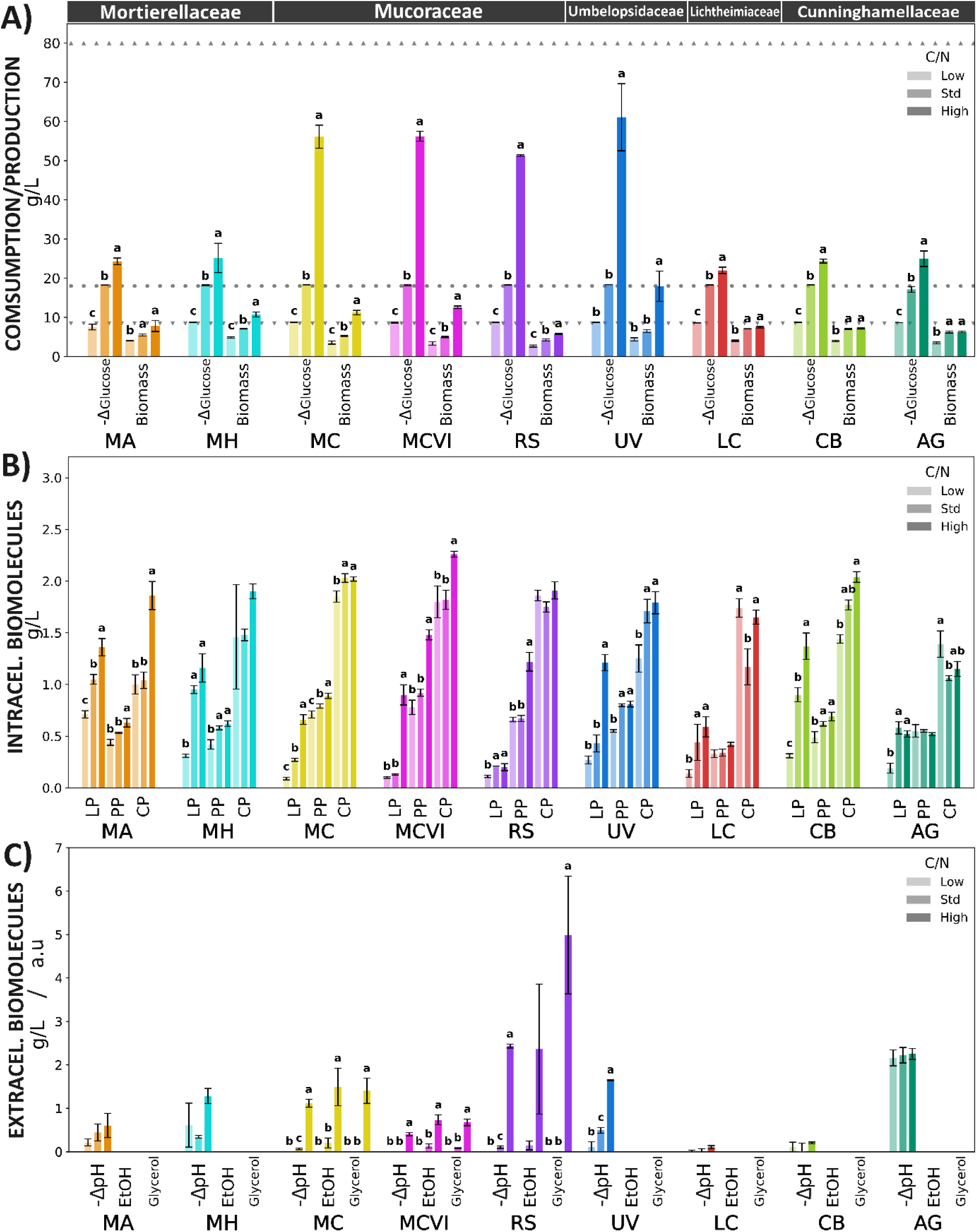
Results from carbon allocation experiment (strain Set I) with *M. alpina* (MA), *M. hyalina* (MH), *M. circinelloides* (MC), *M. circinelloides* (MCVI), *R. stolonifer* (RS) and *U. vinacea* (UV) *C. blakesleeana* (CB), *L. corymbifera* (LC), and *A. glauca* (AG) grown at low, standard and high C/N media. A) Glucose consumption (-Δglucose = -(glucose_final_-glucose_initial_)) and production of biomass for the nine Mucoromycota. Background dashed lines indicate the glucose concentration of each C/N medium (▾ low=8.5 g/L, ● standard = 17 g/L, and ▴ high = 80 g/L). B) Intracellular metabolite production at low, standard and high C/N medium reflected by the spectra ratios lipid-to-protein (LP), phosphate-to-protein (PP) and carbohydrate-to-protein (CP). C) Extracellular metabolite production at low, standard and high C/N, reflected by medium pH acidification (-ΔpH = -(pH_final_ -pH_final_)) as indicator of organic acids, ethanol and glycerol. Error bars represent the standard deviation of three biologically independent replicates. Different letters indicate significant differences (*p* ≤ 0.05) between values of each variable and strain, while its absence indicates no significant differences.

Before the discussion of the results, it is worthwhile to briefly mention the biological roles of these biomolecules. The accumulation of acylglycerols in lipid bodies serves as a reservoir for carbon and energy (Leman, 2009; Thevelein, 1984). Polyphosphates have location-based functions such as energy storage molecules, countering chitosan ions or pH regulation (Achbergerová & Nahálka, 2011; Barbosa De Lima et al., 2003; Do Nascimento Shari’a et al., 2002; Dzurendová et al., 2026; Werner et al., 2007). Saccharides are present either as structural or storage components. Increase on the fungal cell wall thickness, which includes several types of saccharides, is often a response to acid stress; meanwhile, intracellular saccharides, such as glucose, trehalose, and glycogen, are related to carbon and energy storage, among other functions (Crognale et al., 2022; Dzurendova, Zimmermann, Kohler, et al., 2020; Elbein, 2009). The extracellular presence of ethanol is commonly a result of anaerobic respiration activity, although several Mucoromycota species are Crabtree positive, meaning that they can also catabolize glucose through ethanolic fermentation even in presence of oxygen when glucose is available in high concentrations. As previously commented, glycerol is also secreted as byproduct of ethanol production (Safaei et al., 2016). The secretion of short-chain carboxylic acids facilitates metal uptake by solubilization and chelation, promotes the breakdown of substrates to facilitate digestibility (e.g., plant materials), or acts as a defense mechanism against competitors (Jiao et al., 2022; Liaud et al., 2014). The synthesis of all these biomolecules is ultimately supported by glucose consumption, and a preference of producing one over another may reflect differences in species’ lifestyles and phylogenetic relationships.

A first general indication of differences between carbon allocation to intracellular and extracellular biomolecules can be withdrawn from values of biomass yield. Higher values can indicate more preference for intracellular than extracellular biomolecules between species. For example, *Mortierella alpina* species in comparison with *Rhizopus stolonifer* (Fig. 2 and Supplementary file 4, Fig. S1). For each species, biomass yield decreased with higher C/N (meaning less carbon limitation). This trend could be attributed to a combination of factors, including investment in more energy-demanding biomolecules (e.g. lipids in comparison to saccharides), increases in extracellular metabolite production, and prolonged maintenance of metabolic active cells.

Hereafter, these observations will be addressed separately according to the taxonomic level of family.

### Mortierellaceae

*M. alpina* and *M. hyalina* exhibited a moderate maximum of glucose consumption around 25 g/L (Fig. 2a) with a progressive increase in biomass as C/N increases. For both *Mortierella*, the excess of carbon was primarily diverted towards the accumulation of lipids and saccharides (Fig. 2b). In our previous work we indicated that this accumulation of saccharides was likely intracellular (e.g., glycogen, trehalose, etc.) rather than in cell wall polysaccharides (Bolaño-Losada et al., 2025).

While lipid accumulation in this group of fungi has been widely studied, saccharide accumulation received less attention (Chang et al., 2023; Chen et al., 2021). *Mortierella* species commonly display cold adaptability and are frequently isolated from alpine, Antarctic, and Arctic environments, whereas other Mucoromycota representatives are absent (dos Santos et al., 2020; Gonçalves et al., 2012; Perini et al., 2019; Schmidt et al., 2008). The observed accumulation of saccharides may be related not only to a carbon storage strategy but also to physiological adaptation to cold environments, where saccharides like trehalose or polyols like glycerol play a role as cryoprotectants. Indeed, *Mortierella* species possess temperature priming for stress tolerance (Andrade-Linares et al., 2016) and are also considered snow-mold species (Telagathoti et al., 2021).

The term snow-molds is a general term for fungi that thrives beneath snow cover under conditions of darkness, high moisture and low temperatures and, generally, infecting plants. However, *Mortierella* appears to deviate from the typical phytopathogenic assumption of snow molds, because: i) *Mortierella* species do not express relevant plant cell wall degrading enzymes even in presence of inducers such as cellulose or cellobiose, and are not efficient degrading plant litter materials (Bolaño Losada et al., 2025; Leifheit et al., 2024; Schmidt et al., 2008), and ii) they are frequently reported as positive plant-interacting fungal species, promoting plant growth and protecting them from nematode pest threats (Bonfante & Venice, 2020; Büttner et al., 2024).

Therefore, rather than actively infecting and consuming plant material, as other snow-fungi do, *Mortierella* species might remain cryoprotected, relying on internal reserves until winter ends and rapidly benefit from the sugars released by the thaw of the plant material (Matsumotoo Naoyukii & Hsiang Tom, 2016). The lipid bodies rich in highly unsaturated lipids typical of the *Mortierella* species, might remain fluid and easily available to use at low temperatures conditions. These characteristics may help maximize survival during winter while outcompeting non-cold-adapted species. The actual knowledge on these species reinforces this snow-mold “opportunistic” mechanism suggested by Schmidth et al. (Schmidt et al., 2008).

In addition, ice nucleation activity was found to be widespread in *M. alpina*, a mechanism that could function as freeze protection mechanism, cause plant tissue injury, or assist in soil and rock cleavage for mineral release (Fröhlich-Nowoisky et al., 2015). However, more studies are required to further support these hypotheses. A more detailed study on the intracellular saccharide profile, such as that one of Danilova et al. (Danilova et al., 2025) with *Mucor psychrophilus* or Chen et al. (Chen et al., 2021) with *M. alpina*, could help to provide a better understanding of this observed physiology.

Finally, the excess of glucose was also diverted to the production of organic acids, reflected by the media acidification, which increased with the increase of C/N ratio (Fig. 2c). The GC-MS analysis identified lactic acid as the major organic acid produced (Supplementary file 3), which was previously reported for other *Mortierella* species (Uehling et al., 2019). The secretion of lactic acid, among other acids, was suggested to have a role in metabolite exchange with bacteria (Uehling et al., 2019). This could be an explanation for the case of *M. alpina* and *M. hyalina* observed in this study, whose endo- or epi-hyphal bacterial association are common (Telagathoti et al., 2021; Uehling et al., 2019).

### Mucoraceae

*M. circinelloides* (strains MCVI and MC) showed a high maximum glucose uptake (≈ 56 g/L) (Fig. 2a). In general, both *M. circinelloides* strains (MC and MCVI) primarily directed carbon to the production of biomass predominantly enriched with lipids and moderately with phosphates (Fig. 2b). According to our previous works, most of these phosphates are in form of polyphosphates (Dzurendova et al., 2021). It is worth noting that polyphosphate accumulation occurs from the beginning of the fermentation, which seems to be a process independent from nitrogen starvation in contrast to lipid accumulation (Barbosa De Lima et al., 2003; Dzurendová et al., 2026).

The exceeding carbon was also directed towards extracellular biomolecules such as ethanol, glycerol, and organic acids (lactic acid, Supplementary file 3), particularly under high C/N. Additionally, small but consistent differences were observed between these two strains. *M. circinelloides* (MC) allocated carbon more into extracellular biomolecules (ethanol, glycerol, organic acids) to the detriment of biomass generation, which was also slightly lower in lipid and polyphosphate content than in the *M. circinelloides* (MCVI) strain. In addition, a sharper increase in saccharide content was noted for the strain MCVI compared to MC under high C/N conditions. The strain MCVI was previously reported to overproduce more cell wall than the MC strain in response to acidic stress, which increased with higher C/N conditions (Dzurendová et al., 2021; Dzurendova, Zimmermann, Kohler, et al., 2020). It is noteworthy that this species is not commonly associated with large secretion of acids, but rather with moderate to low levels (Millati et al., 2005). Therefore, additional contributing factors, such as CO_2_-driven acidification from aerobic glucose respiration, or extensive uptake of phosphates reducing the medium’s buffering capacity should be considered as an explanation to the pH drop.

On the other hand, *R. stolonifer* exhibited a completely different carbon allocation strategy. Under high C/N conditions, a large amount of glucose consumption (≈ 51 g/L) was observed, while biomass was only moderately produced and enriched mainly in polyphosphates and to lesser extent in lipids (Fig. 2a and 2b). In contrast, a large production of extracellular biomolecules such as ethanol, glycerol and organic acids was observed, specifically in high C/N conditions (Fig. 2c). The production of acids is often promoted by excess of carbon in nitrogen limiting conditions (Wierzchowska et al., 2021). The secretion of large amounts of acids in these conditions is well known in the genus *Rhizopus*, being primarily fumaric and lactic acid, but also succinic in smaller amounts (Martin-Dominguez et al., 2018), in agreement with our study (Supplementary file 3). This metabolic strategy aligns well with the lifestyle of *R. stolonifer* as food spoilage fungus, especially in vegetables and fruits (Elisa et al., 2025). The secretion of organic acids, expression of vegetable cell wall-degrading enzymes, and its rapid hyphal extension rates supports a fast colonization, maceration and nutrient acquisition from the substrate, outcompeting other fungi (Bolaño Losada et al., 2025; Petrasch et al., 2019; Vellozo-Echevarría et al., 2024).

Unlike other species, *R. stolonifer* seems to not only prioritize lipid accumulation as form of energy storage, but polyphosphates too. Since lipid bodies can achieve a significant diameter, occupying nearly the entire lumen of hyphal lumen (Dzurendova, Zimmermann, Kohler, et al., 2020), they may hinder the mobilization of growth machinery needed for rapid hyphal extension and timely enzyme production during infection. In contrast, polyphosphates may serve as a more efficient energy reserve, although not carbon, since they are accumulated in form of granules much smaller than typical lipid bodies. In addition, polyphosphates are accumulated not only intracellularly but also in the cell wall, where it may have a protective function against the high amount of acid produced. Polyphosphates could act as counterions to stabilize positively charged chitosan, preventing gelation (Werner et al., 2007), aid in the retention of divalent cations such as Mg^+2^ and Ca^+2^, or sequester protons to prevent intracellular acidification. Further studies on the location and roles of polyphosphates in this fungus are needed to provide deeper insights. Thus, these observations reflect also its potential as a phosphorus-removing microorganism in wastewater treatment, while simultaneously producing industrially relevant organic acids, e.g. fumaric acid. To give an instance, some works have reported concentrations of phosphorus in *Rhizopus* species between 3-9% over dry weight (Christy et al., 2026; Dzurendova, Zimmermann, Kohler, et al., 2020).

Finaly, in this study, a general observation for the Mucoraceae family species was the production of ethanol, in aerobic conditions and high glucose concentrations (high C/N). This phenomenon, known as the Crabtree effect, has been reported for other species of Mucoraceae family such as *Rhizopus oryzae*, *Mucor indicus*, *Mucor hiemalis*, *Mucor corticolous*, while not in species from different families, such as *Rhizomucor* (Rhizomucoraceae) (Lübbehüsen et al., 2004; Millati et al., 2005), consistent with the results of this study. We noted also the production of glycerol, which was previously commented to be a byproduct during ethanol fermentation. Mucoraceae species are fast growing and assimilating simple sugars (Bolaño Losada et al., 2025). The Crabtree effect generates energy in parallel to the aerobic respiration, which might boost the hyphal extension rate facilitating the competition for substrate colonization, even under limited oxygen availability.

Thus, in *M. circinelloides*, the alcohol dehydrogenase 1 (ADH1) not only participates in the Crabtree effect but also in the consumption of ethanol once other simple sugars are depleted (Lübbehüsen et al., 2004; McIntyre et al., 2002; Millati et al., 2005). Therefore, ethanol is not only produced for extra energy acquisition but can also be reutilized later as a carbon source. This trait, combined with others discussed in our previous works, may contribute to the success of these species in food spoilage (Bolaño Losada et al., 2025; Elisa et al., 2025). Separately, the knock-out of alcohol dehydrogenase could hypothetically redirect carbon to other biomolecules, such as lipids, although a recent study in *M. circinelloides* suggested that this enzyme also impacts the upregulation of lipogenesis genes and lipid production (Shah et al., 2022). Nevertheless, more studies will be required to fully understand these dynamics.

### Umbelopsidaceae

*U. vinacea* species showed the highest glucose consumption (≈65 g/L) and biomass production, channeling carbon predominantly into lipids and moderately into polyphosphates (Fig. 2a and 2b). A high capacity for lipogenesis is common in *Umbelopsis*, reaching astonishing values in the order of 75-80% over total dry weight (Chatzifragkou et al., 2010; Dzurendová et al., 2021; Langseter et al., 2021). The carbon allocation strategy of *U. vinacea* resembles that of *M. circinelloides*, while without ethanol and glycerol production. The Umbelopsidaceae family is an early divergent group and far from the Mucoraceae family, therefore this similarity can be purely convergent (Zhao et al., 2023). Regarding medium acidification, a significant pH decrease was observed mainly in high C/N conditions, in agreement with the production of lactic acid (Fig. 2c and Supplementary file 3). Additional factors such as CO_2_ production and removal of phosphate should also be considered in relation to acidification, as discussed previously for Mucoraceae.

*Umbelopsis* species are considered cosmopolitan soil saprotroph with versatile enzymatic machinery, especially for carbohydrates (Muszewska et al., 2021), but also some species have been reported to possess endophytic traits (Molina et al., 2023; Qin et al., 2018). In addition, lactic acid has been suggested to mediate interactions with endobacteria in some species of *Umbelopsis* (Uehling et al., 2019). The accumulation of polyphosphate in this species can be potentially linked to the interaction with plants by exchanging phosphate for other molecules. Meanwhile, the massive lipid accumulation might have a classical role as energy and carbon reserve supporting growth and stress resilience.

### Cunninghamellaceae and Lichtheimiaceae

*A. glauca* and *C. blakesleeana*, from the family Cunnighamellaceae, and *L. corymbifera*, from the closely related family Lichtheimiaceae, showed maximum glucose consumption values around 22-25 g/L (Fig. 2a). The biggest change in biomass concentration was found from low to standard C/N, while the values were similar, from standard to high C/N. In these species, the excess of carbon invested in intracellular biomolecules was mainly directed towards lipids. Of note, *A. glauca* biomass productivity was slightly lower than *C. blakesleeana* and *L. corymbifera* in all C/N conditions (Fig. 2a). This can be explained by the allocation of carbon towards extracellular biomolecules, which in this case was the secretion of organic acids in all the C/N conditions, reflected by a pronounced acidification (>2-ΔpH) (Fig. 2c). The secretion of organic acids by *A. glauca* seems to be constitutive and independent of typical triggering factors such as carbon excess under nitrogen-limiting conditions. This outstanding observation and the lack of reports in the literature motivated the series of additional experiments discussed in the following sections. In brief, no short-chain carboxylic acids were detected for *L. corymbifera*, lactic acid was detected for *C. blakesleeana,* and lactic acid and oxalic acid were detected for *A. glauca* (Supplementary file 3).

Small differences were also notice between the three species. To give an instance, phosphate-to-protein ratios (PP) were higher in Cunninghamellaceae (*C. blakesleeana* and *A. glauca*) than in Lichtheimiaceae (*L. corymbifera*) indicating differences in phosphorus accumulation capacities between these two families.

### Acid production in Rhizopus stolonifer and Absidia glauca

As discussed in the previous section, *A. glauca* and *R. stolonifer* secreted organic acids that significantly lowered the medium’s pH (3.5-4.5). The acid production by both species was determined in a parallel fermentation experiment. Contextualizing, more than one hundred years ago, in 1911, the production of fumaric acid was reported for *R. stolonifer* (originally *Rhizopus nigricans*), and some years later the production of lactic, succinic, and other acids (Takahashi & Sakaguchi, 1927). In contrast, no reports in the literature have described the secretion of any organic acid in *A. glauca*.

The GC-MS analysis of supernatants from *R. stolonifer* verified the presence of fumaric, succinic, lactic acid, and glycerol under both standard and high C/N conditions (Supplementary file 3). Lactic acid and fumaric acid share pyruvate as a common precursor but diverge along distinct metabolic pathways, whereas succinate is the immediate precursor of fumaric acid in the Krebs cycle (Longacre et al., 1997). On the other hand, a significant presence of oxalic acid, and minor amounts of lactic acid, was detected in the standard and high C/N media for *A. glauca* (Fig. 3). Retention time and mass spectrum values coincided with the derivatized oxalic acid standard run in the same conditions (Supplementary file 3).

**Figure 3.**
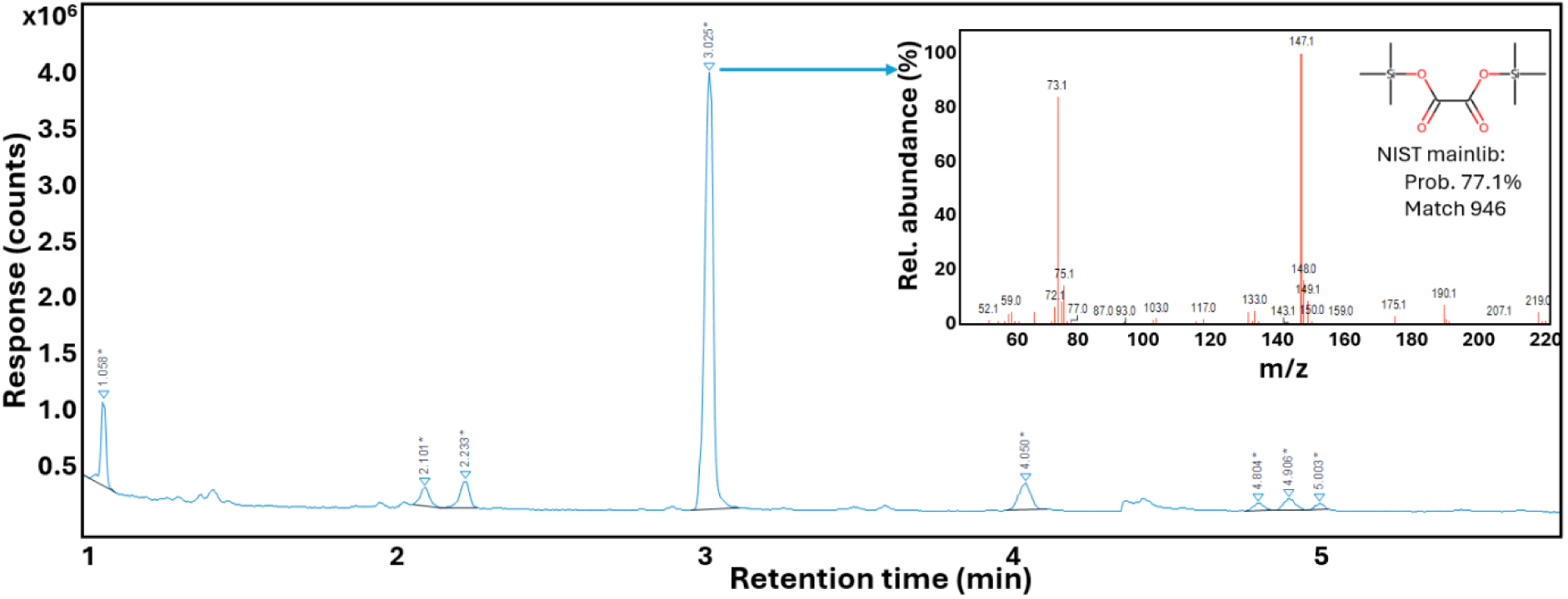
GC-MS chromatogram from BSTFA-derivatized ethyl ether extract of supernatants from *A. glauca* in high C/N medium. The oxalic acid peak is displayed at a retention time of 3.025 min with the associated mass spectrum matching oxalic acid-2TMS derivative from the NIST *mainlib* library with a 77.1% probability.

Oxalic acid is a dicarboxylic acid with stronger acidity (p*K*_a1_ = 1.27 and p*K*_a2_ = 4.27) than fumaric acid (p*K*_a1_ = 3.03, p*K*_a2_ = 4.44). In Mucoromycota, some species, such as *Mucor mucedo*, *Pilobus sp*., or *Gilbertella persicaria* are known to form calcium oxalate crystals located in the cell wall of aerial reproductive structures (Birkby & Preece, 1988; Urbanus et al., 1978; Whitney & Arnott, 1986). Nonetheless, the presence of oxalate crystals does not necessarily imply oxalic acid secretion. Indeed, *Mucor mucedo* forms calcium oxalate crystals, but it was tested negative for secretion of oxalic acid in liquid medium fermentation (Wehmer, 1891a). In this study, we demonstrated, for the first time, that the vegetative hyphae of *A. glauca* secretes significant amounts of oxalic acid during submerged fermentation.

The quantitative HPLC analysis revealed that *A. glauca* sharply acidified the medium (pH ≈ 3.5) due to oxalic acid production at concentrations around 2 g/L, with minor amounts of lactic acid (0.05–0.03 g/L), in both standard and high C/N media. In contrast, *R. stolonifer* produced lower amounts of acids, primarily lactic and fumaric acid, resulting in a more modest pH decrease (4.48 to 5.76). Under standard C/N conditions, lactic acid concentration was higher than fumaric acid (0.32 vs. 0.04 g/L), whereas under high C/N conditions, this trend reversed (0.18 vs. 0.17 g/L), accompanied by an increase in succinic acid production (0.08 vs. 0.22 g/L). This shift suggests a metabolic preference for the fumaric acid pathway over lactic acid under high C/N conditions.

On a different note, differences between cultivations in 24-well plates and 96-well were observed (“Carbon allocation strategies” section). These differences can be associated with variations in the scale, geometry, headspace, and mixing between the two plates formats (Supplementary file 4: Table S1). Previously, the scalability of filamentous fungi cultivation from 24-well MTPs to flask and bioreactor was demonstrated to be reliable (Kosa, Vuoristo, et al., 2018), however this might not be the same from the 96-to 24-well plates. In 24-well MTPs (extra-high deep-well plates, Enzyscreen, The Netherlands, cat. no. CR1424hd), *A. glauca* produced less biomass under standard C/N conditions and consumed slightly more glucose in the high C/N medium compared with the 96-well plate results. In contrast, *R. stolonifer* exhibited lower glucose consumption and a less pronounced pH decrease in the high C/N medium. Similar glucose consumption patterns for the same strains, medium, and 24-well plate type have been reported by Dzurendova et al. (Dzurendova et al., 2021).

It is also important to note differences in growth morphology that could explain the observed differences. *A. glauca* grew as spherical pellets, while *R. stolonifer* grew as a mix of dispersed hyphae and aggregated clumps of biomass. Notably, *R. stolonifer* tended to adhere and grow along the walls of the microplate above the liquid-air interphase. Considering the dimensions of each well and the working volume, the relative well surface available for “wall growth” of *R. stolonifer* is higher in the 96-well plates. Therefore, since the wall growth biomass is not permanently immersed in the medium, the low pH may have less impact on the fungal physiology. Evidently, the different glucose consumption also affected the biomass biochemical profile. For *A. glauca*, the higher glucose consumption correlated well with the higher lipid and organic acid production, while for *R. stolonifer* the less glucose consumed the less polyphosphate, and organic acid was produced (Fig. 4).

**Figure 4.**
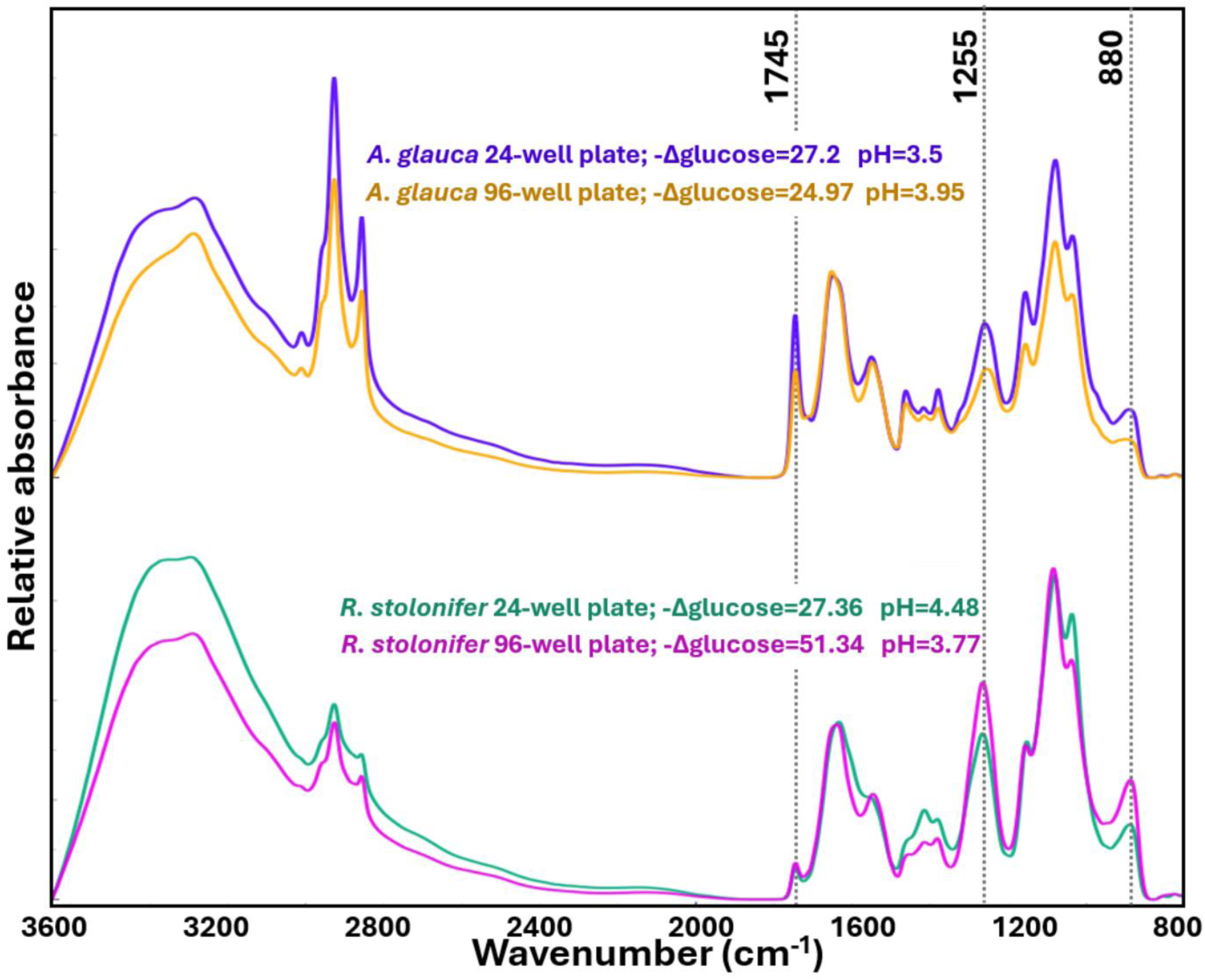
Comparison of spectra of *R. stolonifer* and *A. glauca* CCM F451 grown in high C/N media in 24-well plates *vs* 96-well plates. Note that the spectra were baseline corrected and peak-normalized at 1550 cm^-1^ for easier comparison. Vertical grey dash lines indicate relevant wavenumbers for discussion in the main text.

Revisiting the central question, the oxalic acid production by *A. glauca* can be potentially increased by certain fermentation strategies that have been applied in the case of other microorganisms. For example, *Aspergillus niger* is well-known for citric acid production but also oxalic acid that can be increased when the pH is maintained above 6 (Cameselle et al., 1998), while fumaric acid production by *Rhizopus oryzae* is enhanced with the addition of acid neutralizers (Martin-Dominguez et al., 2018). The oxalic acid production of *A. glauca* may be limited by the acidic stress (≈ 3.5 pH), which could require similar neutralizing strategies to increase its production. Future studies are needed to determine the potential and optimal conditions for oxalic acid production by *A. glauca*. Our findings suggest that high C/N ratios are not the main trigger, while the other medium components, such as the metals present as microelements, could be potential triggers.

Separately, the constituent production of oxalic acid of *A. glauca* can have implications in screening of enzymatic activities. For example, in our previous work (Bolaño Losada et al., 2025), when *A. glauca* was grown in cellobiose-based medium the extracellular protein values were high while the β-glucosidase activities were considerably lower than other Mucoromycota strains. The method used to assess β-glucosidase activity employed p-NPG as substrate, which after its cleavage releases p-nitrophenol promoting yellow coloration. However, p-nitrophenol coloration is pH-dependent, higher at alkaline conditions, requiring from addition of a buffer. However, the buffer solution might have not be sufficient to neutralize the strong acidity of oxalic acid. Indeed, the pH values of *A. glauca* in that study were the lowest among the strains for both cellobiose-and cellulose-based media (Supplementary file 4: Fig. S2).

### Production of oxalic acid in *Absidia* species

The presence of oxalic acid in *Absidia glauca* CCM F451 was particularly unusual because of its acidic strength, chelating nature, and constitutive production independently from a common triggering factor as carbon excess with nitrogen limitation. To assess whether this phenomenon also occurs in other *A. glauca* strains or if the strain CCM F451 represents a hypersecreting case, additional experiments were conducted with the strains *A. glauca* CCM F450, CCM F444 and NRRL 2799. Moreover, other species such as *Absidia heterospora* NRRL 2800, *Absidia pseudocylindrospora* CCM F52, *Absidia psychrophilia* NRRL 3044, *Absidia coerulea* NRRL 1322 were included. Initially, *Absidia blakesleeana* was also included but we noted the reclassification of this species to *Lichtheimia blakesleeana*. Nevertheless, its results are discussed and presented here as an “out of group” reference. All these species were grown in the high C/N medium.

The maximum glucose consumption for the *Absidia* and *Lichtheimia* species in the high C/N medium was between 17.85 and 27.28 g/L, with biomass production of 3.96 to 6.79 g/L (Fig. 5a), rendering biomass yields in the range of 0.21 to 0.36 (Supplementary file 4: Fig. S3). All the *Absidia* strains acidified the medium significantly (1.86 to 2.99-ΔpH values) in contrast to *Lichtheimia blakesleeana* (Fig. 5b). Oxalic acid was detected in all the tested *Absidia* species, while none for *L. blakesleeana* (Supplementary file 3). Lactic acid was also detected in most of the *Absidia* species, although at lower level. The cases of *Absidia* strains in which lactic acid production was not detected, could be due to secretion in concentrations below the detection threshold (Supplementary file 3). The best oxalic acid producer was *A. cuneospora* and the worst *A. psychrophilia*. The lower productivity observed in *A. psychrophilia* was expected since this species is a psychrophile that does not grow above 28 °C, whereas the temperature of incubation in this study was set at 25 °C as common for all the species for easier processing of the high-throughput cultivation (Hesseltine & Ellis, 1964). On the other hand, it must be noted that, according to the Czech Collection of Microorganisms, *A. glauca* CCM F451 is mating type (–) while *A. glauca* CCM F450 is mating type (+). Both mating types produced oxalic acid and therefore its production is independent from the fungal gender.

**Figure 5.**
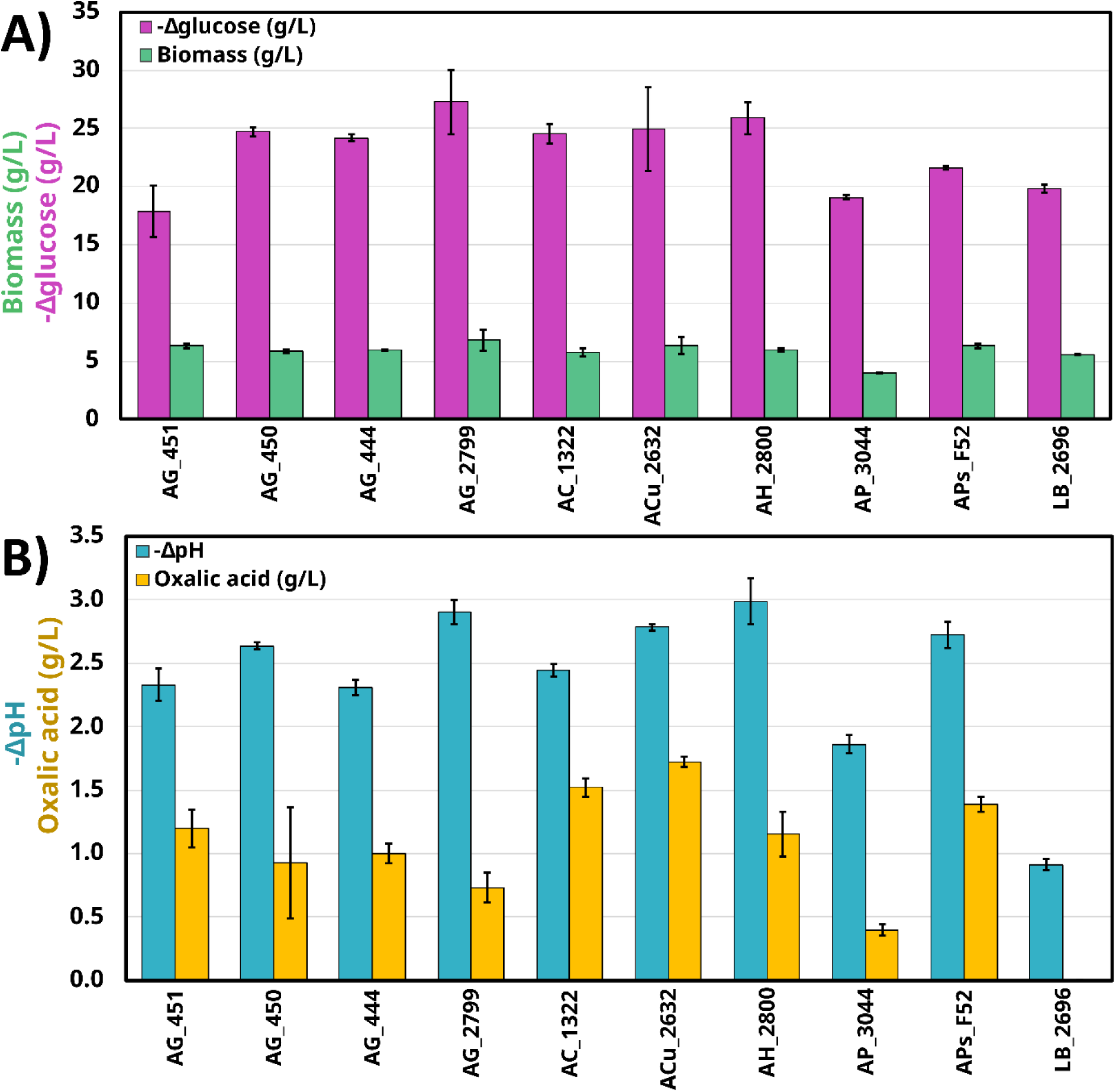
A) Consumption of glucose (-Δglucose), production of biomass, B) medium acidification (-ΔpH) and oxalic acid in supernatant from several *A. glauca* (AG) strains, *A. coerulea* (AC), *A. cuneospora* (ACu), *A. heterospora* (AH), *A. pseudocylindrospora* (APs) *A. psycrophilia* (AP) and *L. blakesleeana* (LB) grew in high C/N medium. The numbers in the labels indicate the corresponding collection number. Error bars represent the standard deviation from three biologically independent replicates.

These results, not only verifies the oxalic production in *A. glauca,* but also suggest that might be extended to the whole *Absidia* genus. Indeed, other *Absidia* species such as *Absidia spinosa* (Ceci et al., 2020) or *Absidia cylindrospora* (Albert et al., 2018) have also been reported to cause strong medium acidification. Therefore, the oxalic acid production could represent a potential discriminating feature of the whole *Absidia* genus. Additionally, earlier studies reported oxalic production in several species of *Mucor* and *Mortierella*, although the methodology was prone to multiple interferences (e.g. phosphates, lactic acid, citric acid, etc.) (Perlman, 1950). In this study we reassessed this claim with a more robust and reliable analytical method (GC-MS) and evidence to supporting it. Therefore, to our knowledge, this is also the first report of oxalic acid secretion in the phylum Mucoromycota.

### Implications of oxalic acid production in *Absidia* species

The discovery of oxalic acid production by *Absidia* species 150 years after the first definition of the genus highlights how little explored its biology is. Indeed, several new species of this genus have been discovered recently (Idnurm, 2025; Wang et al., 2025). In fungi, oxalic acid plays multiple ecological roles: bio-weathering and bio-mineralization for essential element acquisition, assistance in Fenton reactions during lignocellulose-degradation processes, competitive interactions with other fungi, among other functions (Grąz, 2024). From all these functions, lignocellulose degradation is the least likely role in *Absidia*, as Mucoromycota fungi are generally inefficient in this process. Even so, it should be considered that *A. psychrophilia* strain was isolated from ambrosia beetle glands and could play a role in wood together with other symbionts, although this remains uncertain (Hesseltine & Ellis, 1964). Biomineralization, in contrast, is a reasonable role and may explain the prevalence of *Absidia* species in oxalate-carbonate pathway ecosystems (Hervé et al., 2021). These ecosystems are of high interest because they are recognized as important natural carbon-sink systems and their foundations rely on the interactions between oxalogenic plants and fungi, and oxalotrophic bacteria. On the other hand, our observation indicates that *Absidia* accumulates also significant amounts of phosphorus that could be obtained, together with other elements, by bio-weathering of rocks assisted by the oxalic acid secretion (Gadd et al., 2014).

It is worth noting that several *Absidia* species have been used for heavy metal bioremediation due to their high tolerance and adsorption (Albert et al., 2018; Bustamante et al., 2025; Ceci et al., 2020). However, the discovery of oxalic acid secretion reveals a previously overlooked but impactful mechanism that partially explains such results. Oxalic acid, known for chelating and precipitating divalent metal ions, might contribute to *Absidia*’s metal tolerance, especially considering its constitutive secretion demonstrated in this study. Consistently, *Absidia* species have been isolated in high frequencies in metal-finishing industry sewages (Saxena et al., 2006). Future studies should aim to clarify the ecological and physiological role oxalic acid in *Absidia*.

From a biotechnological perspective, oxalic acid is used in the pharmaceutical, textile, leather, chemical, and metallurgical industries (S. Kumar et al., 2023). The demand in these sectors continues to rise, yet large-scale oxalic acid production is still primarily achieved through chemical processes. Although efficient, chemical methods often require multiple synthesis steps and the use of hazardous chemicals and fossil-based precursors. In contrast, oxalic acid can also be produced biologically in a more environmentally friendly manner. The first reports of oxalic acid production by fermentation date back to 1890s using *Aspergillus niger* (Wehmer, 1891b). Since then, there has been a moderate increase in research studies, including also other microorganisms. Nevertheless, oxalic production via fermentation has not yet been adopted industrially. However, given the increasing demand and the tendency to embrace environmentally friendly processes, the interest in oxalic acid biosynthesis is likely to increase in the future (Amenaghawon et al., 2024).

Other novel and innovative applications include biohydrometallurgy for recovery of important metals (Bahaloo-Horeh & Mousavi, 2024), myco-remediation (V. Kumar & Dwivedi, n.d.), or bio-passivation of copper and bronze artefacts for preservation of archeology artefacts (Joseph, 2021).

### New traits supporting the distinction between *Lichtheimia* and *Absidia* species

The *Absidia* genus was described in 1876 by the mycologist Philippe Édouard Léon Van Tieghem based on morphological features (van Tieghem, 1876). Later, other mycologists, re-examined the classification of the genus, using additional physiological characteristics such as growth temperatures, zygospore features and molecular analyses. These efforts ultimately led to the reclassification of *Absidia*-like taxa into three different genera: *Absidia*, *Lichtheimia* and *Lentamyces* (Hoffmann, 2010). Our study further supports the separation of *Lichtheimia* from *Absidia* genera not only due to the secretion of oxalic acid, but also by differences in biochemical composition (Fig. 6).

**Figure 6.**
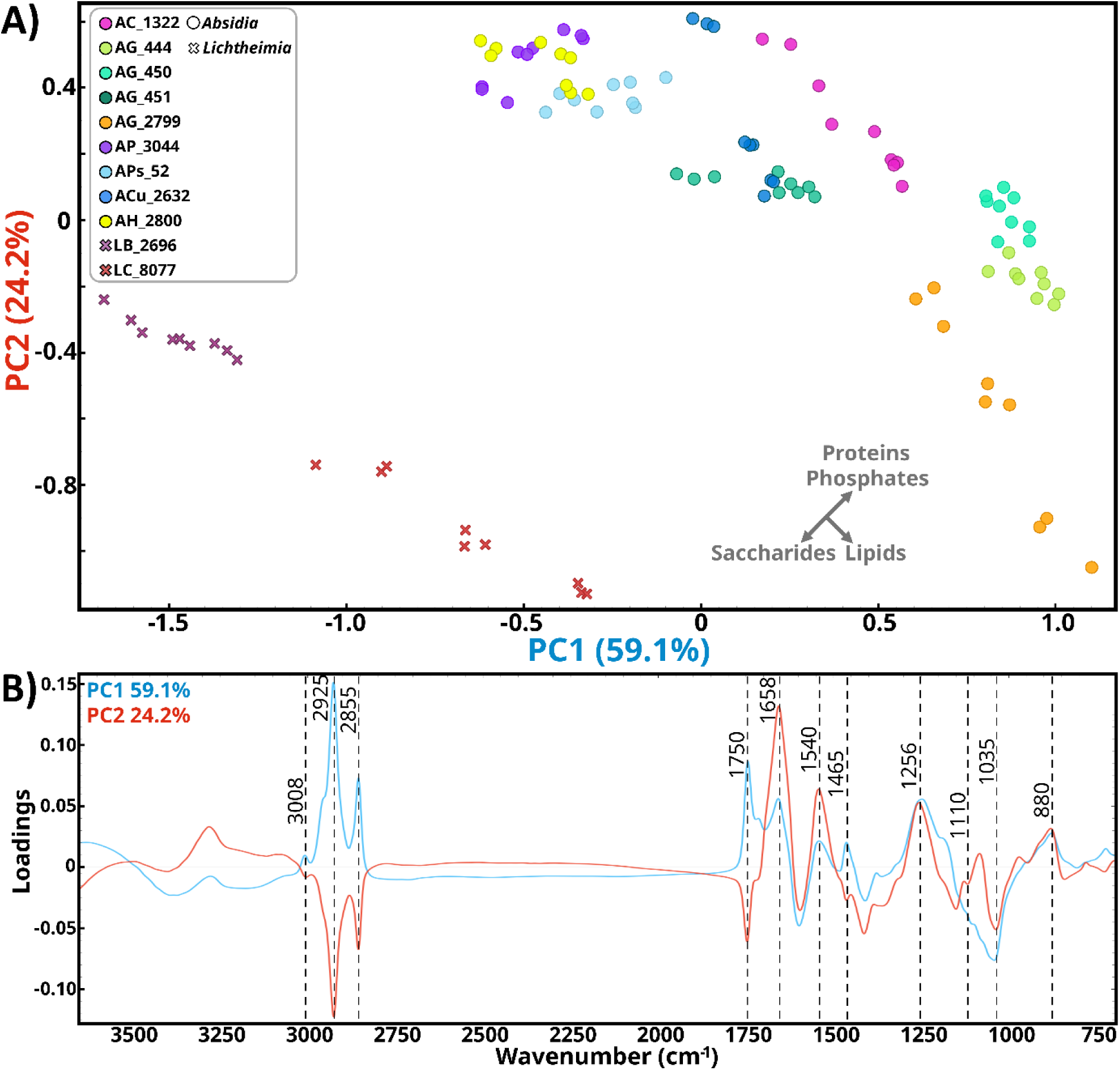
A) PCA scatter plot, and B) score plot of PC1 (blue) and PC2 (red) from the analysis of *A. glauca* (AG), *A. coerulea* (AC), *A. psycrophilia* (AP), *A. heterospora* (AH), *A. cuneospora* (ACu), *A. pseudocylindrospora* (APs), *L. blakesleeana* (LB), and *L. corymbifera* (LC) grow at high C/N medium. Each species in the PCA plot is represented by nine dots, resulting from three biological independent replicates, each measured three times. Numbers in the legend names indicate their corresponding culture collection code that is also available in Table 1. An approximate axis reflecting increases in particular biomolecules based on the principal component’s interpretation is provided in the PCA plot. Explained variance of the first five principal components were 69.1%, 21.3%, 6.4%, 1.9% and 0.7%.

The PCA analysis of FTIR spectra from the biomass of *Absidia* species, *L. corymbifera* and *L. blakesleeana* revealed a clear separation of the samples based on genus (Fig. 6a). The principal component PC1 (59.1% explained variance) displayed moderate negative loadings in saccharides-related bands, strong positive loadings in lipid-related bands (3008, 2925, 2855, 1750, and 1465 cm^-1^), moderate positive loadings in phosphate-related bands (1256 and 880 cm^-1^), and mild positive loadings in protein-related bands (1658 and 1540 cm^-1^) (Fig. 6b). On the other hand, PC2 (24.2% explained variance) displayed moderate negative loadings in lipid-related bands, strong positive loadings in protein-related bands, strong positive loadings in phosphorus-related bands (including the weak 1085 cm^-1^ peak), and strong-moderate negative loadings in saccharide-related bands (mainly 1035 and 1110 cm^-1^). In the PCA score plot *Absidia* species were separated from *Lichtheimia* along the down left diagonal direction, indicating that *Absidia* species contain higher amounts of phosphorus and proteins, while *Lichtheimia* species contain more saccharides. Differences among *Absidia* species were mainly related to lipid content and slightly to protein and phosphate content, whereas among *Lichtheimia* species were mostly due to lipid content. For comparison, a stack graph with the averaged spectra of each specie with the most relevant metabolite-related peaks is shown in Supplementary file 4: Fig. S4. Same differences were found between three *Lichtheimia* strains and ten *Absidia* strains from the study of Kosa et al. (Kosa, Zimmermann, et al., 2018), despite slight differences in the cultivation set up and medium with respect to this study. The spectra of Kosa et al. (Kosa, Zimmermann, et al., 2018) were treated and analysed in same manner that in this study and the PCA scatter and PC loading plots can be found in Supplementary file 4: Fig. S5.

To sum up, our results highlight that, in addition to oxalic acid secretion, the abundant accumulation of phosphorus could represent another physiological characteristic to differentiate *Absidia* from *Lichtheimia*. Moreover, these findings reinforce the value of FTIR spectroscopy as a powerful tool for chemical analysis of fungal biomass and for species classification based on characteristic biochemical fingerprints, an application widely recognized in the literature (Colabella et al., 2017; Shapaval et al., 2010; Tafintseva et al., 2018).

As demonstrated in our previous studies, intense phosphorus signals in this type of fungal biomasses are indicative of extensive polyphosphate accumulation (Dzurendova et al., 2021). For validation of the observed results, the Fig. 7a shows the spectra and total phosphorus content of *A. glauca* CCM F451 and *L. corymbifera* CCM F8077, together with *R. stolonifer* VKM F400 as a high phosphorus content reference. In general, the higher the relative intensity of the 1265 and 880 cm^-1^ peaks, the higher the total phosphorus content. The phosphorus content in *A. glauca* was almost twice as high as in *L. corymbifera*, supporting the statement that the two genera also differ in terms of phosphate accumulation. Additionally, a strong correlation between phosphorus and magnesium content (R^2^=0.879) (Fig. 7b) and a slightly lower with calcium content (R^2^=0.473) (Fig. 7c) was found. These observations encompass with the role of calcium and magnesium as counter-ions for polyphosphate accumulation (Choi et al., 2011; Dzurendova et al., 2021; Dzurendova, Zimmermann, Tafintseva, Kohler, Horn, et al., 2020). The content values of these elements together sodium and potassium are show in Supplementary file 4: Table S2.

**Figure 7.**
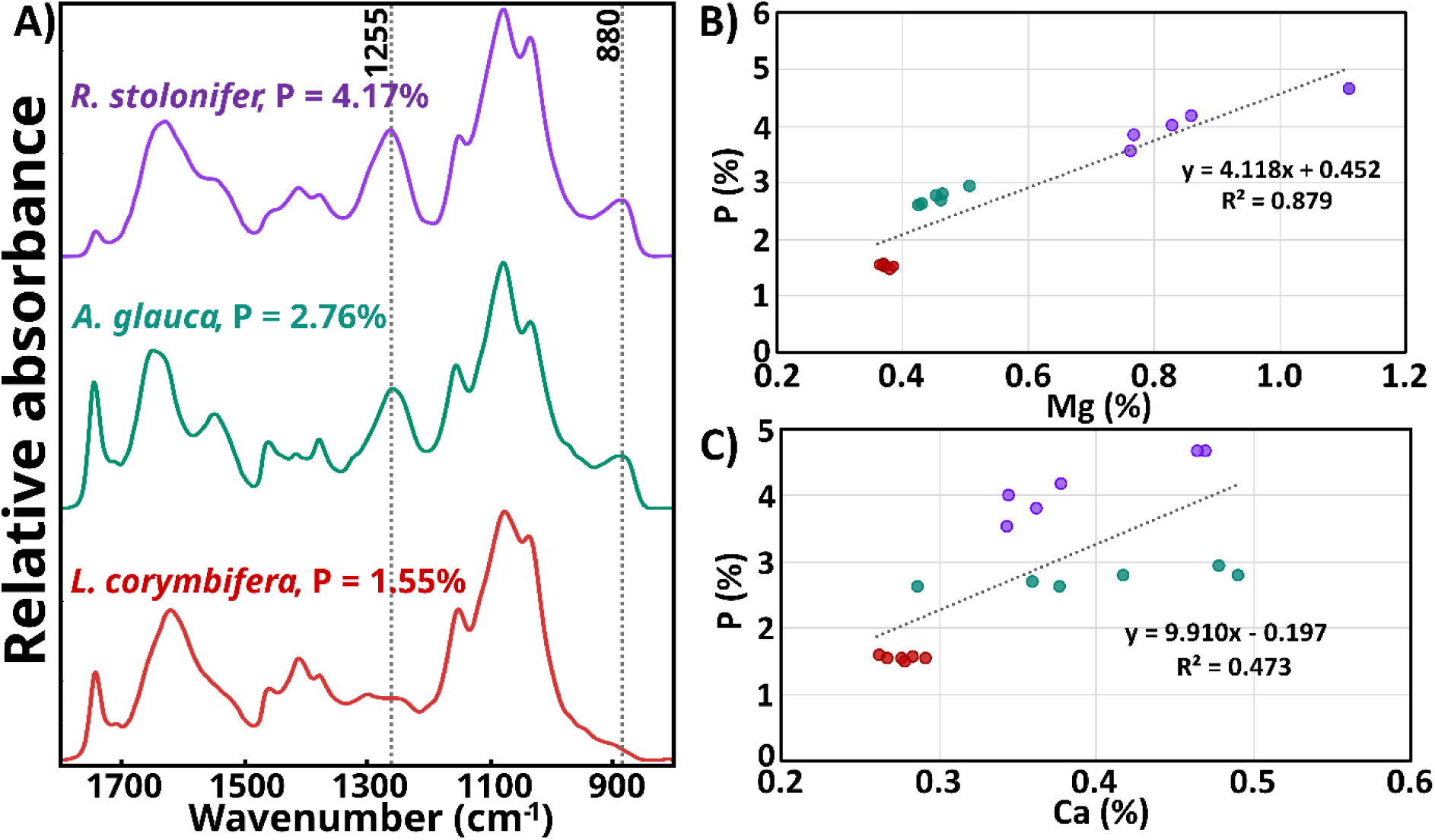
A) Stack graph of spectra from *R. stolonifer* (VKM F400) in violet, *A. glauca* (CCM F451) in green and *L. corymbifera* (CCM F8077) in red grow in 24-well plates and high C/N medium. The marked peaks at 1255 and 880 cm^-1^ correspond to P=O and P-O-P stretching vibrations, respectively. Alongside, the percentage of phosphorus (P) content is provided to validate the spectral analysis. Both spectra and phosphorus content result from the average of three biologically independent replicates. B) Correlation plot of phosphorus (P %) and magnesium (Mg %) content and C) phosphorus and calcium (Ca %) from samples of *R. stolonifer*, *A. glauca* and *L. corymbifera,* colored accordingly.

## 4. Conclusions

The carbon allocation towards different biomolecules was studied for the nine strains of Mucoromycota species. Combined high throughput approach, specifically HTS-FTIR spectroscopy and Duetz microtiter plate-based cultivation-system, proved to be particularly efficient for the aim of this study because it enabled: 1) cultivation under many conditions with rapid sample processing, and 2) the acquisition of comprehensive multichemical IR-based information. Owing to these advantages, many classical single-analyte methods could be bypassed as a first-line strategy, saving considerable time, reagents, and materials. Subsequent analyses with targeted methods can then be applied more selectively, once the likely compounds of interest have been identified. In the present work, FT-IR spectra alone provided sufficient chemical information to address part of the research questions.

As anticipated, carbon allocation strategies differed across genera and families. The results presented in this study may serve as a resource for comparative genomics or as a basis for investigating the functional roles of specific biomolecules, such as polyphosphates, through laboratory or field studies. They also highlight potential patterns for metabolic engineering, for instance identifying biomolecules that could be targeted for knockout to redirect carbon flux, or pinpointing species with potential for co-production strategies in biorefinery applications. While the selected species set was limited to strains with biotechnological relevance, this study established methodological foundations for investigating carbon allocation strategies at a broader scale, easily transferable to other filamentous fungi. Indeed, additional Mucoromycota representatives could be include in future studies to further expand the observations made in this study.

On the other hand, our findings underscore just how underexplored many Mucoromycota species remain. The discovery of constitutive oxalic acid production not only in *A. glauca* but also in other *Absidia* species emphasizes this point. This finding may carry significant evolutionary implications, as closely related genera such as *Cunninghamella* and *Lichtheimia* strains did not exhibit this characteristic. Moreover, oxalic acid could play ecologically relevant roles, for example in the persistence of these fungi in heavy metal–contaminated soils or in oxalate–carbonate pathway ecosystems. In brief, this discovery opens an important new field of investigation into the biology, ecology and biotechnology of *Absidia*.

## Acknowledgments

The authors are grateful for the gratuity of the microbial strains provided by the USDA-ARS Culture Collection (NRRL). On the other hand, to Iria Bolaño Losada and Vlad Bhlazko for help with data visualization. The featured image was created with BioRender under a licensed subscription.

## Author contributions

Conceived the research idea: CBL. Designed the experiments: CBL. Methodology: CBL, BZ, SA. Performed the experiments and acquired data: CBL, DB, SA. Discussed the results: CBL, DB, BZ, SJH, AK, VS. Analyzed the data: CBL, BZ. Writing the manuscript: CBL. Discussed and revised the manuscript: CBL, BZ, DB, SA, SJH, AK, VS. All authors read and approve the final manuscript.

## Supplementary data

Supplementary data is available at FEMS Journal online.

## Consent for publication

Does not apply.

## Data availability

The datasets supporting the conclusions of this article are available in the Zenodo repository upon reasonable request from the authors: https://10.5281/zenodo.19008411

## References

Achbergerová, L., & Nahálka, J. (2011). Polyphosphate - an ancient energy source and active metabolic regulator. Microbial Cell Factories, 10, 1–14. 10.1186/1475-2859-10-63

Albert, Q., Leleyter, L., Lemoine, M., Heutte, N., Rioult, J. P., Sage, L., Baraud, F., & Garon, D. (2018). Comparison of tolerance and biosorption of three trace metals (Cd, Cu, Pb) by the soil fungus Absidia cylindrospora. Chemosphere, 196, 386–392. 10.1016/j.chemosphere.2017.12.156

Amenaghawon, A. N., Ayere, J. E., Amune, U. O., Otuya, I. C., Abuga, E. C., Anyalewechi, C. L., Okoro, O. V., Okolie, J. A., Oyefolu, P. K., Eshiemogie, S. O., Osahon, B. E., Omede, M., Eshiemogie, S. A., Igemhokhai, S., Okedi, M. O., Kusuma, H. S., Muojama, O. E., Shavandi, A., & Darmokoesoemo, H. (2024). A comprehensive review of recent advances in the applications and biosynthesis of oxalic acid from bio-derived substrates. Environmental Research, 251(P2), 118703. 10.1016/j.envres.2024.118703

Andrade-Linares, D. R., Veresoglou, S. D., & Rillig, M. C. (2016). Temperature priming and memory in soil filamentous fungi. Fungal Ecology, 21, 10–15. 10.1016/j.funeco.2016.02.002

Bahaloo-Horeh, N., & Mousavi, S. M. (2024). Analyzing the Effects of Culture Media Additives on Oxalic Acid Bioproduction for Use in Metal Bioleaching. Waste and Biomass Valorization, 15(5), 2687–2703. 10.1007/s12649-023-02381-4

Barbosa De Lima, M. A., Do Nascimento, A. E., De Souza, W., Fukushima, K., & De Campos-Takaki, G. M. (2003). Effects of phosphorus on polyphosphate accumulation by Cunninghamella elegans. Brazilian Journal of Microbiology, 34(4), 363–372. 10.1590/s1517-83822003000400016

Birkby, K., & Preece, T. (1988). Mycotechnology: Calcium oxalate crystals on the sporangium of Pilobolus. Topics in Catalysis, 2(2), 68–69. 10.1016/S0269-915X(88)80009-0

Bolaño Losada, C., Slaný, O., Byrtusová, D., Zimmermann, B., Horn, S. J., Kohler, A., & Shapaval, V. (2025). Compatible traits of oleaginous Mucoromycota fungi for lignocellulose-based simultaneous saccharification and fermentation. Biotechnology for Biofuels and Bioproducts, 18(1). 10.1186/s13068-025-02621-w

Bolaño-Losada, C., Zimmermann, B., Horn, S. J., Kohler, A., & Shapaval, V. (2025). Effect of hydrogen peroxide and carbon-to-nitrogen ratio on growth and biochemical profile in oleaginous Mucoromycota. Microbial Cell Factories, 24(1). 10.1186/s12934-025-02863-1

Bonfante, P., & Venice, F. (2020). Mucoromycota: going to the roots of plant-interacting fungi. Fungal Biology Reviews, 34(2), 100–113. 10.1016/j.fbr.2019.12.003

Bustamante, N., Ortiz, J., Soto, J., Dias, N. B., Vega, C., & Arriagada-Escamilla, C. (2025). Evaluating Fungal Metal Tolerance Using MALDI-TOF MS: A Rapid Alternative to Conventional MIC Methods. Journal of Basic Microbiology. 10.1002/jobm.70064

Büttner, H., Rassbach, J., Schultz, C., Popp, J., Gressler, M., & Hertweck, C. (2024). Beneficial Soil Fungus Kills Predatory Nematodes with Dehydropeptides Translocating into the Animal Gut. Journal of the American Chemical Society, 146(50), 34702–34710. 10.1021/jacs.4c12989

Cameselle, C., Bohlmann, J., Núñez, M., & Lema, J. M. (1998). Oxalic acid production by Aspergillus niger Part I: Influence of sucrose and milk whey as carbon source. Bioprocess Engineering, 19, 247–252.

Ceci, A., Spinelli, V., Massimi, L., Canepari, S., & Persiani, A. M. (2020). Fungi and arsenic: Tolerance and bioaccumulation by soil saprotrophic species. Applied Sciences (Switzerland), 10(9). 10.3390/app10093218

Chang, L., Chen, H., Yang, B., Chen, H., & Chen, W. (2023). Redistributing carbon flux by impairing saccharide synthesis to enhance lipid yield in oleaginous fungus Mortierella alpina. ACS Synthetic Biology, 12(6), 1750–1760. 10.1021/acssynbio.3c00046

Chatzifragkou, A., Fakas, S., Galiotou-Panayotou, M., Komaitis, M., Aggelis, G., & Papanikolaou, S. (2010). Commercial sugars as substrates for lipid accumulation in Cunninghamella echinulata and Mortierella isabellina fungi. European Journal of Lipid Science and Technology, 112(9), 1048–1057. 10.1002/ejlt.201000027

Chen, H., Chen, H., Lu, H., Tang, X., Zhang, H., Chen, Y. Q., & Chen, W. (2021). Carbohydrate analysis of Mortierella alpina by colorimetry and HPLC–ELSD to reveal accumulation differences of sugar and lipid. Biotechnology Letters, 43(7), 1289–1301. 10.1007/s10529-021-03120-2

Cherubini, F. (2010). The biorefinery concept: Using biomass instead of oil for producing energy and chemicals. Energy Conversion and Management, 51(7), 1412–1421. 10.1016/j.enconman.2010.01.015

Choi, H.-J., Yu, S.-W., Lee, S.-M., & Yu, S.-Y. (2011). Effects of Potassium and Magnesium in the Enhanced Biological Phosphorus Removal Process Using a Membrane Bioreactor. Water Environment Research, 83(7), 613–621.

Christy, A. J., Wang, R., Wikandari, R., & Taherzadeh, M. J. (2026). Mineral composition of cultivation media affects mineral bioaccessibility of edible filamentous fungus Rhizopus oligosporus. Future Foods, 13, 100960. 10.1016/j.fufo.2026.100960

Colabella, C., Corte, L., Roscini, L., Shapaval, V., Kohler, A., Tafintseva, V., Tascini, C., & Cardinali, G. (2017). Merging FT-IR and NGS for simultaneous phenotypic and genotypic identification of pathogenic Candida species. PLoS ONE, 12(12). 10.1371/journal.pone.0188104

Crognale, S., Russo, C., Petruccioli, M., & D’annibale, A. (2022). Chitosan Production by Fungi: Current State of Knowledge, Future Opportunities and Constraints. In Fermentation (Vol. 8, Number 2). MDPI. 10.3390/fermentation8020076

Danilova, O. A., Ianutsevich, E. A., Kochkina, G. A., & Tereshina, V. M. (2025). Adaptation of the psychrophilic Mucor psychrophilus (Mucorales, Mucoromycota) to lower temperatures and under conditions of heat and osmotic shocks. Fungal Biology, 129(2), 101532. 10.1016/j.funbio.2024.101532

Das, R. K., Brar, S. K., & Verma, M. (2016). Fumaric Acid: production and applications aspects. In S. K. Brar, S. J. Sarma, & K. Pakshirajan (Eds.), Platform Chemical Biorefinery: Future Green Chemistry (1st ed.). Elsevier. 10.1016/B978-0-12-802980-0.00008-0

Demšar, J., Curk, T., Erjavec, A., Gorup, Č., Hočevar, T., Milutinovič, M., Možina, M., Polajnar, M., Toplak, M., Starič, A., Štajdohar, M., Umek, L., Žagar, L., Žbontar, J., Žitnik, M., & Zupan, B. (2013). Orange: Data mining toolbox in Python. Journal of Machine Learning Research, 14, 2349–2353.

Do Nascimento Shari’a, A. E., Do Nascimento, A. E., Barbosa De Lima, M. A., De Campos-Takaki, G. M., & De Souza, W. (2002). Polyphosphate in Zygomycetes: A cytochemical study. Brazilian Journal of Microbiology, 33(2), 119–126. 10.1590/s1517-83822002000200004

dos Santos, J. A., Meyer, E., & Sette, L. D. (2020). Fungal community in antarctic soil along the retreating collins glacier (Fildes peninsula, King George Island). Microorganisms, 8(8), 1–15. 10.3390/microorganisms8081145

Dzurendová, S., Almklov Magnussen, E., Shapaval, V., Kohler, A., & Zimmermann, B. (2026). Vibrational spectroscopy methodology for profiling of mixed carbon metabolism in carotenogenic and oleaginous Mucor circinelloides. Microbial Cell Factories. 10.1186/s12934-026-02963-6

Dzurendova, S., Bolaño Losada, C., Dupuy-Galet, B. X., Slany, O., Fjaer, K., Di Bartolomeo, F., Markussen, S., Wentzel, A., Várnai, A., Degn Hansen, L., Horn, S. J., Kohler, A., Shapaval, V., & Zimmermann, B. (2025). Application of vibrational spectroscopies as process analytical techniques for monitoring fermentation and the conversion of lignocellulosic biomass by oleaginous filamentous fungi. Microbial Cell Factories, 24(1). 10.1186/s12934-025-02868-w

Dzurendova, S., Losada, C. B., Dupuy-Galet, B. X., Fjær, K., & Shapaval, V. (2022). Mucoromycota fungi as powerful cell factories for modern biorefinery. Applied Microbiology and Biotechnology, 1–15.

Dzurendová, S., Shapaval, V., Tafintseva, V., Kohler, A., Byrtusová, D., Szotkowski, M., Márová, I., & Zimmermann, B. (2021). Assessment of biotechnologically important filamentous fungal biomass by Fourier transform Raman spectroscopy. International Journal of Molecular Sciences, 22(13). 10.3390/ijms22136710

Dzurendova, S., Zimmermann, B., Kohler, A., Reitzel, K., Nielsen, U. G., Dupuy--Galet, B. X., Leivers, S., Horn, S. J., & Shapaval, V. (2021). Calcium affects polyphosphate and lipid accumulation in Mucoromycota fungi. Journal of Fungi, 7(4). 10.3390/jof7040300

Dzurendova, S., Zimmermann, B., Kohler, A., Tafintseva, V., Slany, O., Certik, M., & Shapaval, V. (2020). Microcultivation and FTIR spectroscopy-based screening revealed a nutrient-induced co-production of high-value metabolites in oleaginous Mucoromycota fungi. PLoS ONE, 15(6 June), 1–34. 10.1371/journal.pone.0234870

Dzurendova, S., Zimmermann, B., Tafintseva, V., Kohler, A., Ekeberg, D., & Shapaval, V. (2020). The influence of phosphorus source and the nature of nitrogen substrate on the biomass production and lipid accumulation in oleaginous Mucoromycota fungi. Applied Microbiology and Biotechnology, 104(18), 8065–8076. 10.1007/s00253-020-10821-7

Dzurendova, S., Zimmermann, B., Tafintseva, V., Kohler, A., Horn, S. J., & Shapaval, V. (2020). Metal and phosphate ions show remarkable influence on the biomass production and lipid accumulation in oleaginous Mucor circinelloides. Journal of Fungi, 6(4), 1–23. 10.3390/jof6040260

Elbein, A. D. (2009). Cytoplasmic carbohydrate molecules: trehalose and glycogen. In A. P. Moran, O. Holst, P. J. Brennan, & M. von Itzstein (Eds.), Microbial glycobiology. Structures, relevance and applications. (pp. 185–202). Elsevier.

Elisa, A., Souza, D. A., Luiz, A., Monteiro, C., Santiago, D. A., Cupertino, A., Brito, D. Q., Alves, J., Cristina, M., De Souza, M., Machado, A. R., Elisa, A., & Souza, D. A. (2025). New reports of Rhizopus and Mucor (Mucorales, Mucoromycota) causing post - harvest rot in tropical fruits in Brazil. Journal of Plant Diseases and Protection, 132(1), 1–16. 10.1007/s41348-024-01035-4

Fröhlich-Nowoisky, J., Hill, T. C. J., Pummer, B. G., Yordanova, P., Franc, G. D., & Pöschl, U. (2015). Ice nucleation activity in the widespread soil fungus Mortierella alpina. Biogeosciences, 12(4), 1057–1071. 10.5194/bg-12-1057-2015

Gadd, G. M., Bahri-Esfahani, J., Li, Q., Rhee, Y. J., Wei, Z., Fomina, M., & Liang, X. (2014). Oxalate production by fungi: Significance in geomycology, biodeterioration and bioremediation. In Fungal Biology Reviews (Vol. 28, Numbers 2–3, pp. 36–55). Elsevier Ltd. 10.1016/j.fbr.2014.05.001

Gonçalves, V. N., Vaz, A. B. M., Rosa, C. A., & Rosa, L. H. (2012). Diversity and distribution of fungal communities in lakes of Antarctica. FEMS Microbiology Ecology, 82(2), 459–471. 10.1111/j.1574-6941.2012.01424.x

Grąz, M. (2024). Role of oxalic acid in fungal and bacterial metabolism and its biotechnological potential. World Journal of Microbiology and Biotechnology, 40(6), 1–11. 10.1007/s11274-024-03973-5

Guo, S., Kohler, A., Zimmermann, B., Heinke, R., Stöckel, S., Rösch, P., Popp, J., & Bocklitz, T. (2018). Extended Multiplicative Signal Correction Based Model Transfer for Raman Spectroscopy in Biological Applications. Analytical Chemistry, 90(16), 9787–9795. 10.1021/acs.analchem.8b01536

Hervé, V., Simon, A., Randevoson, F., Cailleau, G., Rajoelison, G., Razakamanarivo, H., Bindschedler, S., Verrecchia, E., & Junier, P. (2021). Functional diversity of the litter-associated fungi from an oxalate–carbonate pathway ecosystem in madagascar. Microorganisms, 9(5), 1–12. 10.3390/microorganisms9050985

Hesseltine, C. W., & Ellis, J. J. (1964). The Genus Absidia: Gongronella and Cylindrical-Spored Species of Absidia. Mycologia, 56(4), 568–601. 10.1080/00275514.1964.12018145

Hoffmann, K. (2010). Identification of the genus Absidia (Mucorales, Zygomycetes): a comprehensive taxonomic revision. In Y. Gherbawy & K. Voigt (Eds.), Molecular identification of fungi (pp. 439–460). Springer.

Hu, H., Li, P., Li, S., Wang, X., Mohamed, H., López-García, S., Liu, Q., Garre, V., & Song, Y. (2024). The role of areA in lipid accumulation in high lipid-producing fungus Mucor circinelloides WJ11. Biochimica et Biophysica Acta-Molecular and Cell Biology of Lipids, 1869(2). 10.1016/j.bbalip.2023.159450

Idnurm, A. (2025). Absidia urquhartii (Mucoromycota) is a new species that is prevalent in Victoria, Australia. Phytotaxa, 687(1), 1–8. 10.11646/phytotaxa.687.1.1

Jiao, W., Liu, X., Li, Y., Li, B., Du, Y., Zhang, Z., Chen, Q., & Fu, M. (2022). Organic acid, a virulence factor for pathogenic fungi, causing postharvest decay in fruits. Molecular Plant Pathology, 23(2), 304–312. 10.1111/mpp.13159

Joseph, E. (2021). Biopassivation Method for the Preservation of Copper and Bronze Artefacts. Frontiers in Materials, 7. 10.3389/fmats.2020.613169

Karimi, K., Brandberg, T., Edebo, L., & Taherzadeh, M. J. (2005). Fed-batch cultivation of Mucor indicus in dilute-acid lignocellulosic hydrolyzate for ethanol production. Biotechnology Letters, 27(18), 1395–1400. 10.1007/s10529-005-0688-2

Kohler, A., Heitmann Solheim, J., Tafintseva, V., Zimmermann, B., & Shapaval, V. (2020). Model-Based Pre-Processing in Vibrational Spectroscopy. In S. Brown, R. Tauler, & B. Walczak (Eds.), Comprehensive Biotechnology (2nd ed., Vol. 4, Number January 1996, pp. 83–100). Elsevier.

Kosa, G., Shapaval, V., Kohler, A., & Zimmermann, B. (2017). FTIR spectroscopy as a unified method for simultaneous analysis of intra-and extracellular metabolites in high-throughput screening of microbial bioprocesses. Microbial Cell Factories, 16(1), 1–11. 10.1186/s12934-017-0817-3

Kosa, G., Vuoristo, K. S., Horn, S. J., Zimmermann, B., Afseth, N. K., Kohler, A., & Shapaval, V. (2018). Assessment of the scalability of a microtiter plate system for screening of oleaginous microorganisms. Applied Microbiology and Biotechnology, 102(11), 4915– 4925. 10.1007/s00253-018-8920-x

Kosa, G., Zimmermann, B., Kohler, A., Ekeberg, D., Afseth, N. K., Mounier, J., & Shapaval, V. (2018). High-throughput screening of Mucoromycota fungi for production of low-and high-value lipids. Biotechnology for Biofuels, 11(1), 1–17. 10.1186/s13068-018-1070-7

Kumar, S., Panwar, P., Sehrawat, N., Upadhyay, S., Kumar, S., Kumar, A., Singh, M., & Yadav, M. (2023). Oxalic acid: recent developments for cost-effective microbial production. Physical Sciences Reviews, 9(2), 891–907. 10.1515/psr-2022-0167

Kumar, V., & Dwivedi, S. K. (n.d.). Mycoremediation of heavy metals: processes, mechanisms, and affecting factors. 10.1007/s11356-020-11491-8/Published

Langseter, A. M., Dzurendova, S., Shapaval, V., Kohler, A., Ekeberg, D., & Zimmermann, B. (2021). Evaluation and optimisation of direct transesterification methods for the assessment of lipid accumulation in oleaginous filamentous fungi. Microbial Cell Factories, 20(1), 1–15. 10.1186/s12934-021-01542-1

Leifheit, E. F., Camenzind, T., Lehmann, A., Andrade-Linares, D. R., Fussan, M., Westhusen, S., Wineberger, T. M., & Rillig, M. C. (2024). Fungal traits help to understand the decomposition of simple and complex plant litter. FEMS Microbiology Ecology, 100(5). 10.1093/femsec/fiae033

Leman, J. (2009). Lipids, Production. Encyclopedia of Microbiology, Third Edition, 393–406. 10.1016/B978-012373944-5.00170-X

Liaud, N., Giniés, C., Navarro, D., Fabre, N., Crapart, S., Herpoël-Gimbert, I., Levasseur, A., Raouche, S., & Sigoillot, J.-C. (2014). Exploring fungal biodiversity: organic acid production by 66 strains of filamentous fungi. Fungal Biology and Biotechnology, 1, 1. doi:10.1186/s40694-014-0001-z

Longacre, A., Reimers, J. M., Gannon, J. E., & Wright, B. E. (1997). Flux analysis of glucose metabolism in rhizopus oryzae for the purpose of increasing lactate yields. Fungal Genetics and Biology, 21(1), 30–39. 10.1006/fgbi.1996.0952

Losada, C. B., Di Bartolomeo, F., Wentzel, A., Markussen, S., Dzurendova, S., Zimmermann, B., Fjaer, K., Slany, O., Várnai, A., Hansen, L. D., Horn, S. J., Eijsink, V. G. H., Patil, V., & Shapaval, V. (2025). Simultaneous production of fatty acids and amino polysaccharides from Norway spruce hydrolysates using oleaginous Mucor circinelloides. Scientific Reports, 15(1). 10.1038/s41598-025-98549-0

Lübbehüsen, T. L., Nielsen, J., & McIntyre, M. (2004). Aerobic and anaerobic ethanol production by Mucor circinelloides during submerged growth. Applied Microbiology and Biotechnology, 63(5), 543–548. 10.1007/s00253-003-1394-4

Martin-Dominguez, V., Estevez, J., De Borja Ojembarrena, F., Santos, V. E., & Ladero, M. (2018). Fumaric acid production: A biorefinery perspective. Fermentation, 4(2). 10.3390/fermentation4020033

Matsumotoo Naoyukii, & Hsiang Tom. (2016). Snow Mold. The Battle Under Snow Between Fungal Pathogens and Their Plant Hosts. Springer Science+Business Media Singapore. DOI10.1007/978-981-10-0758-3SchoolofEnvironmentalSciencesUniversityofGuelphGuelph,OntarioCanada

McIntyre, M., Breum, J., Arnau, J., & Nielsen, J. (2002). Growth physiology and dimorphism of Mucor circinelloides (syn. racemosus) during submerged batch cultivation. Applied Microbiology and Biotechnology, 58(4), 495–502. 10.1007/s00253-001-0916-1

Millati, R., Edebo, L., & Taherzadeh, M. J. (2005). Performance of Rhizopus, Rhizomucor, and Mucor in ethanol production from glucose, xylose, and wood hydrolyzates. Enzyme and Microbial Technology, 36(2–3), 294–300. 10.1016/j.enzmictec.2004.09.007

Molina, L., Rajchenberg, M., de Errasti, A., Vogel, B., Coetzee, M. P. A., Aime, M. C., & Pildain, M. B. (2023). Sapwood mycobiome varies across host, plant compartment and environments in Nothofagus forests from Northern Patagonia. Molecular Ecology, 32(23), 6599–6618. 10.1111/mec.16771

Muszewska, A., Okrasińska, A., Steczkiewicz, K., Drgas, O., Orłowska, M., Perlińska-Lenart, U., Aleksandrzak-Piekarczyk, T., Szatraj, K., Zielenkiewicz, U., Piłsyk, S., Malc, E., Mieczkowski, P., Kruszewska, J. S., Bernat, P., & Pawłowska, J. (2021). Metabolic

Potential, Ecology and Presence of Associated Bacteria Is Reflected in Genomic Diversity of Mucoromycotina. Frontiers in Microbiology, 12. 10.3389/fmicb.2021.636986

Perini, L., Gostinčar, C., & Gunde-Cimerman, N. (2019). Fungal and bacterial diversity of Svalbard subglacial ice. Scientific Reports, 9(1), 1–15. 10.1038/s41598-019-56290-5

Perlman, D. (1950). Observations on the production of ethanol by fungi and yeasts. American Journal of Botany, 37(3), 237–241. http://www.jstor.org/stable/2437908

Petrasch, S., Silva, C. J., Mesquida-Pesci, S. D., Gallegos, K., van den Abeele, C., Papin, V., Fernandez-Acero, F. J., Knapp, S. J., & Blanco-Ulate, B. (2019). Infection strategies deployed by Botrytis cinerea, Fusarium acuminatum, and Rhizopus stolonifer as a function of tomato fruit ripening stage. Frontiers in Plant Science, 10(March), 1–17. 10.3389/fpls.2019.00223

Qin, D., Wang, L., Han, M., Wang, J., Song, H., Yan, X., Duan, X., & Dong, J. (2018). Effects of an endophytic fungus umbelopsis dimorphaon the secondary metabolites of host-plant kadsura angustifolia. Frontiers in Microbiology, 9(NOV). 10.3389/fmicb.2018.02845

Ratledge, C. (2006). Microbial production of gamma-linolenic acid. In C. Akoh (Ed.), Handbook of Functional Lipids. (2). CRC press.

Safaei, Z., Karimi, K., & Zamani, A. (2016). Impact of phosphate, potassium, yeast extract, and trace metals on chitosan and metabolite production by Mucor indicus. International Journal of Molecular Sciences, 17(9). 10.3390/ijms17091429

Saxena, P., Bhattacharyya, A. K., & Mathur, N. (2006). Nickel tolerance and accumulation by filamentous fungi from sludge of metal finishing industry. Geomicrobiology Journal, 23(5), 333–340. 10.1080/01490450600762522

Schmidt, S. K., Wilson, K. L., Meyer, A. F., Gebauer, M. M., & King, A. J. (2008). Phylogeny and ecophysiology of opportunistic “snow molds” from a subalpine forest ecosystem. Microbial Ecology, 56(4), 681–687. 10.1007/s00248-008-9387-6

Shah, A. M., Mohamed, H., Fazili, A. B. A., Yang, W., & Song, Y. (2022). Investigating the Effect of Alcohol Dehydrogenase Gene Knockout on Lipid Accumulation in Mucor circinelloides WJ11. Journal of Fungi, 8(9). 10.3390/jof8090917

Shapaval, V., Deniset-Besseau, A., Dubava, D., Dzurendova, S., Heitmann Solheim, J., & Kohler, A. (2023). Multiscale spectroscopic analysis of lipids in dimorphic and oleaginous Mucor circinelloides accommodate sustainable targeted lipid production. Fungal Biology and Biotechnology, 10(1), 1–18. 10.1186/s40694-023-00148-z

Shapaval, V., Møretrø, T., Suso, H. P., Åsli, A. W., Schmitt, J., Lillehaug, D., Martens, H., Böcker, U., & Kohler, A. (2010). A high-throughput microcultivation protocol for FTIR spectroscopic characterization and identification of fungi. Journal of Biophotonics, 3(8– 9), 512–521. 10.1002/jbio.201000014

Tafintseva, V., Vigneau, E., Shapaval, V., Cariou, V., Qannari, E. M., & Kohler, A. (2018). Hierarchical classification of microorganisms based on high-dimensional phenotypic data. Journal of Biophotonics, 11(3). 10.1002/jbio.201700047

Takahashi, T., & Sakaguchi, K. (1927). On the physiology of Rhizopus species. Part IV. The relation beween the carbon source and the production of alcohol and acids, especially the formation of succinic acid from glycine. Bulletin of the Agricultural Chemical Society of Japan, 3(4–6), 42–45.

Tang, X., Zan, X., Zhao, L., Chen, H., Chen, Y. Q., Chen, W., Song, Y., & Ratledge, C. (2016). Proteomics analysis of high lipid-producing strain Mucor circinelloides WJ11: an explanation for the mechanism of lipid accumulation at the proteomic level. Microbial Cell Factories, 1–16. 10.1186/s12934-016-0428-4

Tang, X., Zhao, L., Chen, H., Chen, Y. Q., Chen, W., Song, Y., & Ratledge, C. (2015). Complete genome sequence of a high lipid-producing strain of Mucor circinelloides WJ11 and comparative genome analysis with a low lipid-producing strain CBS 277.49. PLoS ONE, 10(9), 1–11. 10.1371/journal.pone.0137543

Telagathoti, A., Probst, M., & Peintner, U. (2021). Habitat, Snow-Cover and Soil pH, Affect the Distribution and Diversity of Mortierellaceae Species and Their Associations to Bacteria. Frontiers in Microbiology, 12. 10.3389/fmicb.2021.669784

Thevelein, J. M. (1984). Regulation of trehalose mobilization in fungi. Microbiological Reviews, 48(1), 42–59. 10.1128/mmbr.48.1.42-59.1984

Toplak, M., Birarda, G., Read, S., Sandt, C., Rosendahl, S. M., Vaccari, L., Demšar, J., & Borondics, F. (2017). Infrared Orange: Connecting Hyperspectral Data with Machine Learning. Synchrotron Radiation News, 30(4), 40–45. 10.1080/08940886.2017.1338424

Uehling, J. K., Entler, M. R., Meredith, H. R., Millet, L. J., Timm, C. M., Aufrecht, J. A., Bonito, G. M., Engle, N. L., Labbé, J. L., Doktycz, M. J., Retterer, S. T., Spatafora, J. W., Stajich, J. E., Tschaplinski, T. J., & Vilgalys, R. J. (2019). Microfluidics and Metabolomics Reveal Symbiotic Bacterial–Fungal Interactions Between Mortierella elongata and Burkholderia Include Metabolite Exchange. Frontiers in Microbiology, 10. 10.3389/fmicb.2019.02163

Urbanus, A. J. F. L. M., Van Den Ende, H., Koch, B., Mycologia, S., Aug, J., & Aug, N. J. (1978). Calcium Oxalate Crystals in the Wall of Mucor mucedo REFERENCES Linked references are available on JSTOR for this article: reference # references _ tab _ contents You may need to log in to JSTOR to access the linked references. 70(4), 829–842.

van Tieghem, P. É. L. (1876). Troisième mémoire sur les Mucorinées. Annales Des Siences Naturalles, Botanique, VI(4), 312–399.

Vellozo-Echevarría, T., Barrett, K., Vuillemin, M., & Meyer, A. S. (2024). Mini-Review: The Distinct Carbohydrate Active Enzyme Secretome of Rhizopus spp. Represents Fitness for Mycelium Remodeling and Solid-State Plant Food Fermentation. In ACS Omega (Vol. 9, Number 32, pp. 34185–34195). American Chemical Society. 10.1021/acsomega.4c04378

Wang, Y.-X., Ding, Z.-Y., Ji, X.-Y., Meng, Z., & Liu, X.-Y. (2025). Unveiling Species Diversity Within Early-Diverging Fungi from China VI: Four Absidia sp. nov. (Mucorales) in Guizhou and Hainan. Microorganisms, 13(6), 1315. 10.3390/microorganisms13061315

Wehmer, C. (1891a). Entstehung und physiologische Bedentung der Oxalsäure im Stoffwechsel einiger Pilze: I. Botanische Zeitung, 233–239.

Wehmer, C. (1891b). Entstehung und physiologische Bedeutung der Oxalsäure im Stoffwechsel einiger Pilze XIII. Botanische Zeitung, 417.

Werner, T. P., Amrhein, N., & Freimoser, F. M. (2007). Specific localization of inorganic polyphosphate (poly P) in fungal cell walls by selective extraction and immunohistochemistry. Fungal Genetics and Biology, 44(9), 845–852. 10.1016/j.fgb.2007.01.008

Whitney, K. D., & Arnott, H. J. (1986). Morphology and Development of Calcium Oxalate Deposits in Gilbertella Persicaria (Mucorales). Mycologia, 78(1), 42–51. 10.1080/00275514.1986.12025202

Wierzchowska, K., Zieniuk, B., Nowak, D., & Fabiszewska, A. (2021). Phosphorus and nitrogen limitation as a part of the strategy to stimulate microbial lipid biosynthesis. Applied Sciences (Switzerland), 11(24). 10.3390/app112411819

Zhao, H., Lv, M., Liu, Z., Zhang, M., Wang, Y., Ju, X., Song, Z., Ren, L., Jia, B., Qiao, M., & Liu, X. (2021). High-yield oleaginous fungi and high-value microbial lipid resources from Mucoromycota. Bioenergy Research, 14(4), 1196–1206. 10.1007/s12155-020-10219-3

Zhao, H., Nie, Y., Zong, T. K., Wang, K., Lv, M. L., Cui, Y. J., Tohtirjap, A., Chen, J. J., Zhao, C. L., Wu, F., Cui, B. K., Yuan, Y., Dai, Y. C., & Liu, X. Y. (2023). Species diversity, updated classification and divergence times of the phylum Mucoromycota. In Fungal Diversity (Vol. 123, Number 1). Springer Netherlands. 10.1007/s13225-023-00525-4

