## Supplementary file 2 for "Carbon allocation strategies across Mucoromycota and the discovery of oxalic acid secretion in *Absidia*"

### Carbon allocation strategies across Mucoromycota and the discovery of oxalic acid secretion in *Absidia*

Cristian Bolaño Losada[1], Boris Zimmermann[1], Dana Byrtusova[1], Sven Andreas Högfeldt[1], Svein Jarle Horn[2], Achim Kohler[1], Volha Shapaval[1]

[1] Faculty of Science and Technology, Norwegian University of Life Sciences (NMBU), 1432 Ås, Norway

[2] Faculty of Chemistry, Biotechnology and Food Science, Norwegian University of Life Sciences (NMBU), 1432 Ås, Norway.

This additional file contains several figures with spectra from each strain from the Set I grown on media with different C/N ratios: low (red), standard (green) and high (blue). For comparison, the spectra were preprocessed by data truncation from 3650 to 700 $cm^{-1}$ and extended multiplicative signal correction (EMSC) (an MSC model extended by linear and quadratic components and polynomial order 2). The spectra were averaged for each C/N condition and the biologically independent replicates of the corresponding strain. The average spectra are displayed together the corresponding standard deviation in shaded color.

*Absidia glauca CCM F451*

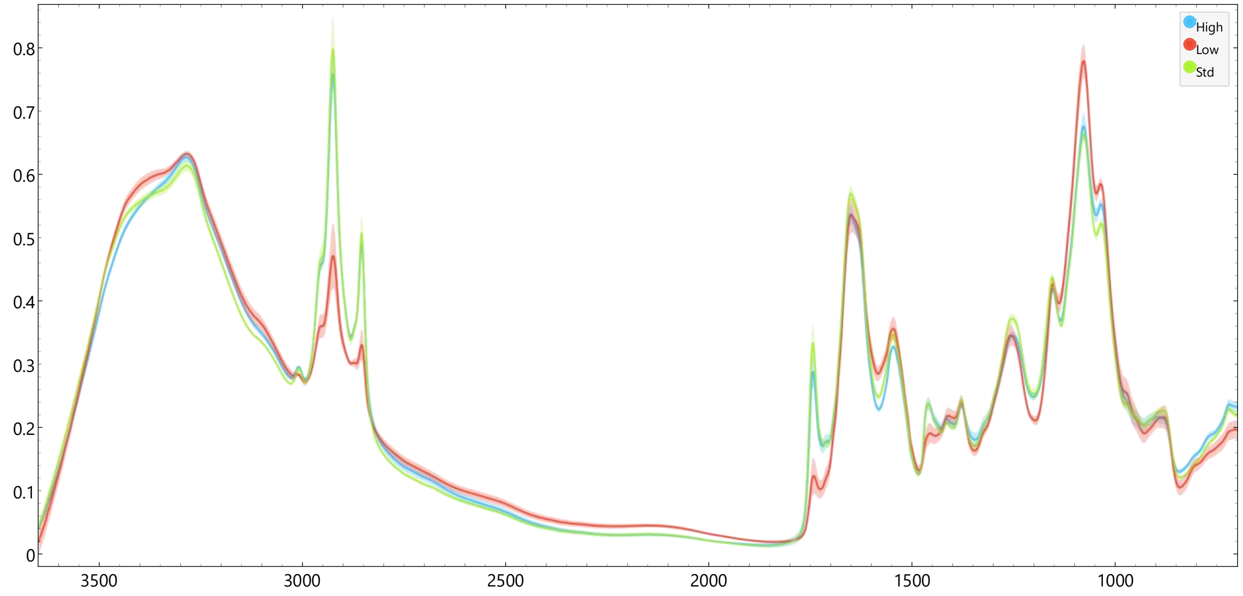


*Cunninghamella blakesleeana CCM F705*

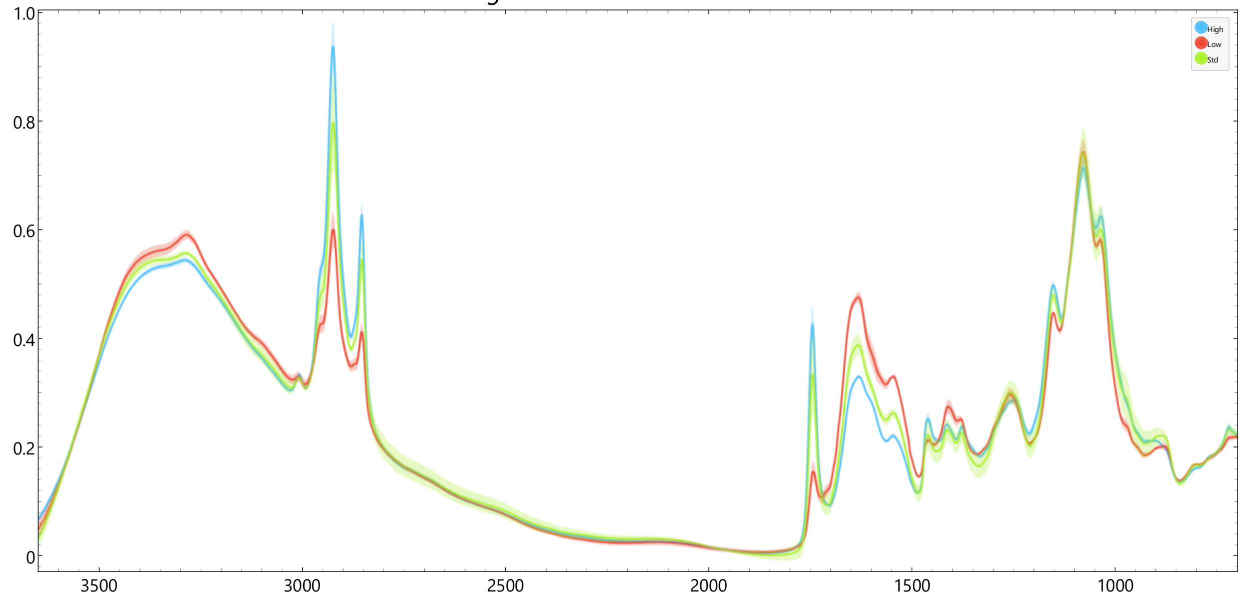

*Lichtheimia corymbifera CCM F8077*

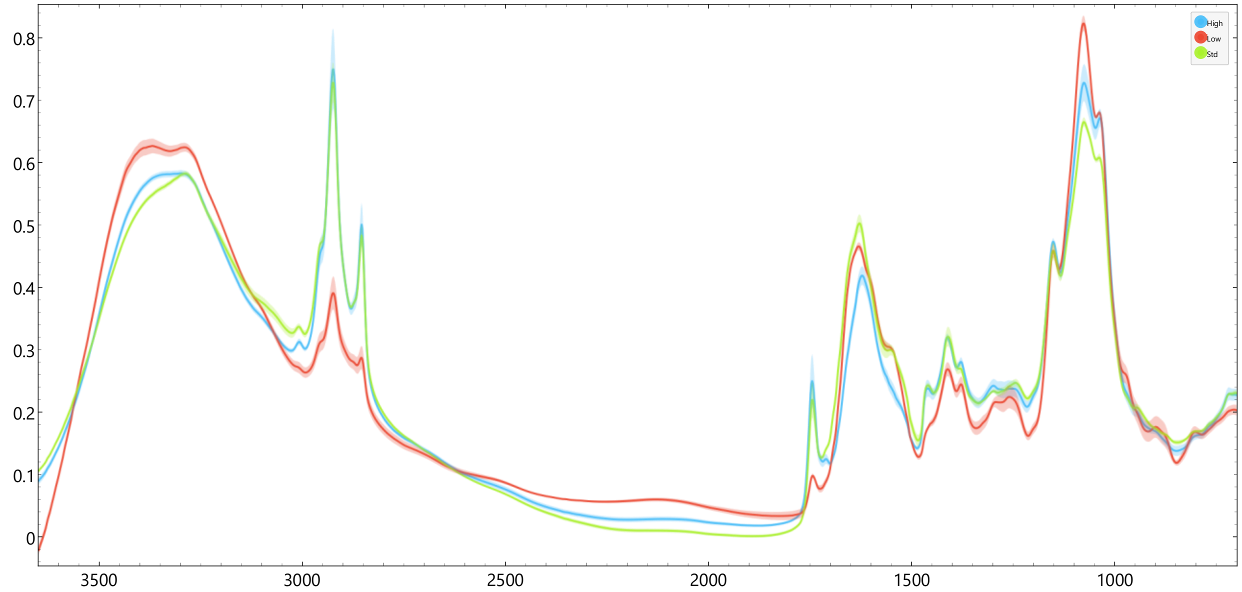


*Mortierella alpina ATCC 32222*

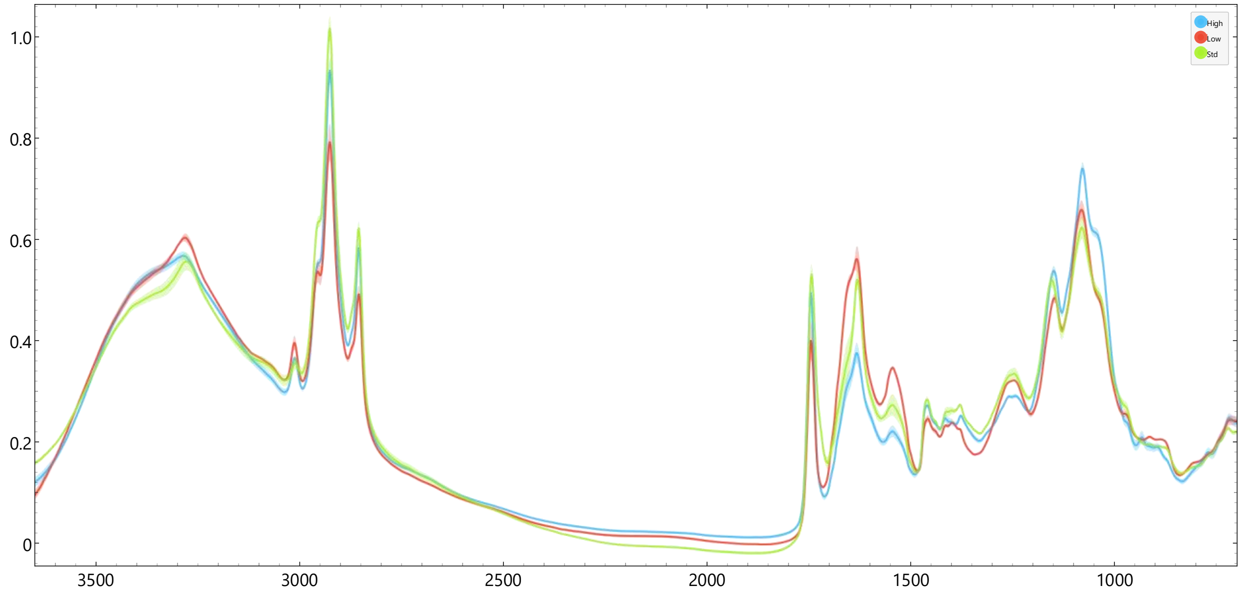

#### *Mucor circinelloides CCM F220*

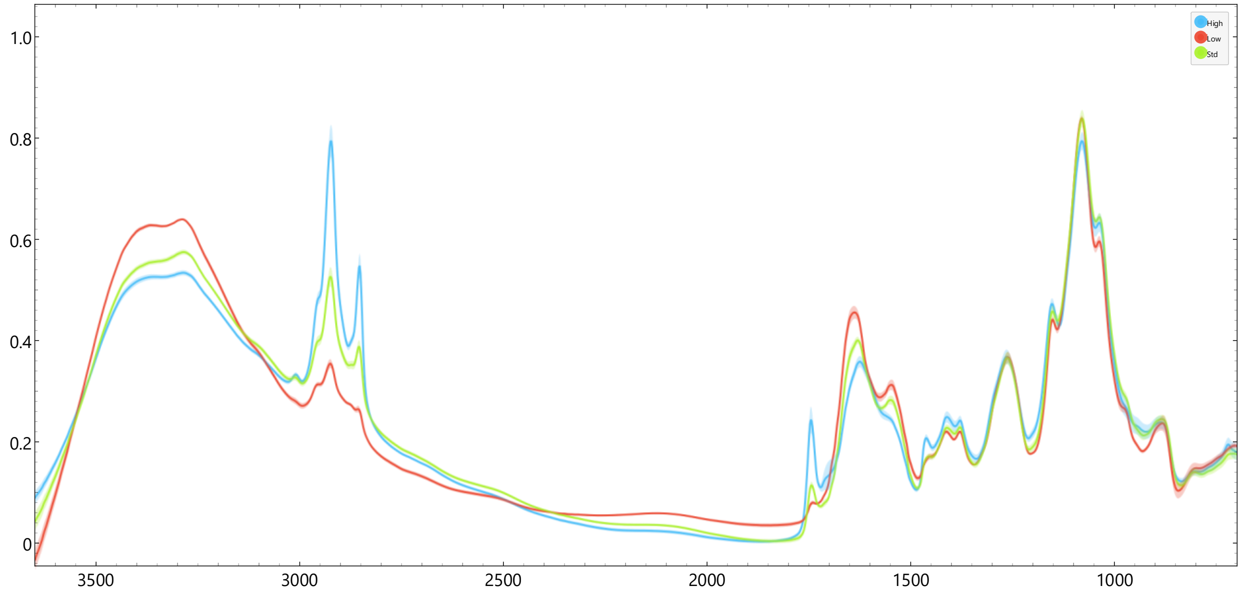


#### *Mucor circinelloides VI04473*

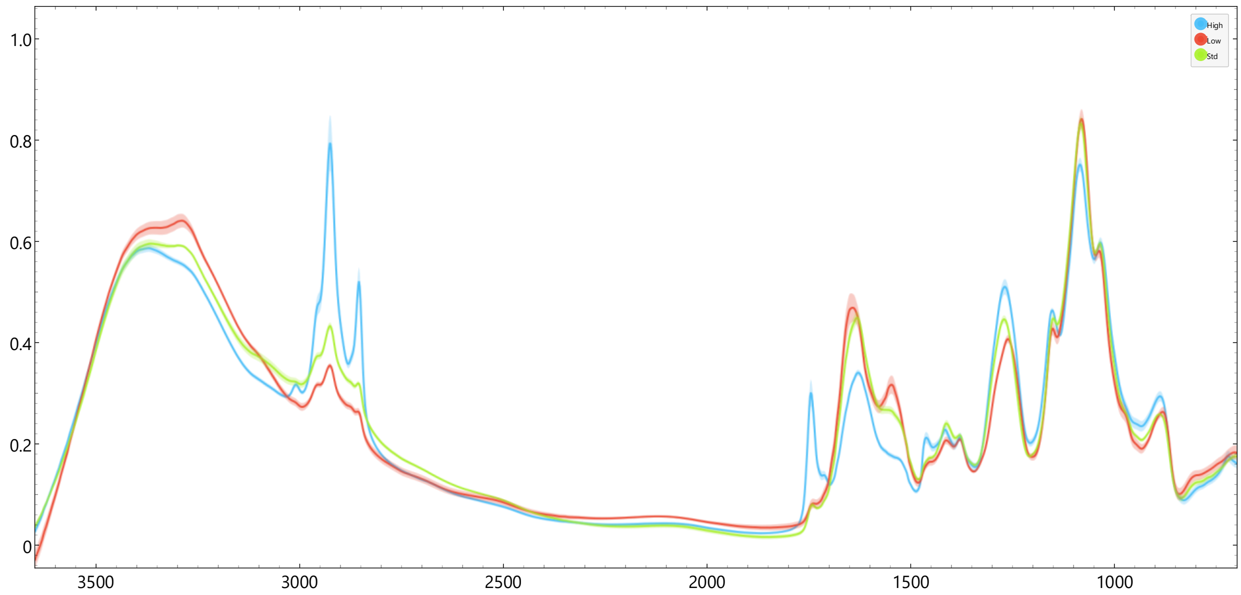

*Mortierella hyalina VKM F1629*

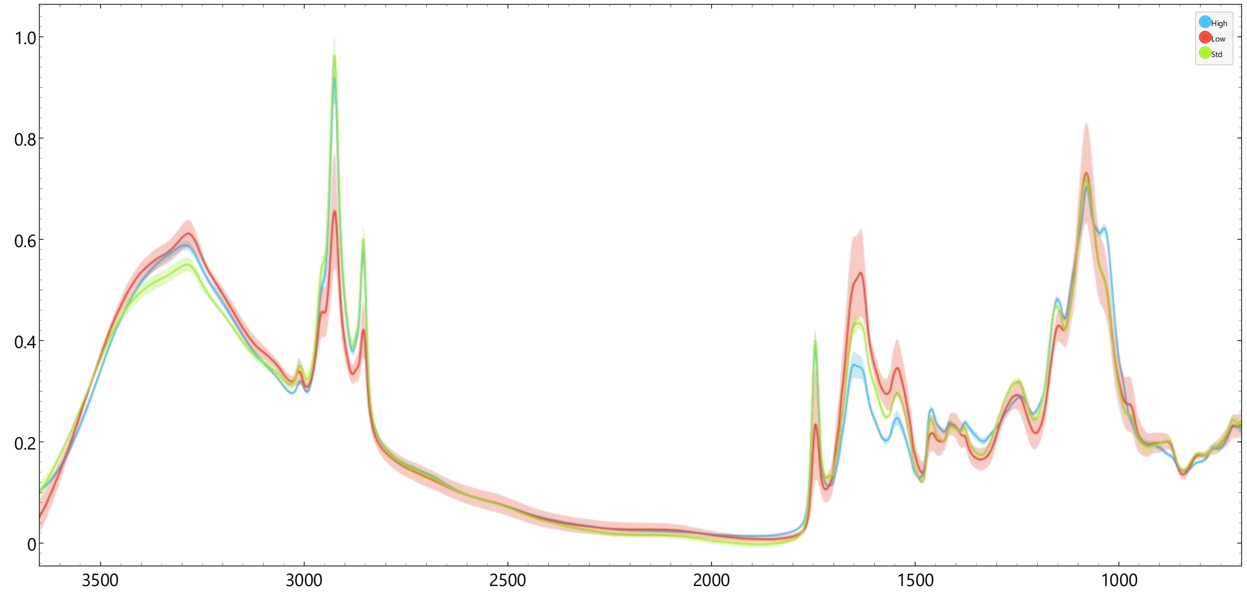


*Rhizopus stolonifer VKM F400*

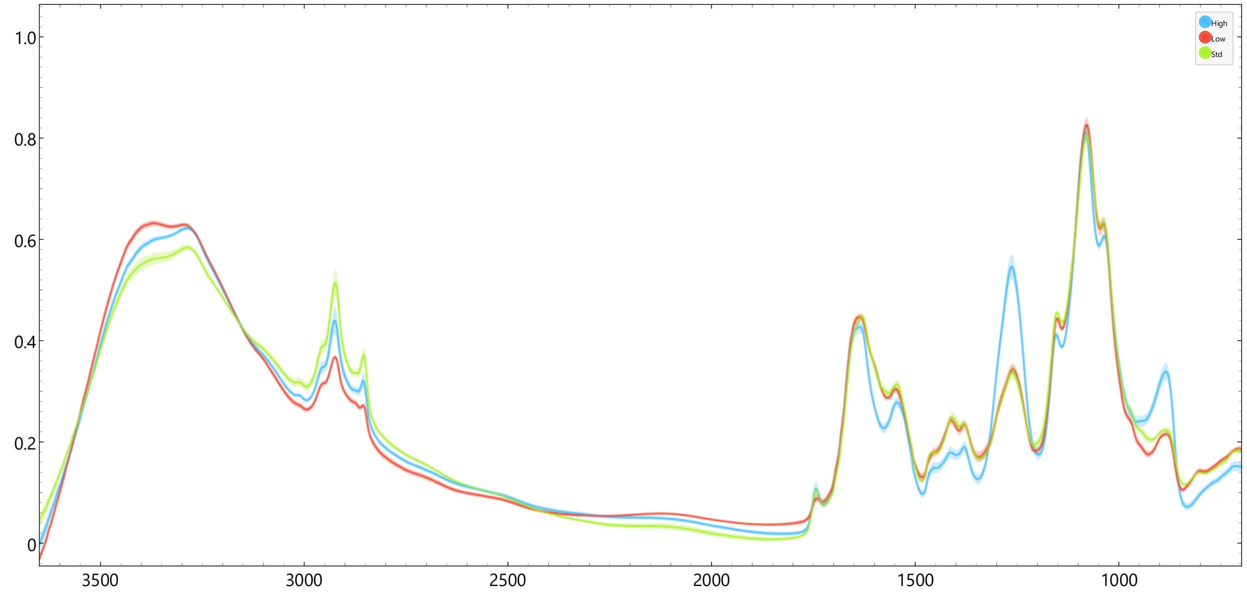

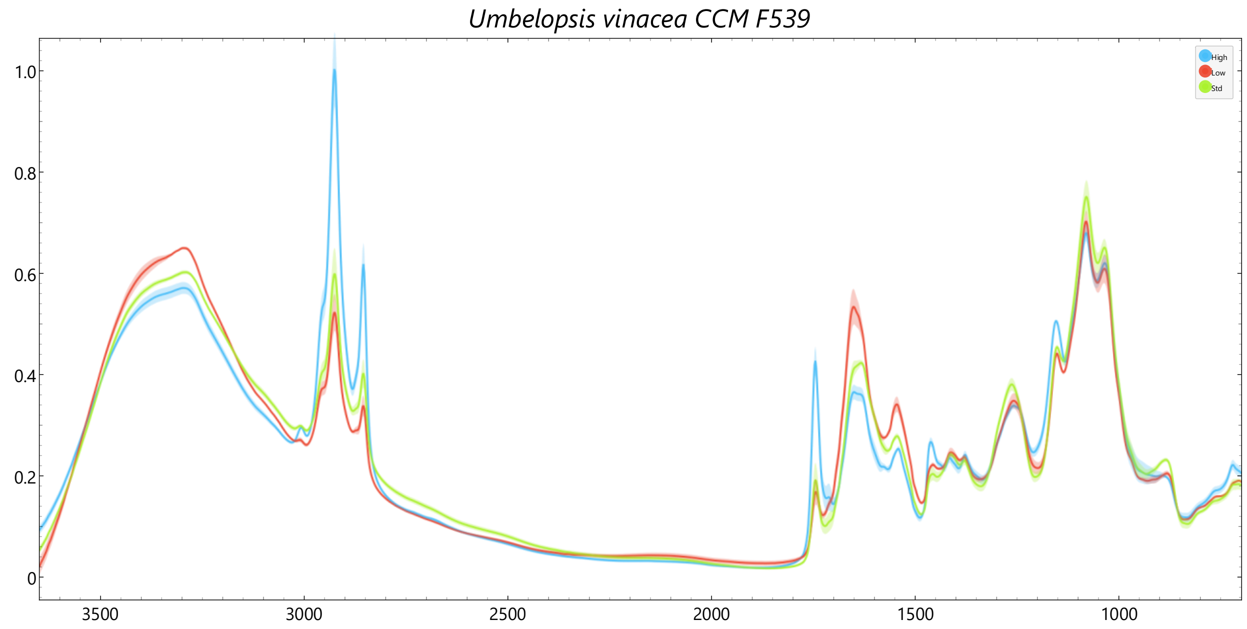

Umbelopsis vinacea CCM F539
High
Low
Std
1.0
0.8
0.6
0.4
0.2
0
3500
3000
2500
2000
1500
1000
