## Supplementary file 3 for "Carbon allocation strategies across Mucoromycota and the discovery of oxalic acid secretion in *Absidia*"

### Carbon allocation strategies across Mucoromycota and the discovery of oxalic acid secretion in *Absidia*

Cristian Bolaño Losada[1], Boris Zimmermann[1], Dana Byrtusova[1], Sven Andreas Högfeldt[1], Svein Jarle Horn[2], Achim Kohler[1], Volha Shapaval[1]

[1] Faculty of Science and Technology, Norwegian University of Life Sciences (NMBU), 1432 Ås, Norway

[2] Faculty of Chemistry, Biotechnology and Food Science, Norwegian University of Life Sciences (NMBU), 1432 Ås, Norway.

#### Index

#### Introduction and summary of results

This additional file contains the GC-MS results from BSTFA derivatized diethylether extracts from the supernatants of fermentations from all the strains used in this study. This procedure was done to determine presence of short-chain carboxylic acids and other extracellular metabolites.
The following table summarizes the main metabolites found for each sample.

Table S1. List of strains used in the study and short-chain carboxylic acids detected by GC-MS from the supernatants after incubation in high C/N medium. Cells marked as * correspond to glycerol in *M. cicinelloides* strains, which was detected by ion exchange chromatography in lower amounts but not by diethylether extraction and GC-MS since it is not an optimal procedure for this type of molecule. The detection of glycerol in *R. stolonifer* was likely due to the high concentrations of its supernatants.

| **Species** | **Culture collection code** | **Succinic acid** | **Lactic acid** | **Oxalic acid** | **Fumaric acid** | **Glycerol** |
|---|---|---|---|---|---|---|
| *Absidia coerulea* | NRRL 1322 | | | ■ | | |
| *Absidia cuneospora* | NRRL 2632 | | ■ | ■ | | |
| *Absidia glauca* | CCM F451 | | ■ | ■ | | |
| *Absidia glauca* | CCM F450 | | ■ | ■ | | |
| *Absidia glauca* | CCM F444 | | | ■ | | |
| *Absidia glauca* | NRRL 2799 | | ■ | ■ | | |
| *Absidia heterospora* | NRRL 2800 | | | ■ | | |
| *Absidia pseudocylindrospora* | CCM F52 | | ■ | ■ | | |
| *Absidia psychrophilia* | NRRL 3044 | | ■ | ■ | | |
| *Cunninghamella blakesleeana* | CCM F705 | | ■ | | | |
| *Lichtheimia blakesleeana* | NRRL 2696 | | | | | |
| *Lichtheimia corymbifera* | CCM F8077 | | | | | |
| *Mortierella alpina* | ATCC 32222 | | ■ | | | |
| *Mortierella hyalina* | VKM F1629 | | ■ | | | |
| *Mucor circinelloides* | CCM F220 | | ■ | | | * |
| *Mucor circinelloides* | VI04473 | | ■ | | | * |
| *Rhizopus stolonifer* | VKM F400 | ■ | ■ | | ■ | ■ |
| *Umbelopsis vinacea* | CCM F539 | | ■ | | | |

The following pages display the individual results for each strain. In each sample section, the GC-MS chromatogram is shown first, followed by the mass spectrum of each metabolite peak and its corresponding match/probability with compounds in the *mainlib* library. Peaks with low match or probability values were not considered.

A sample of pure oxalic acid BSTFA derivatized was added to confirm retention time and spectrum.

#### Oxalic acid standard solution BSTFA derivatized

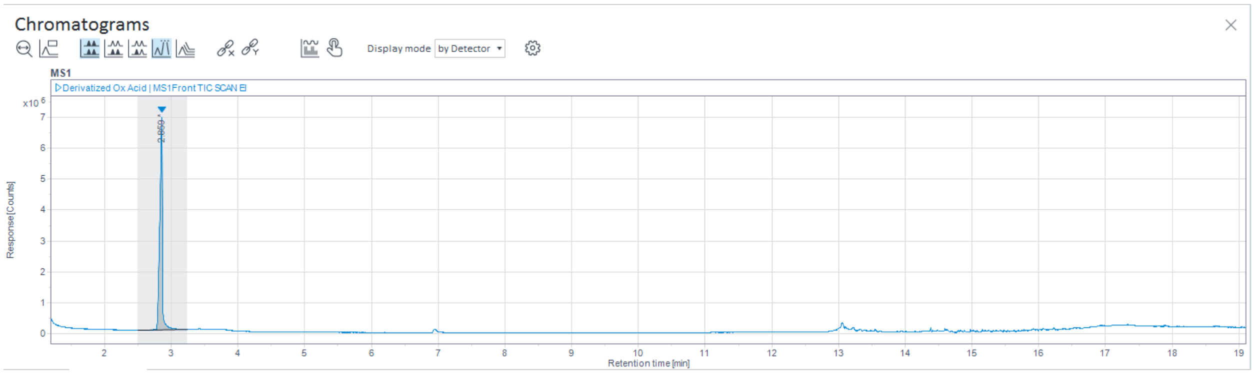

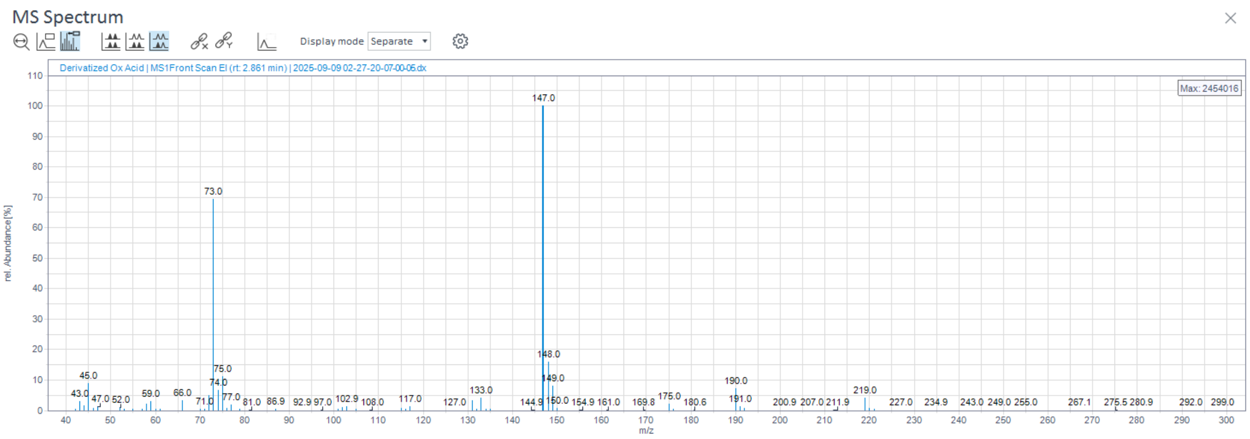

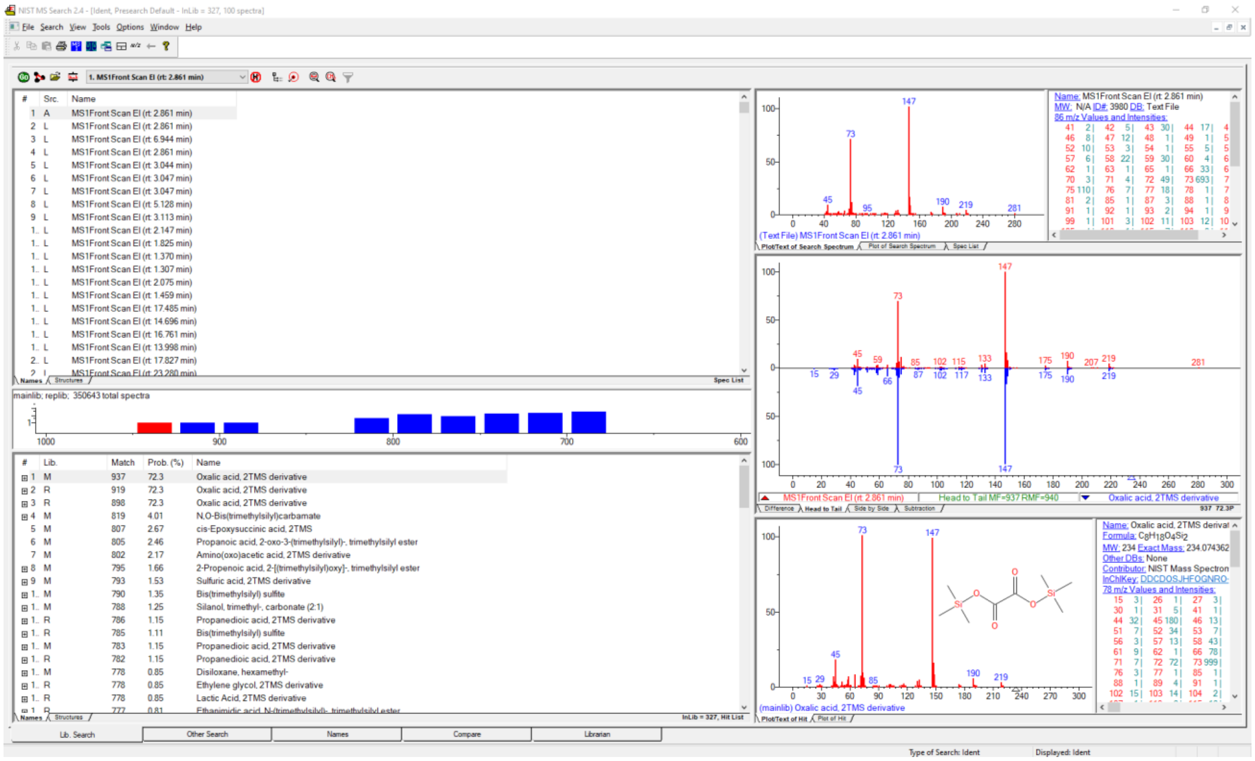

#### *Absidia coerulea* NRRL 1322

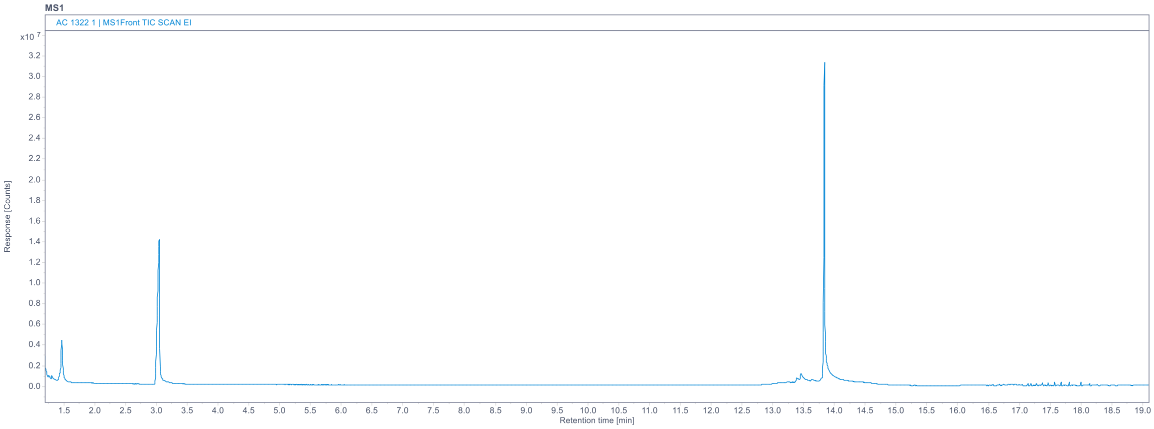

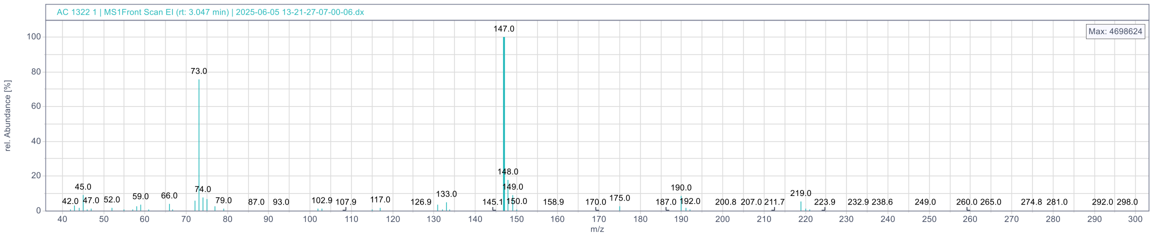

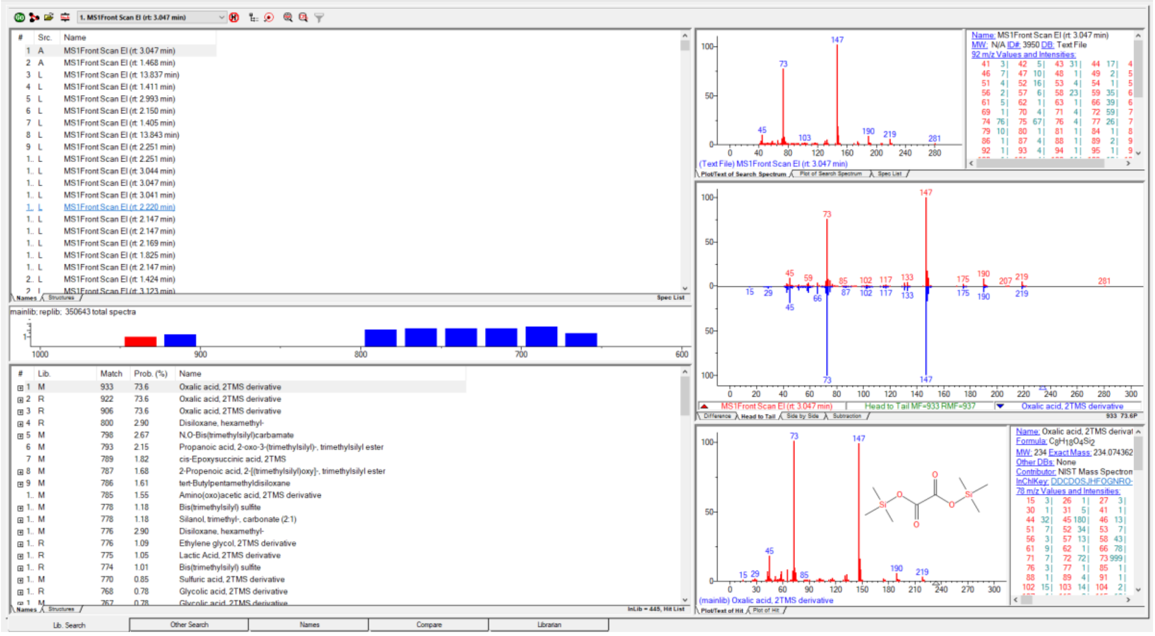

#### *Absidia cunespora* NRRL 2632

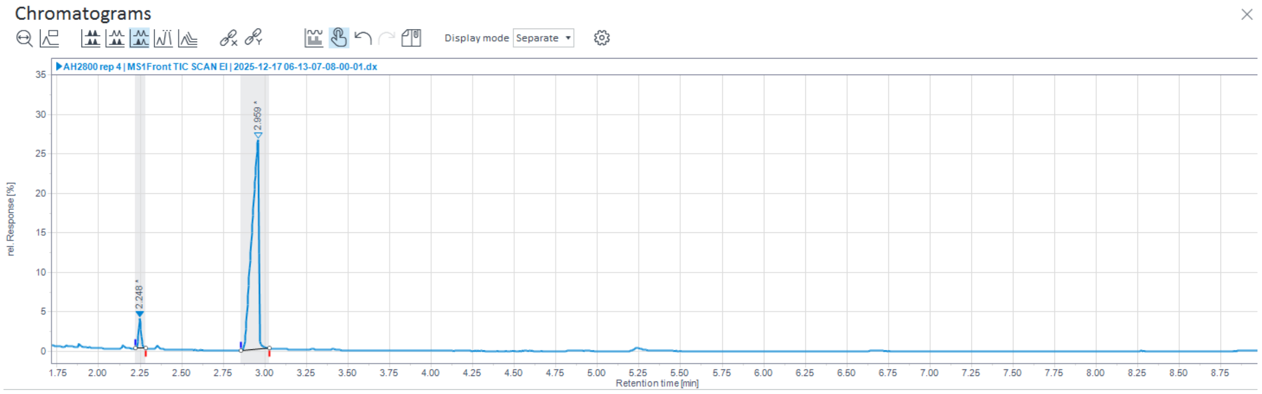

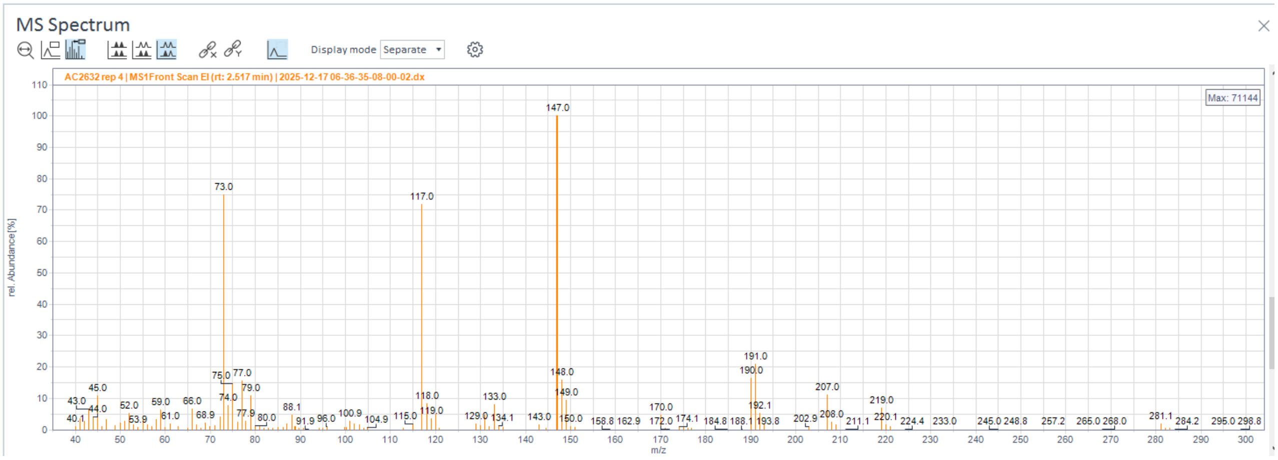

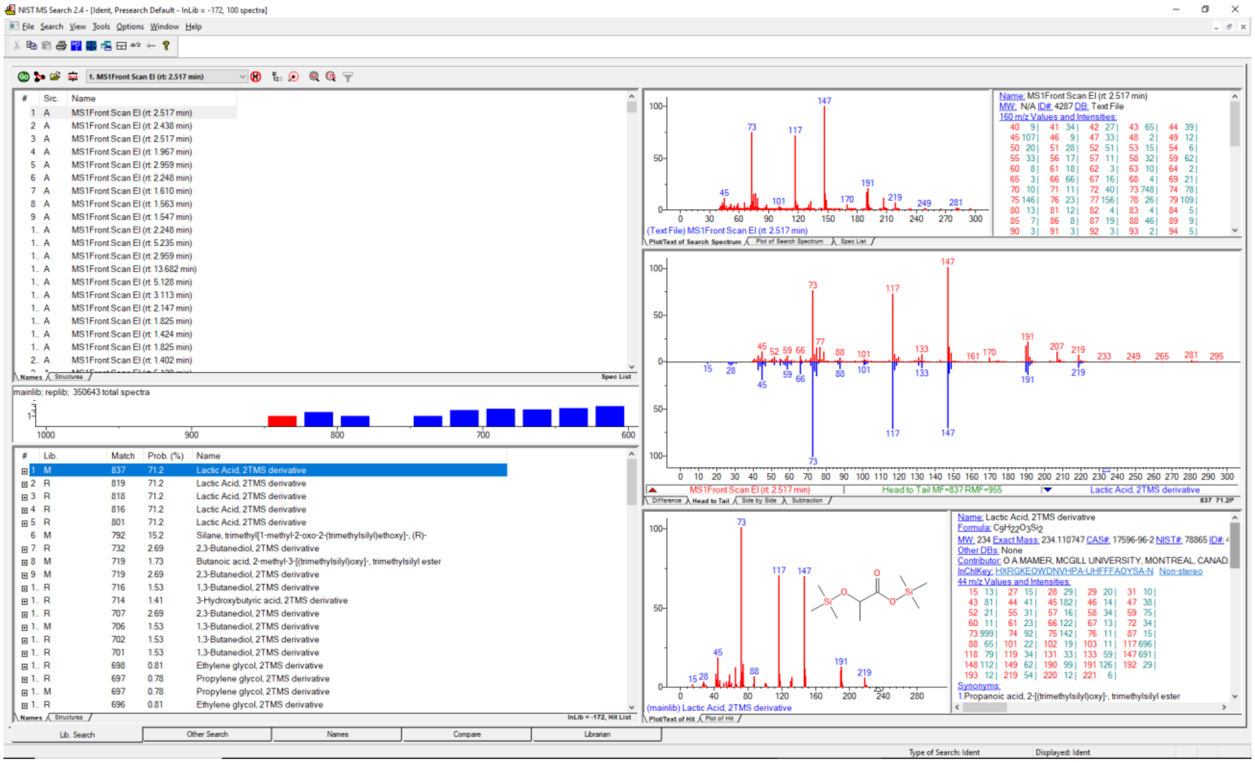

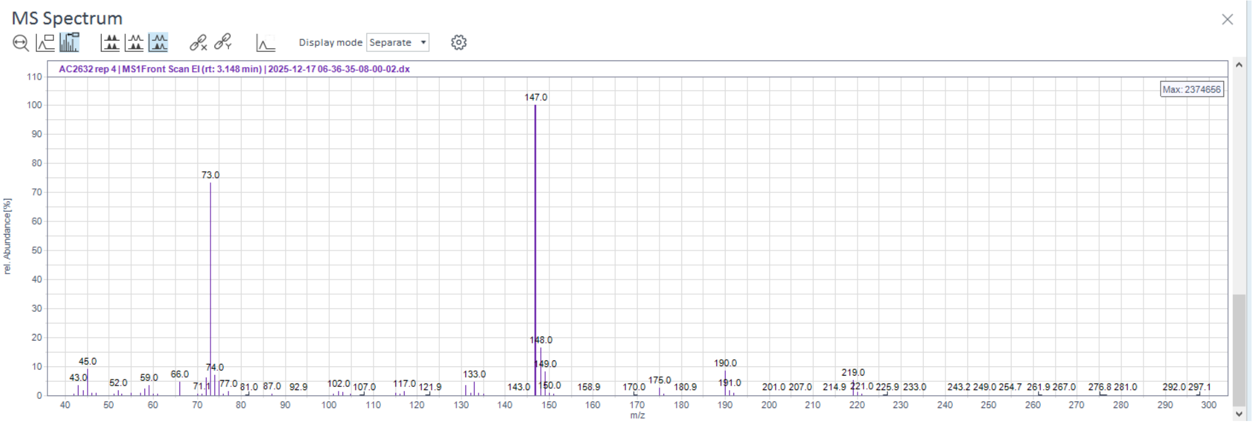
MS Spectrum
Display mode Separate
AC2632 rep 4 | MS1Front Scan EI (rt: 3.148 min) | 2025-12-17 06-36-35-08-00-02.dx
Max: 2374656
rel. Abundance [%]
m/z
147.0
73.0
148.0
45.0
74.0
149.0
190.0
219.0
133.0

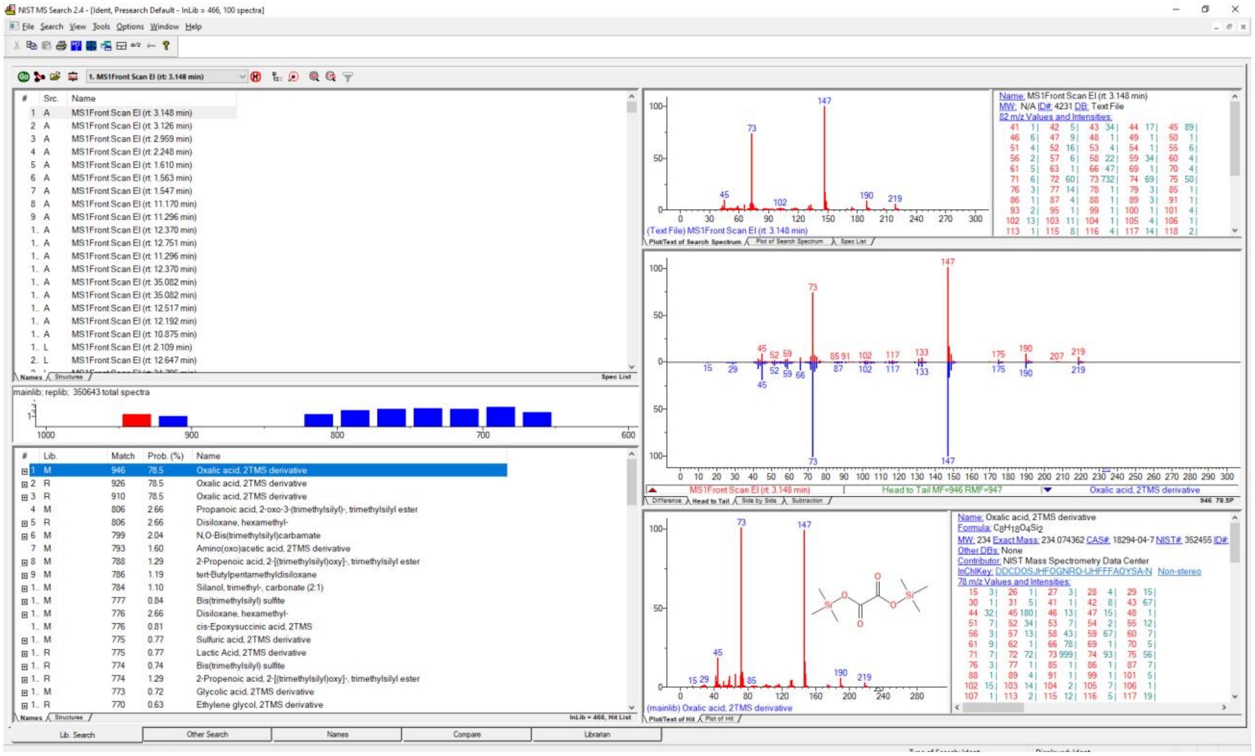
NIST MS Search 2.4 - [Ident, Presearch Default - InLib = 466, 100 spectra]
File Search View Tools Options Window Help
1. MS1Front Scan EI (rt: 3.148 min)
mainlib; replib; 350643 total spectra
Name: MS1Front Scan EI (rt: 3.148 min)
MW: N/A ID#: 4231 DB: Text File
82 m/z Values and Intensities:
(Text File) MS1Front Scan EI (rt: 3.148 min)
Head to Tail MF=946 RMF=947
Oxalic acid, 2TMS derivative
Oxalic acid, 2TMS derivative
Propanoic acid, 2-oxo-3-(trimethylsilyl)-, trimethylsilyl ester
Disiloxane, hexamethyl-
N,O-Bis(trimethylsilyl)carbamate
Amino(oxo)acetic acid, 2TMS derivative
2-Propenoic acid, 2-[(trimethylsilyl)oxy]-, trimethylsilyl ester
tert-Butylpentamethyldisiloxane
Silanol, trimethyl-, carbonate (2:1)
Bis(trimethylsilyl) sulfite
cis-Epoxysuccinic acid, 2TMS
Sulfuric acid, 2TMS derivative
Lactic Acid, 2TMS derivative
Glycolic acid, 2TMS derivative
Ethylene glycol, 2TMS derivative
Name: Oxalic acid, 2TMS derivative
MW: 234 Exact Mass: 234.074362 CAS#: 18294-04-7 NIST#: 352455
Other DBs: None
Contributor: NIST Mass Spectrometry Data Center
InChIKey: DDCDOSJHFOGNRO-UHFFFAOYSA-N Non-stereo
78 m/z Values and Intensities:
(mainlib) Oxalic acid, 2TMS derivative
Lib. Search
Other Search
Names
Compare
Librarian

#### *Absidia glauca* CCM F451

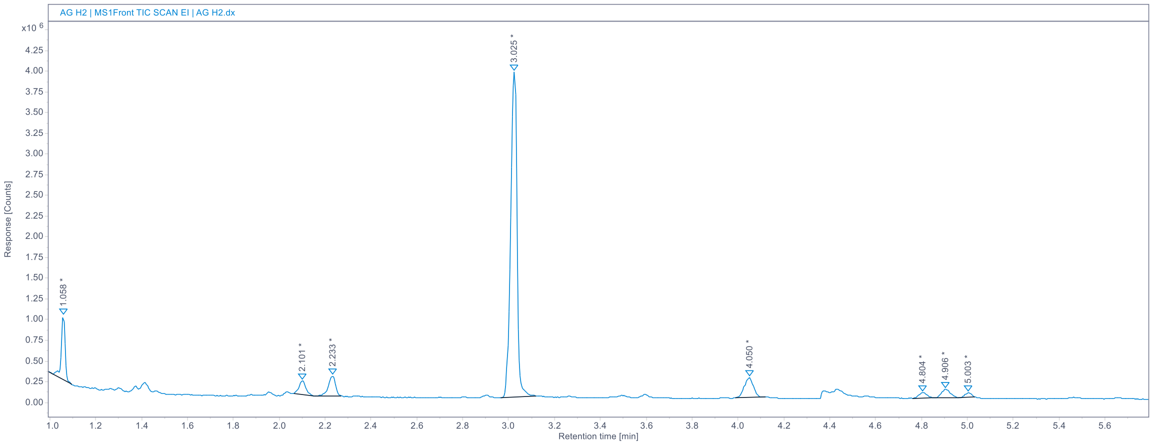

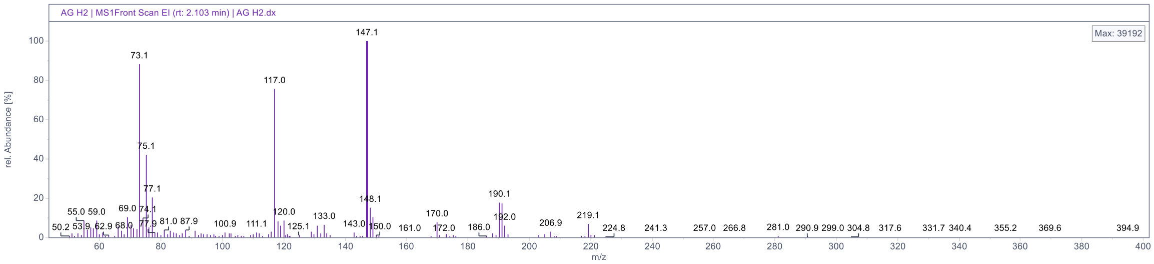

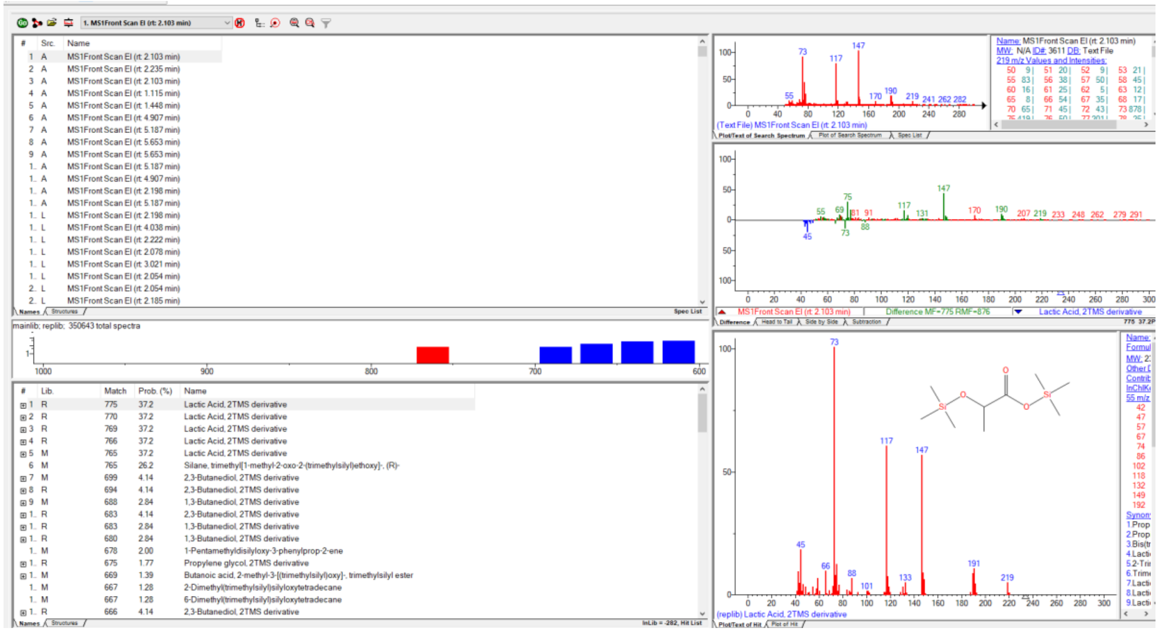

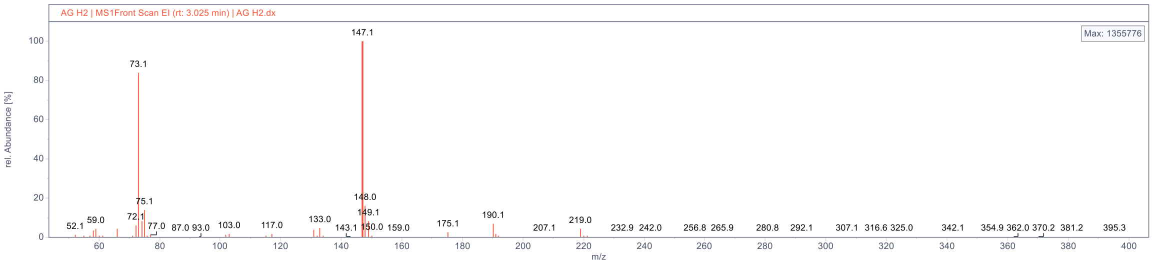

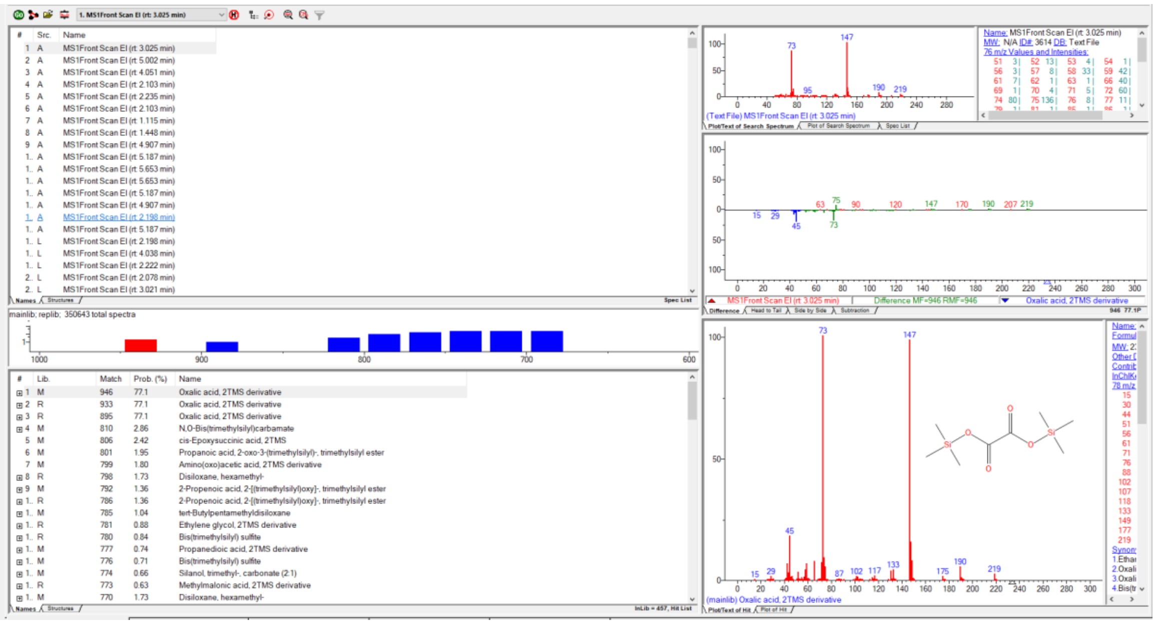

#### *Absidia glauca* CCM F450

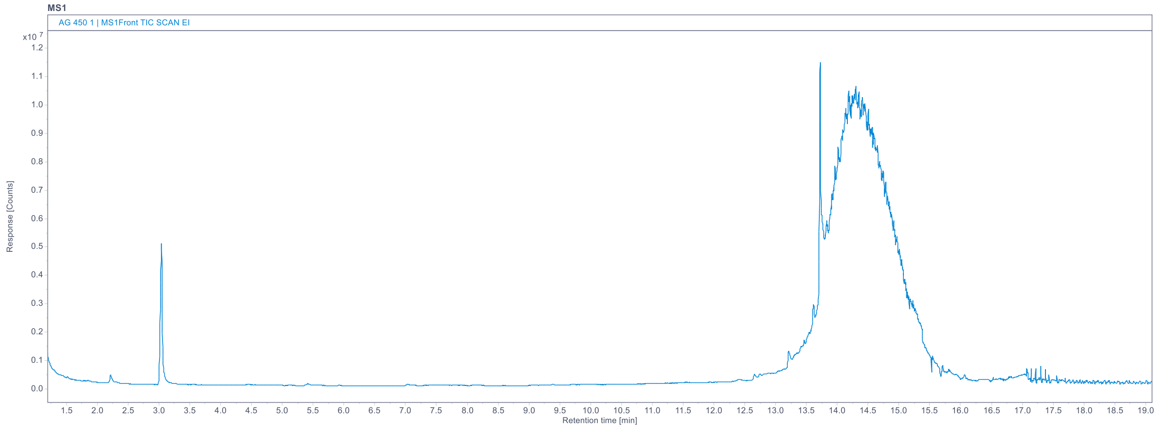

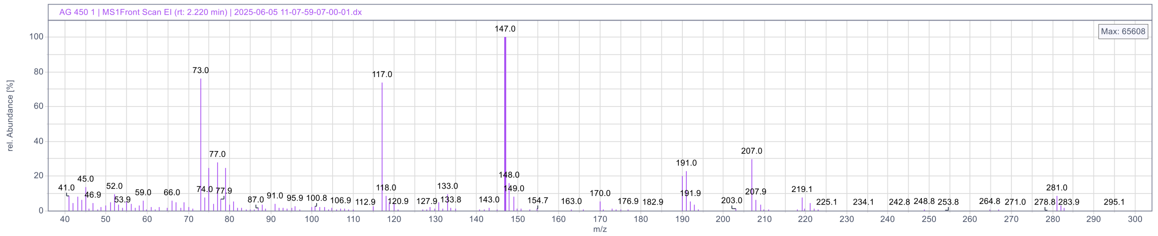

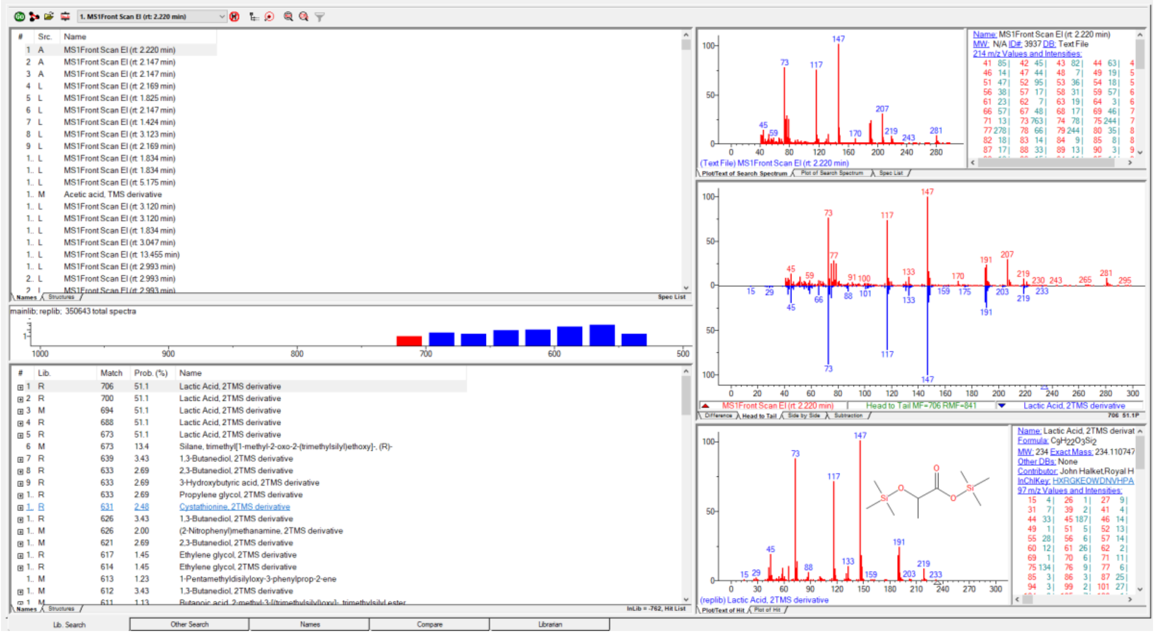

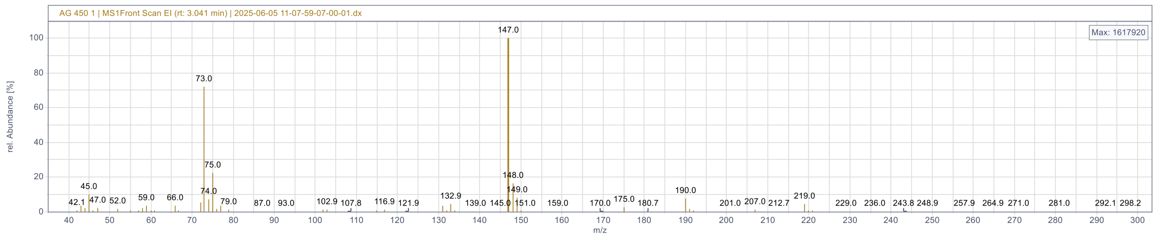

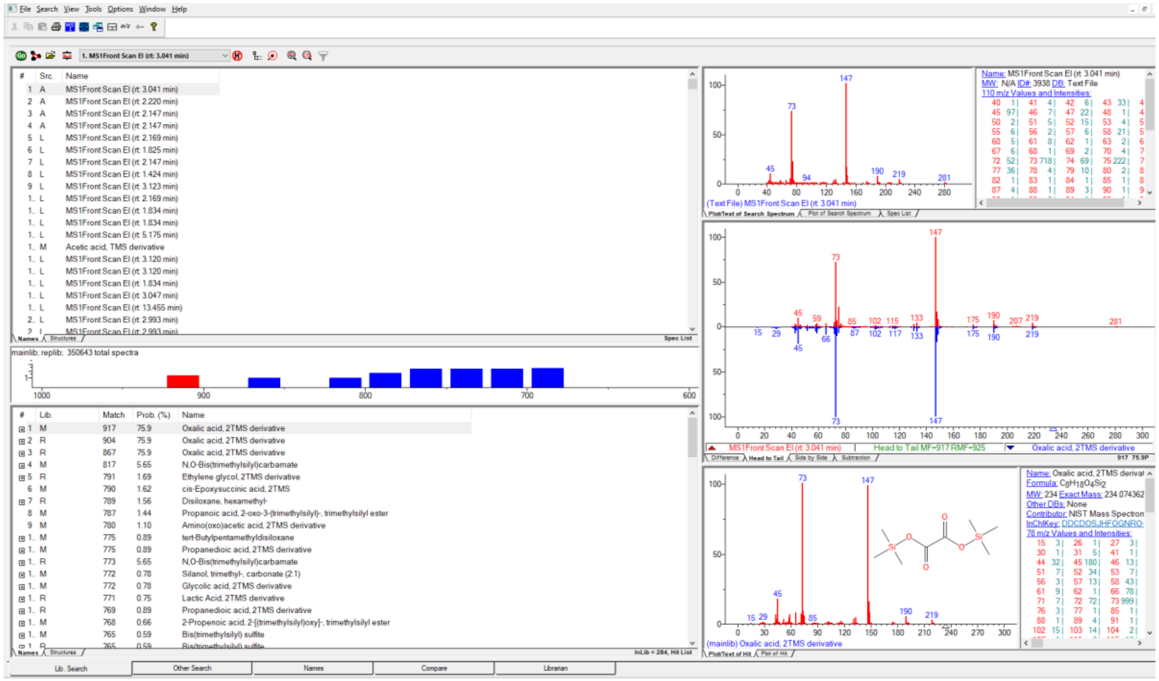

#### *Absidia glauca* CCM F444

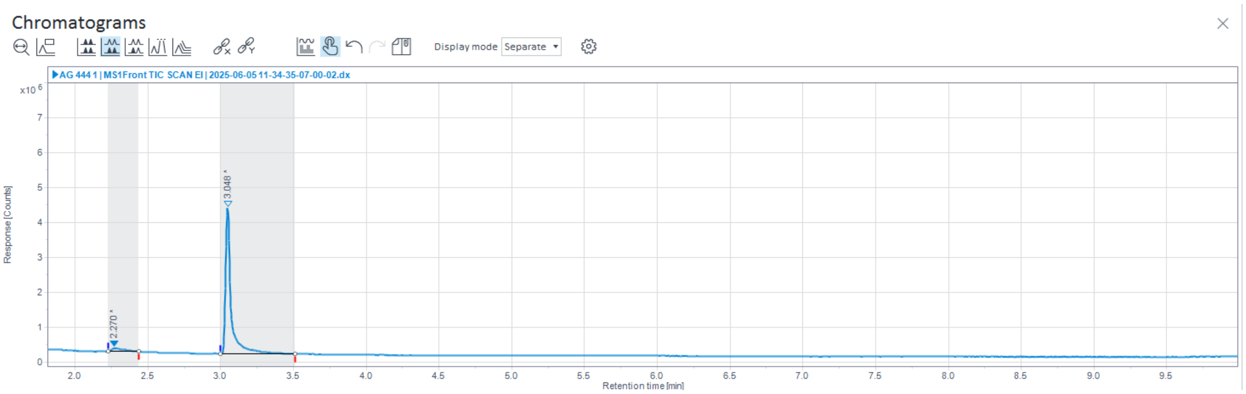

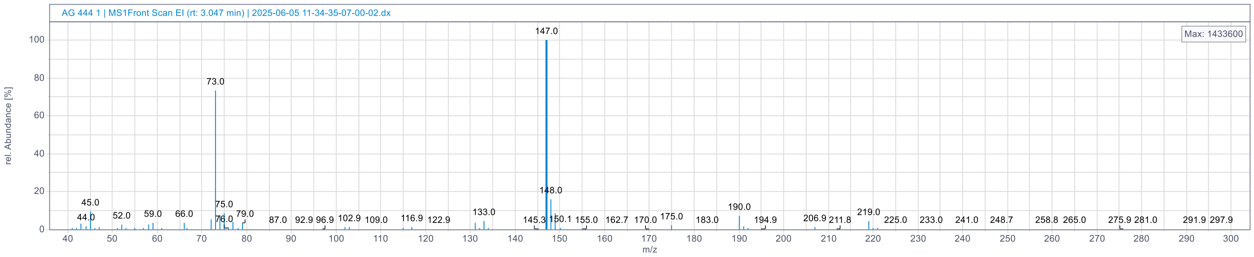

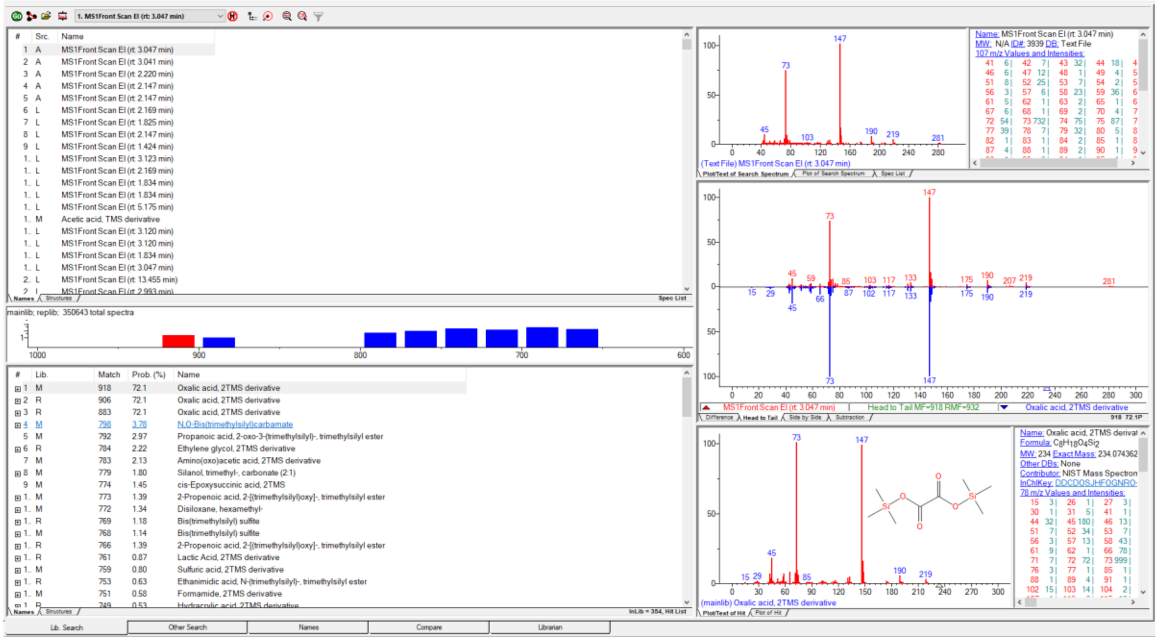

#### *Absidia glauca* NRRL 2799

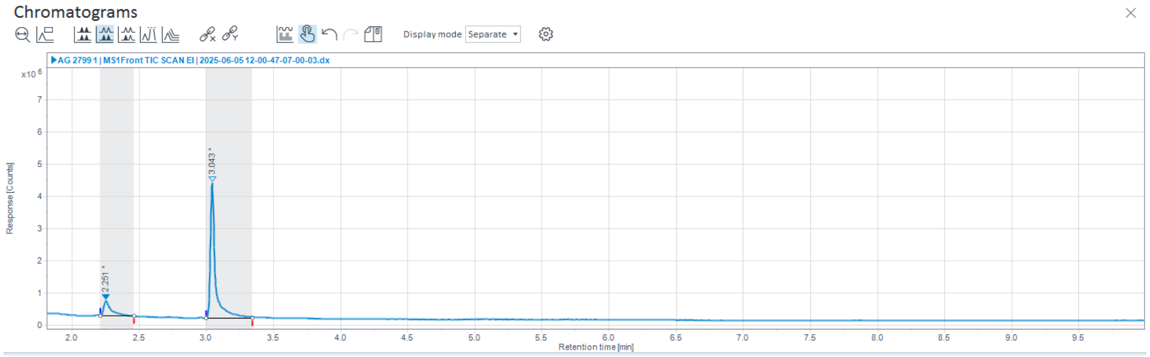

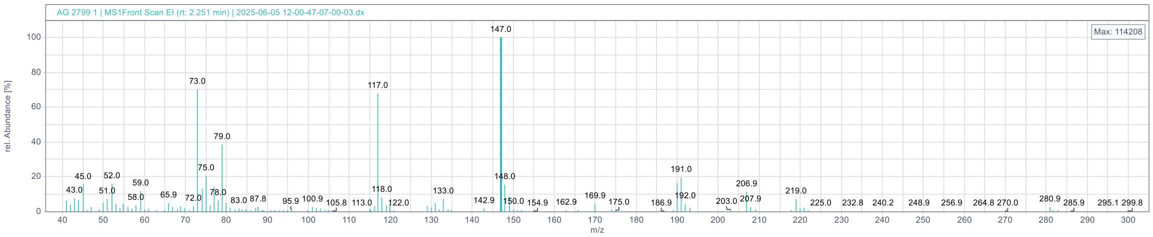

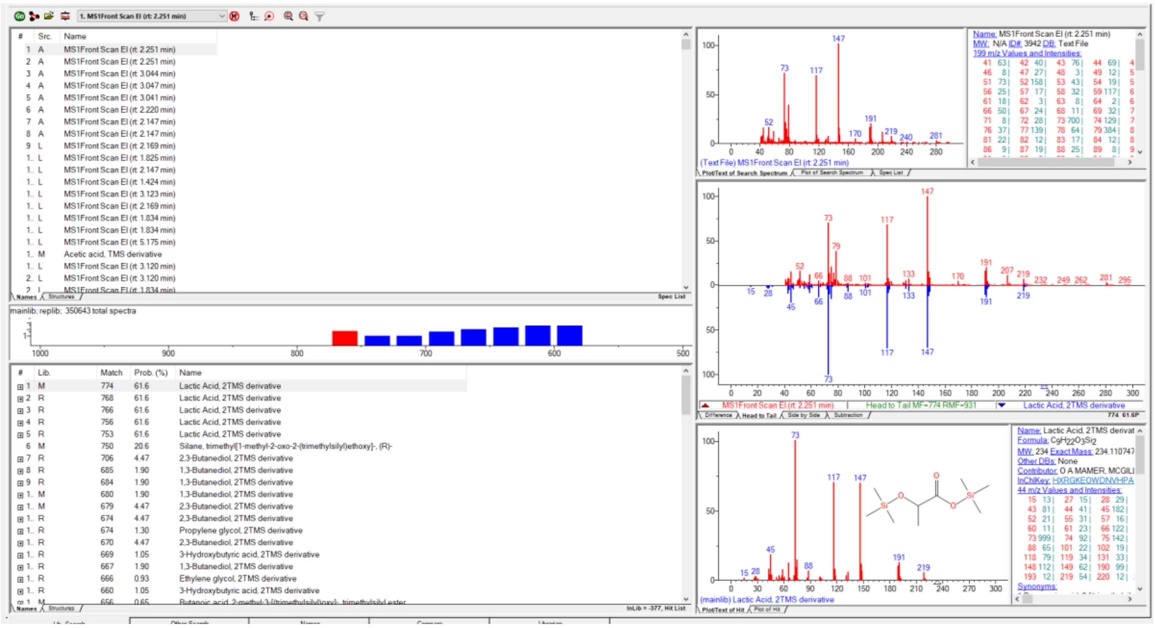

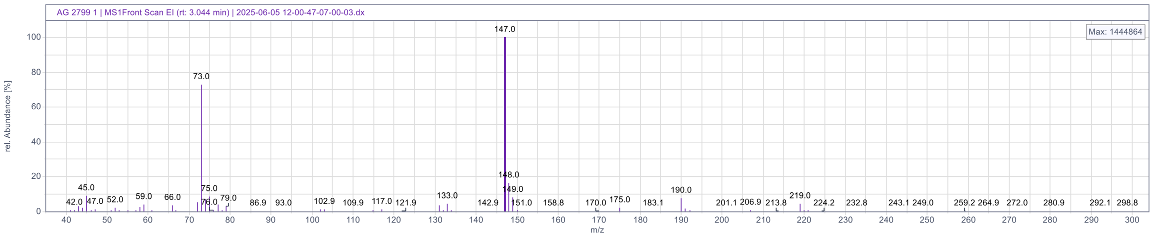

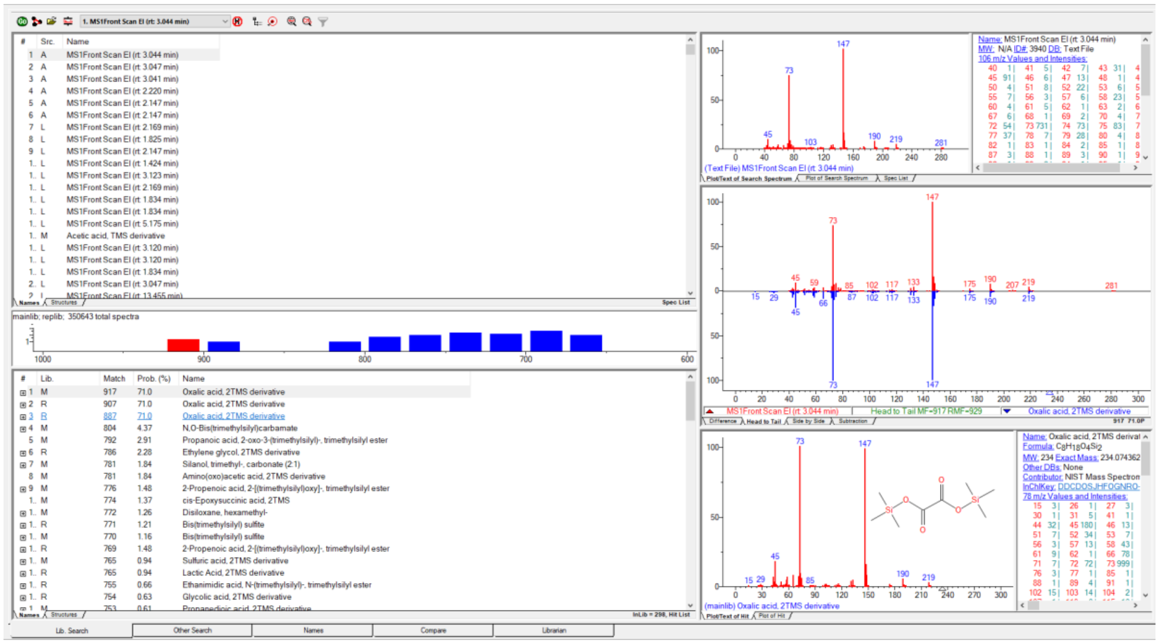

#### *Absidia heterospora* NRRL 2800

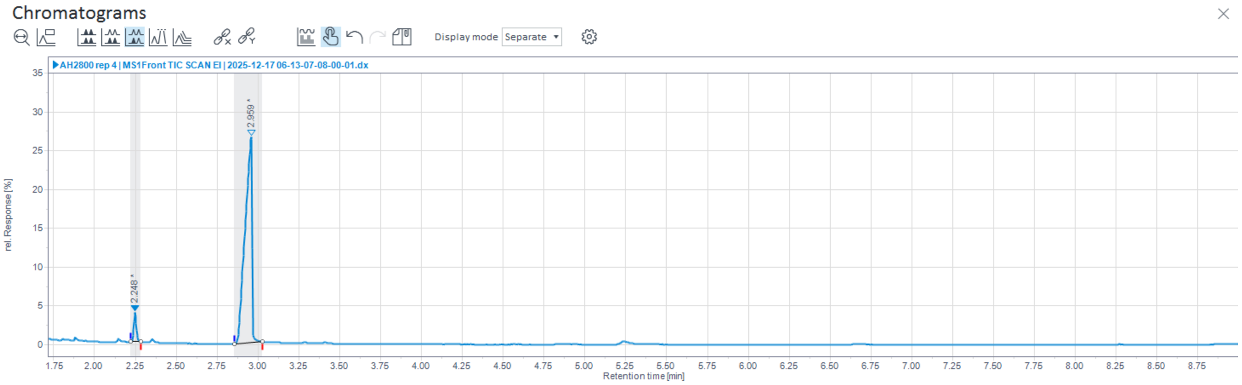

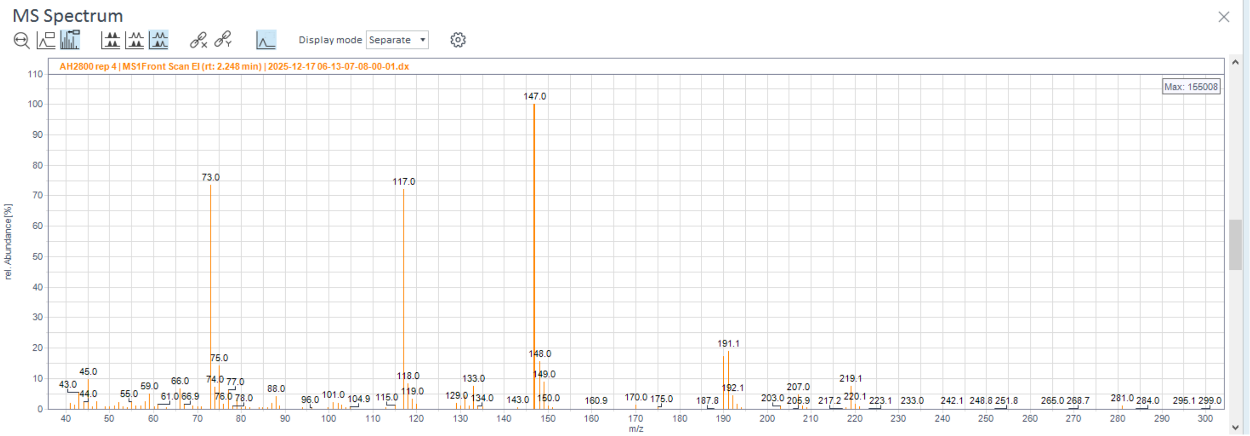

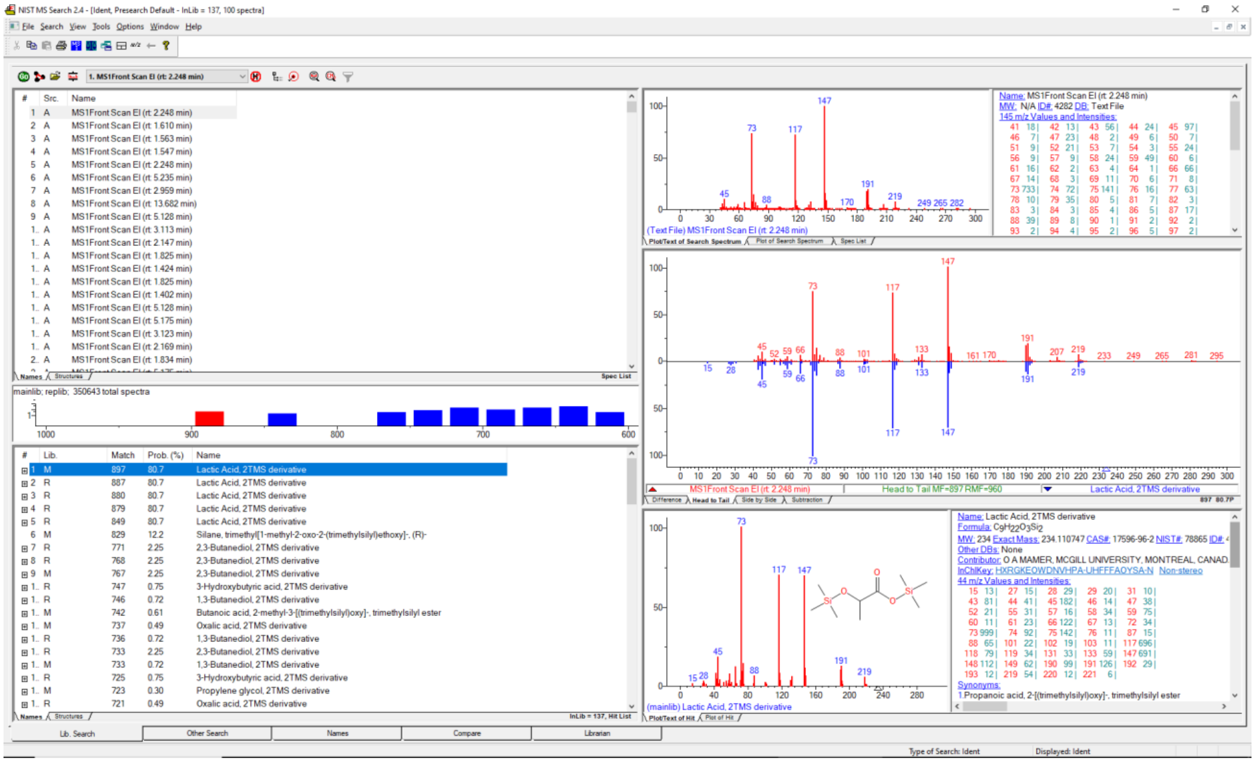

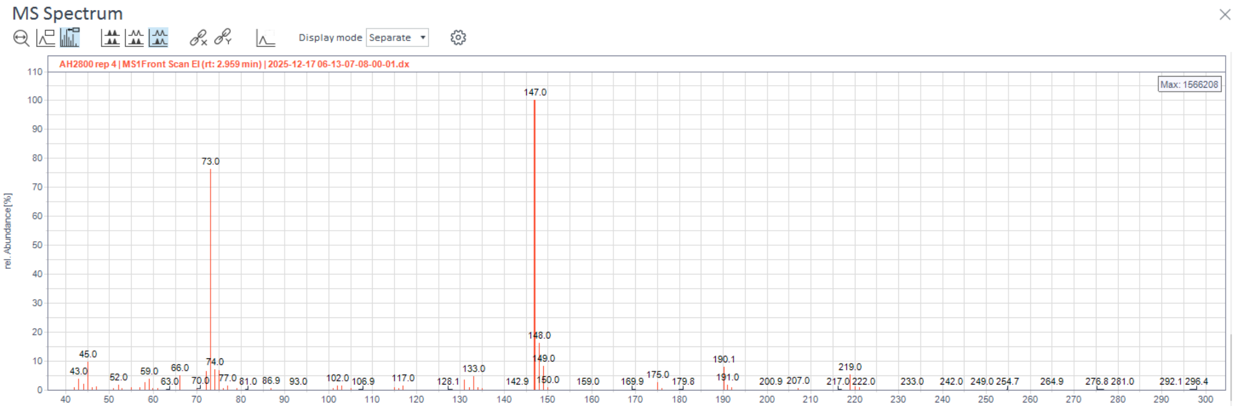
MS Spectrum
Display mode Separate
AH2800 rep 4 | MS1Front Scan EI (rt: 2.959 min) | 2025-12-17 06-13-07-08-00-01.dx
Max: 1566208
rel. Abundance [%]
147.0
73.0
148.0
149.0
45.0
190.1
219.0
133.0

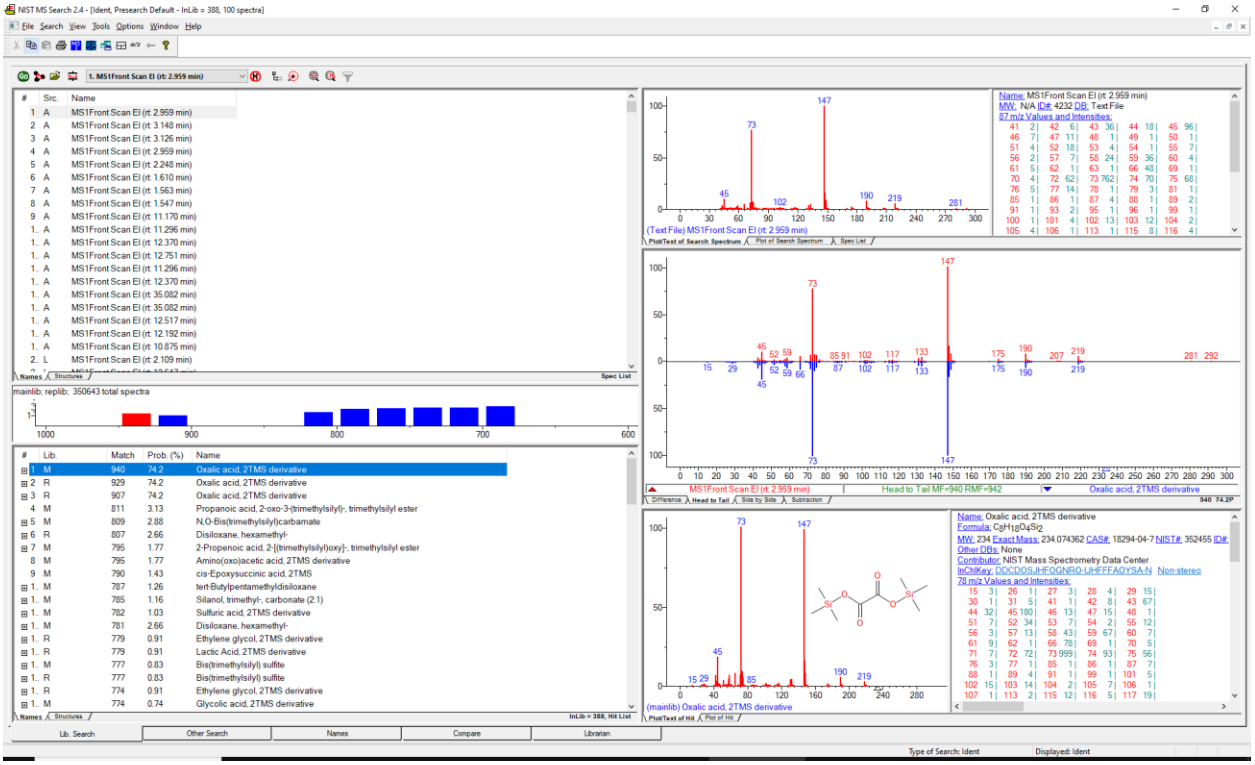
NIST MS Search 2.4 - [Ident, Presearch Default - InLib = 388, 100 spectra]
1. MS1Front Scan EI (rt: 2.959 min)
mainlib; replib; 350643 total spectra
Oxalic acid, 2TMS derivative
Propanoic acid, 2-oxo-3-(trimethylsilyl)-, trimethylsilyl ester
N,O-Bis(trimethylsilyl)carbamate
Disiloxane, hexamethyl-
2-Propenoic acid, 2-[(trimethylsilyl)oxy]-, trimethylsilyl ester
Amino(oxo)acetic acid, 2TMS derivative
cis-Epoxysuccinic acid, 2TMS
tert-Butylpentamethyldisiloxane
Silanol, trimethyl-, carbonate (2:1)
Sulfuric acid, 2TMS derivative
Ethylene glycol, 2TMS derivative
Lactic Acid, 2TMS derivative
Bis(trimethylsilyl) sulfite
Glycolic acid, 2TMS derivative
Head to Tail MF=940 RMF=942
Name: Oxalic acid, 2TMS derivative
Formula: C8H18O4Si2
MW: 234 Exact Mass: 234.074362 CAS#: 18294-04-7 NIST#: 352455
Contributor: NIST Mass Spectrometry Data Center
InChIKey: DDCDOSJHFOGNRO-UHFFFAOYSA-N

#### *Absidia pseudocylindrospora* CCM F52

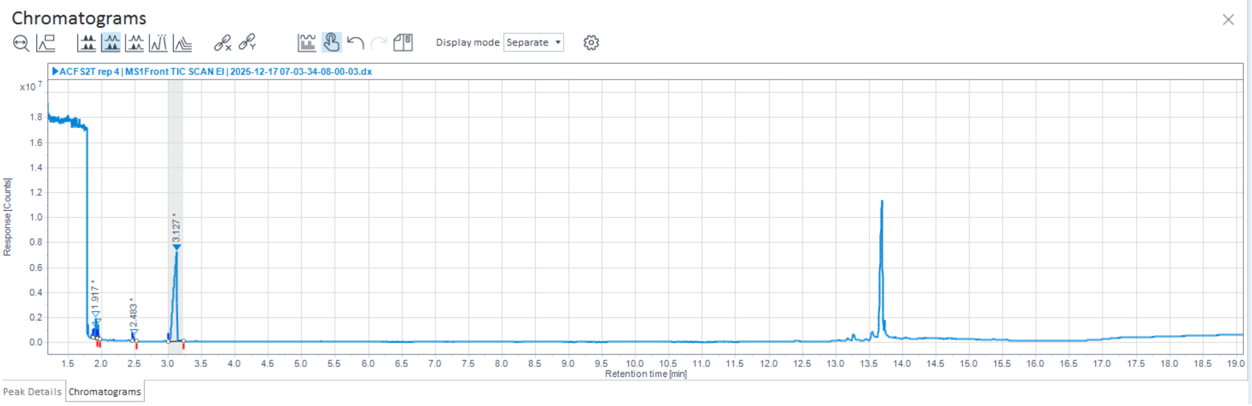

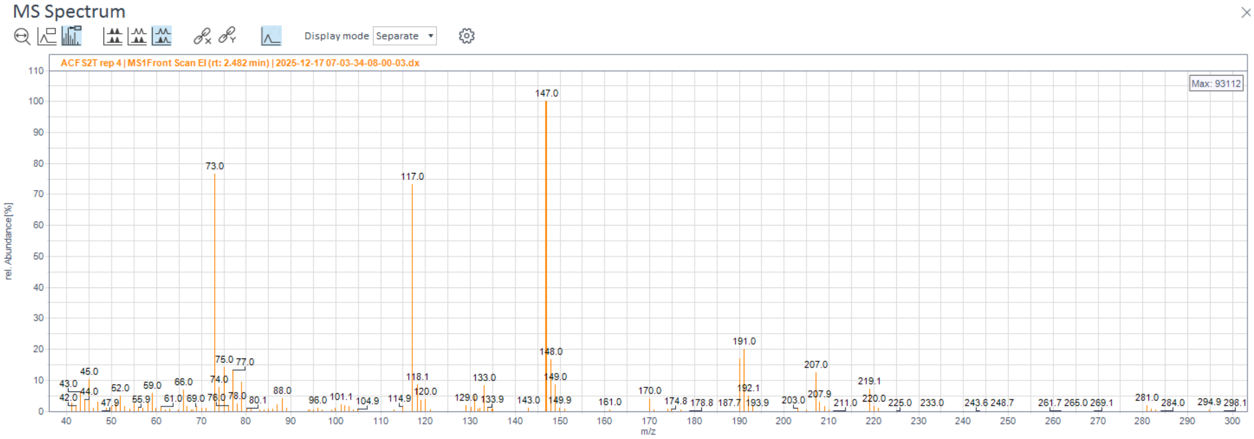

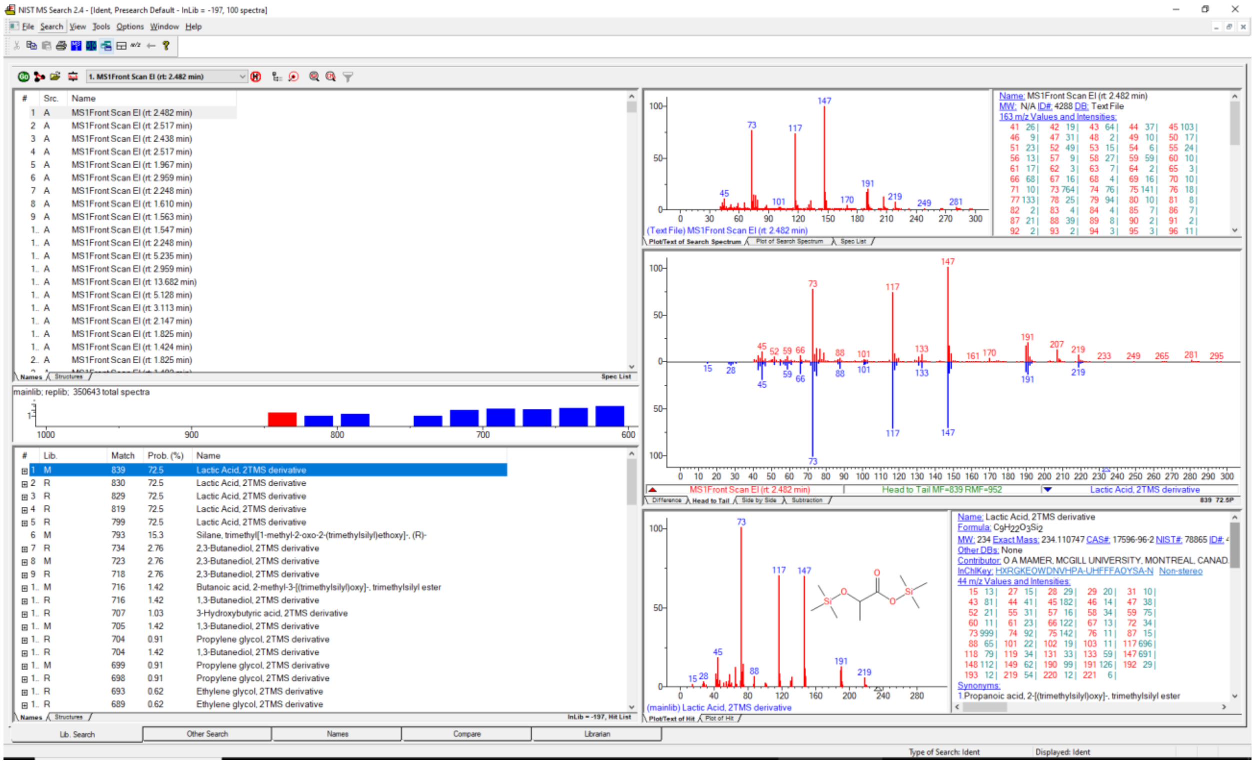

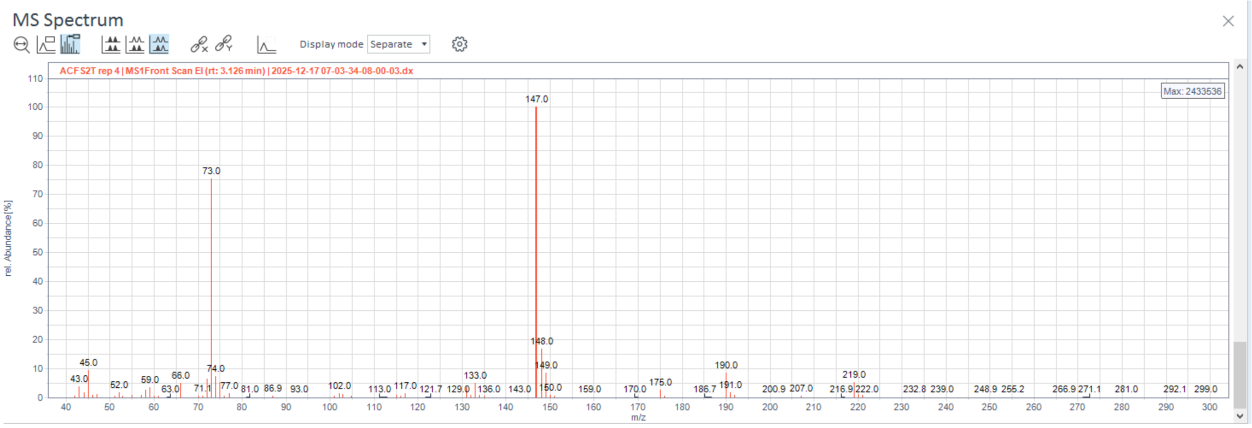
MS Spectrum
Display mode Separate
ACF S2T rep 4 | MS1Front Scan EI (rt: 3.126 min) | 2025-12-17 07-03-34-08-00-03.dx
Max: 2433536
rel. Abundance [%]
m/z
147.0
73.0
148.0
149.0
45.0
190.0
219.0

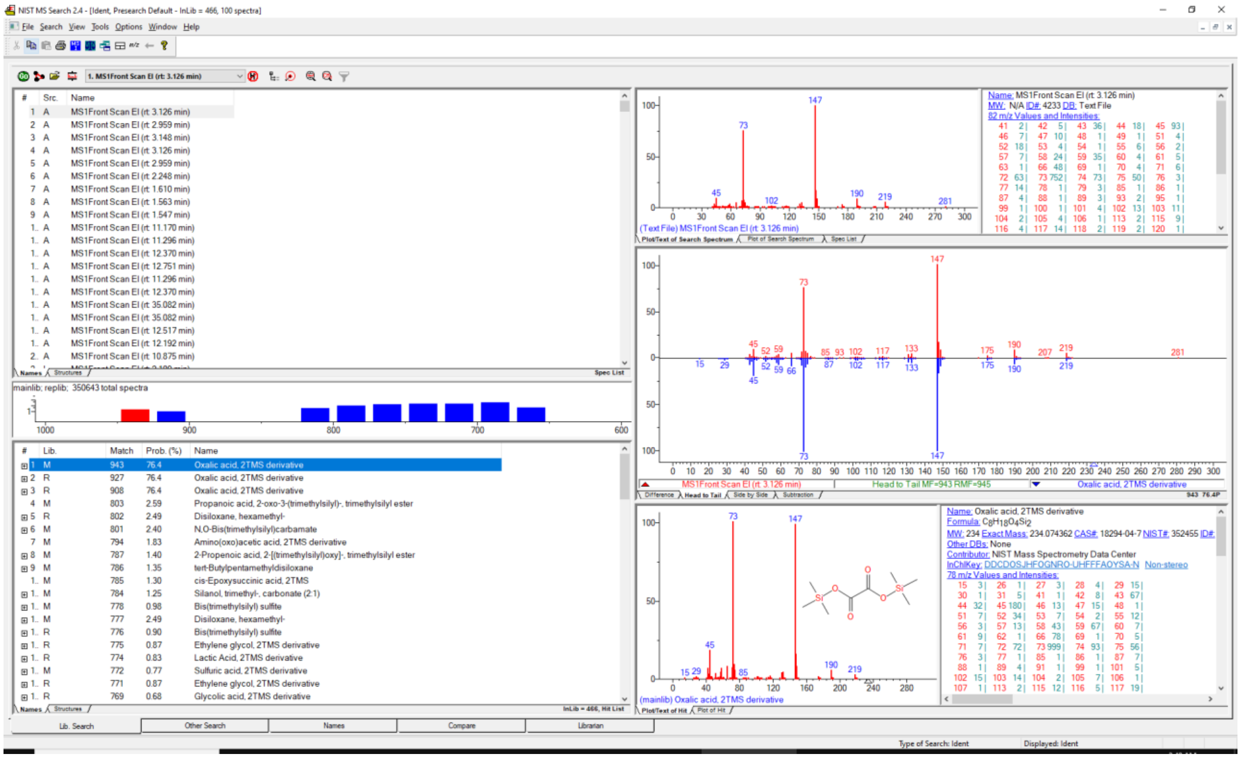
NIST MS Search 2.4
1. MS1Front Scan EI (rt: 3.126 min)
mainlib; replib; 350643 total spectra
Oxalic acid, 2TMS derivative
Name: Oxalic acid, 2TMS derivative
NIST Mass Spectrometry Data Center
(Text File) MS1Front Scan EI (rt: 3.126 min)
(mainlib) Oxalic acid, 2TMS derivative
Head to Tail MF=943 RMF=945
Lib. Search
Other Search
Names
Compare
Librarian

#### *Absidia psychrophilia* NRRL 3044

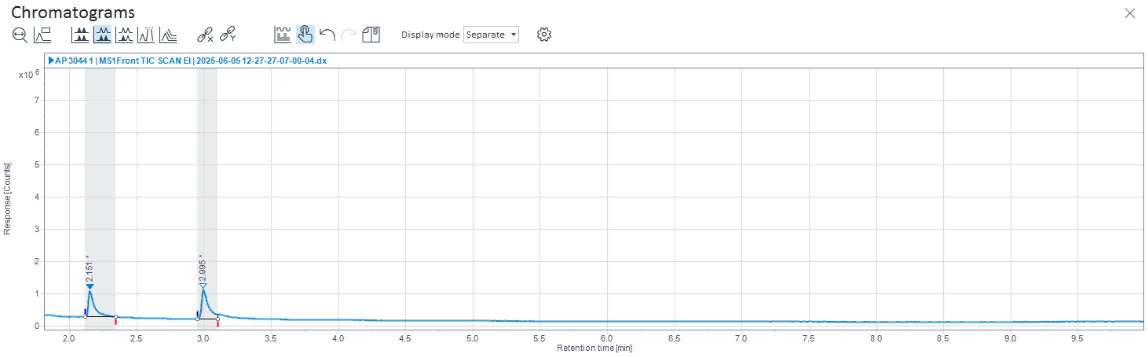

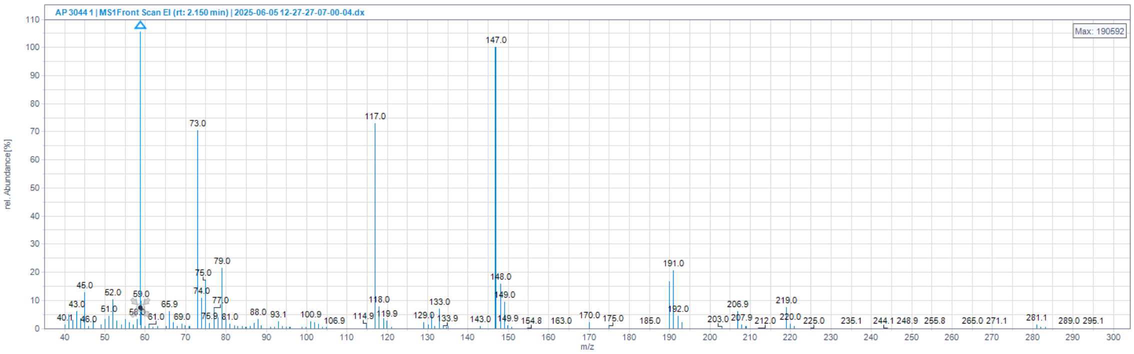

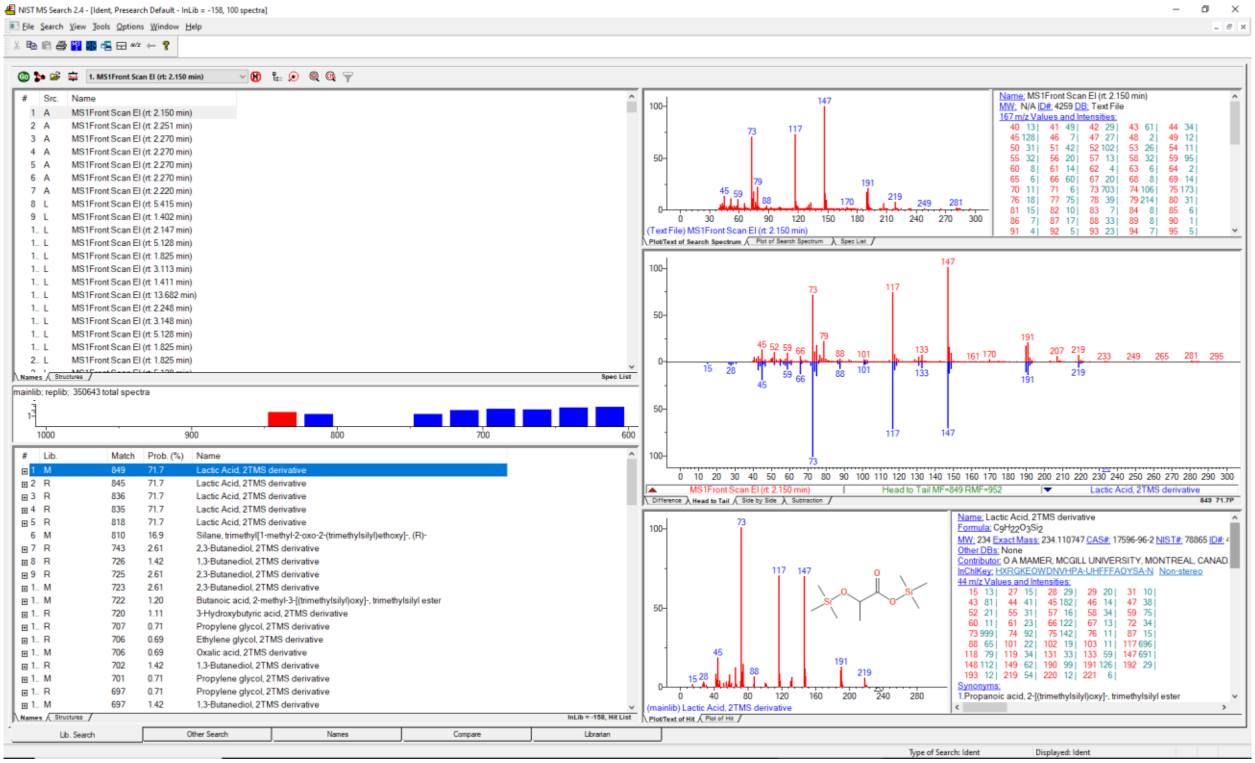

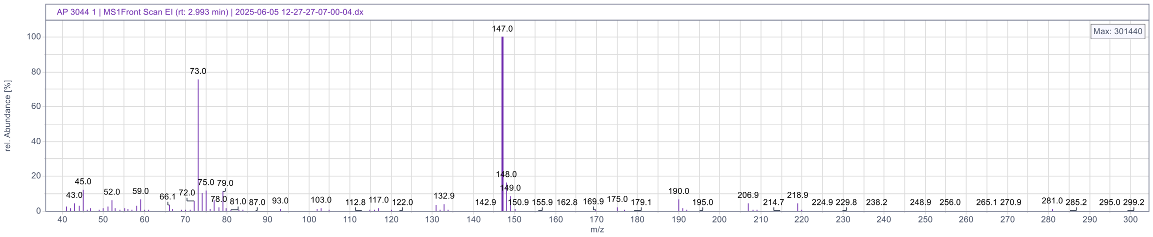

AP 3044 1 | MS1Front Scan EI (rt: 2.993 min) | 2025-06-05 12-27-27-07-00-04.dx
Max: 301440
rel. Abundance [%]
m/z
147.0
73.0

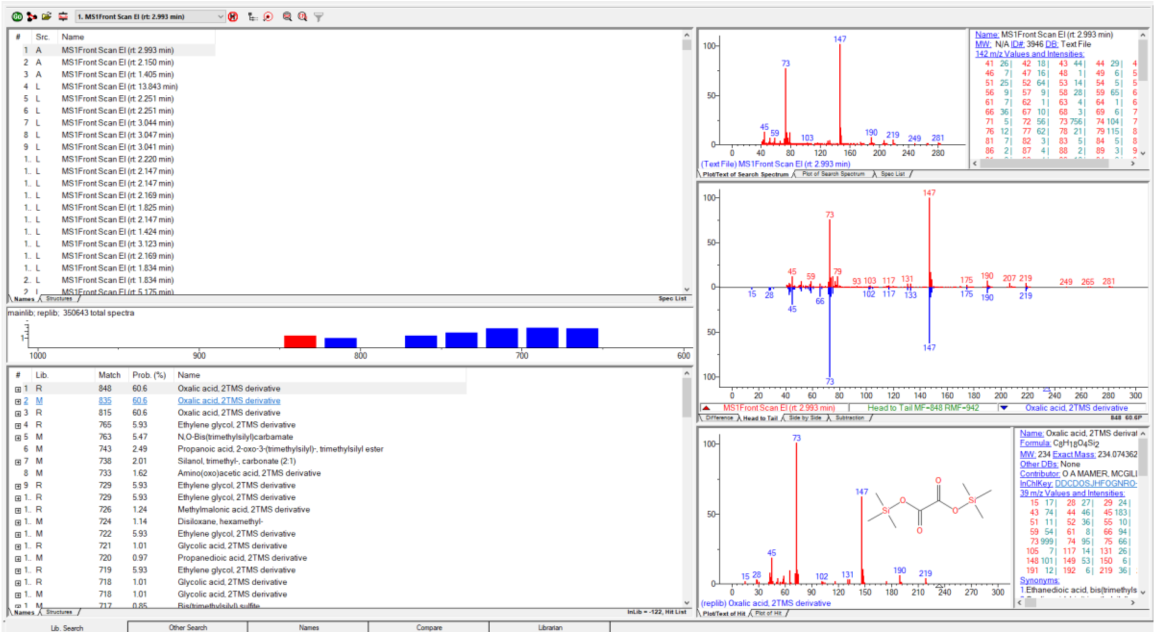

1. MS1Front Scan EI (rt: 2.993 min)
mainlib; replib; 350643 total spectra
Oxalic acid, 2TMS derivative
Ethylene glycol, 2TMS derivative
N,O-Bis(trimethylsilyl)carbamate
Propanoic acid, 2-oxo-3-(trimethylsilyl)-, trimethylsilyl ester
Silanol, trimethyl-, carbonate (2:1)
Amino(oxo)acetic acid, 2TMS derivative
Methylmalonic acid, 2TMS derivative
Disiloxane, hexamethyl-
Glycolic acid, 2TMS derivative
Propanedioic acid, 2TMS derivative
Head to Tail MF=848 RMF=942
Name: Oxalic acid, 2TMS derivative
Formula: C8H18O4Si2
MW: 234 Exact Mass: 234.074362
Lib. Search
Other Search
Names
Compare
Librarian

#### *Cunninghamella blakesleeana* CCM F705

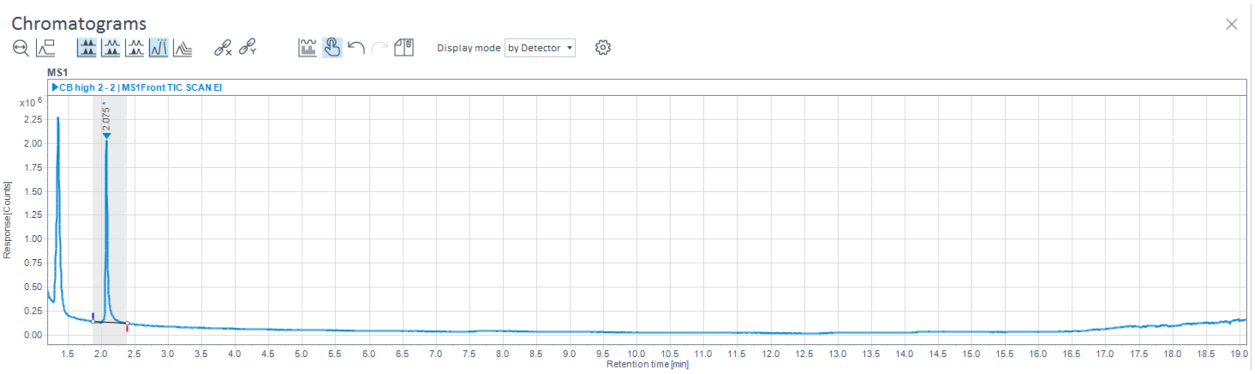

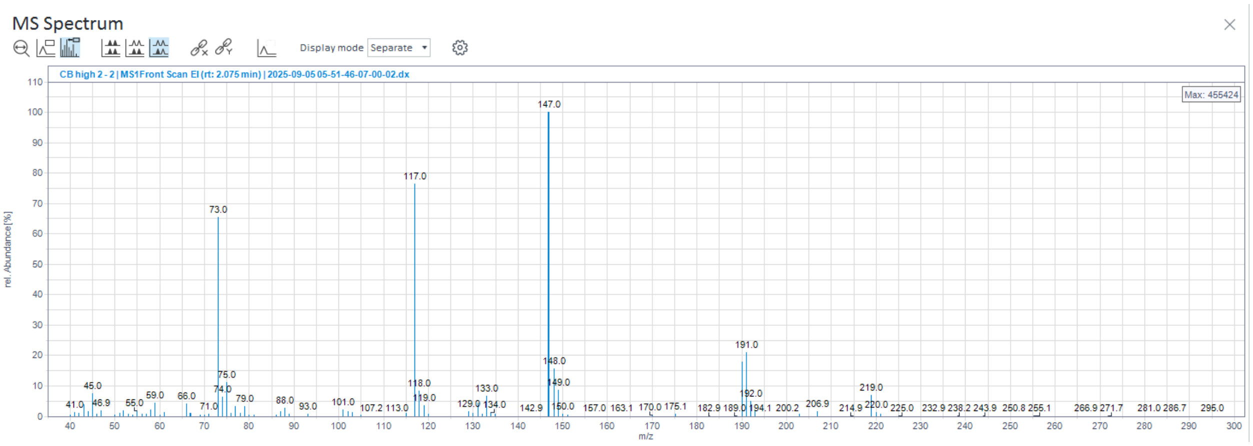

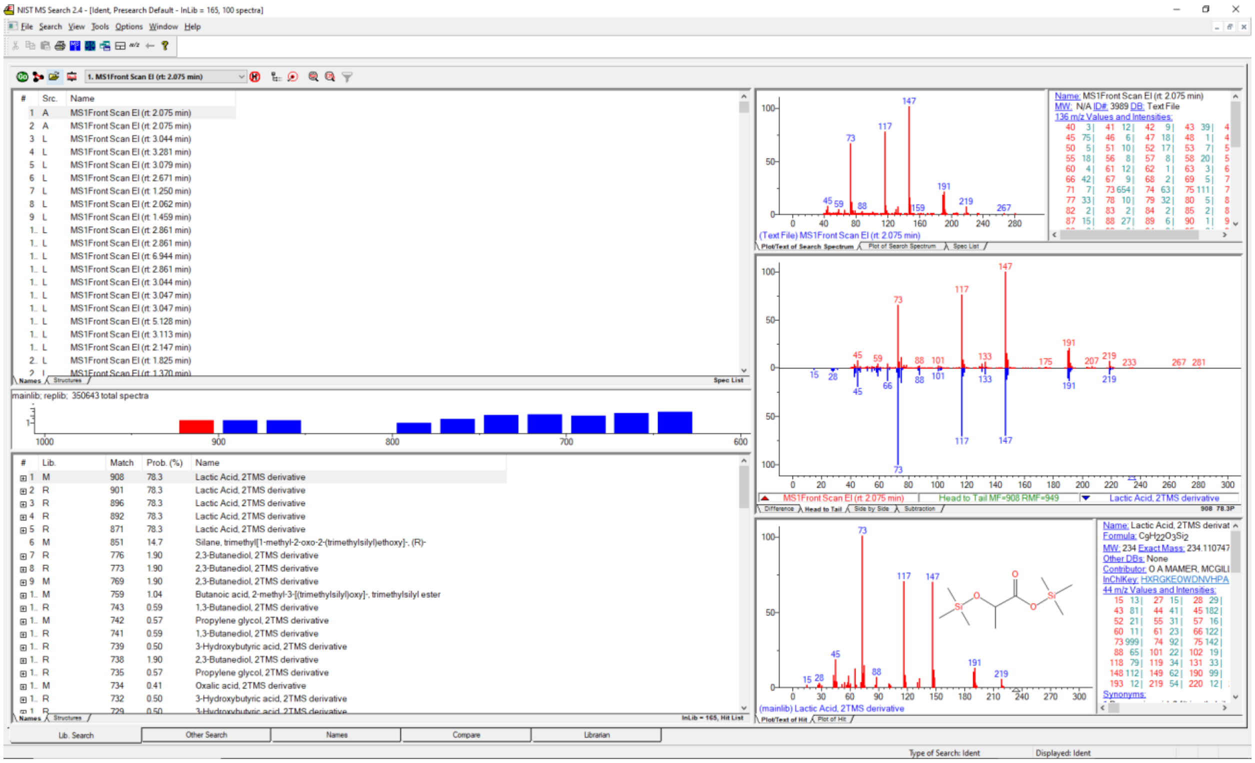

***Lichtheimia blakesleeana* NRRL 2696**

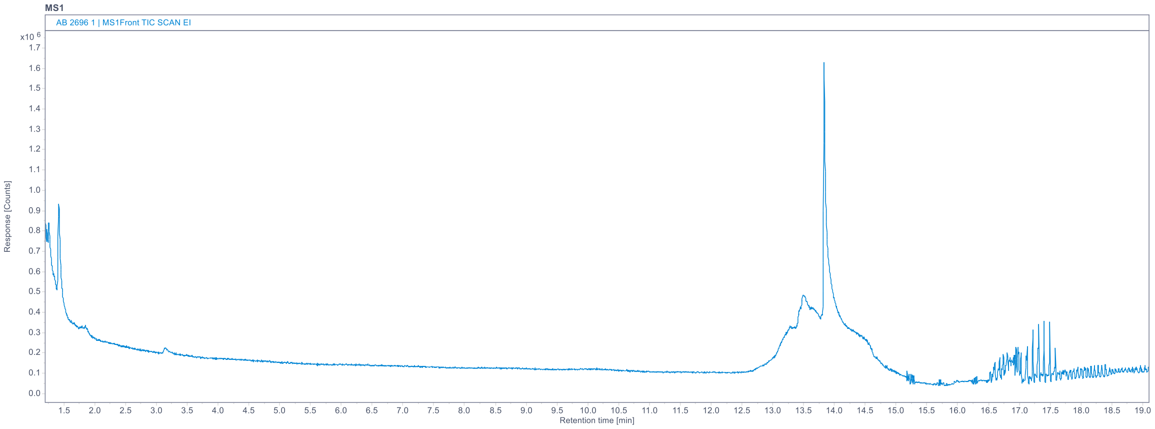

No short-chain acids detected in this sample

***Lichtheimia corymbifera* CCM F8077**

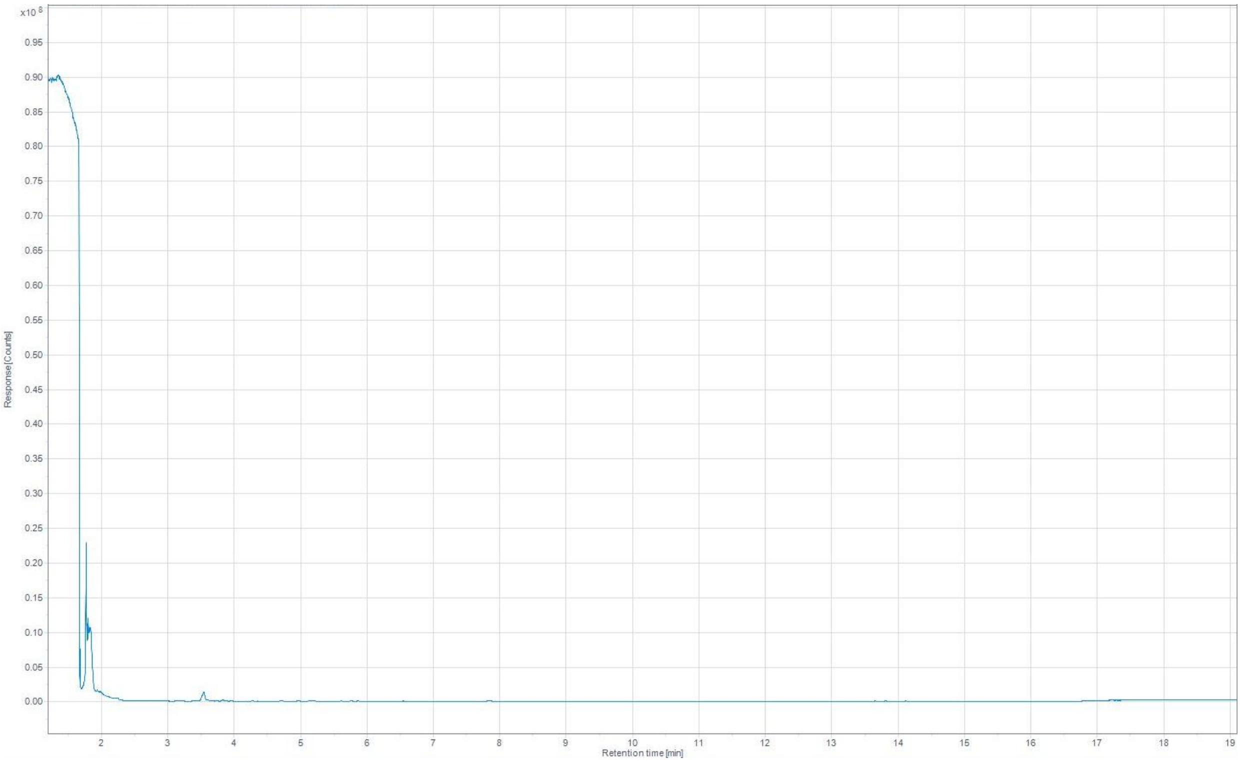

No short-chain acids detected in this sample

#### ***Mortierella alpina* ATCC 32222**

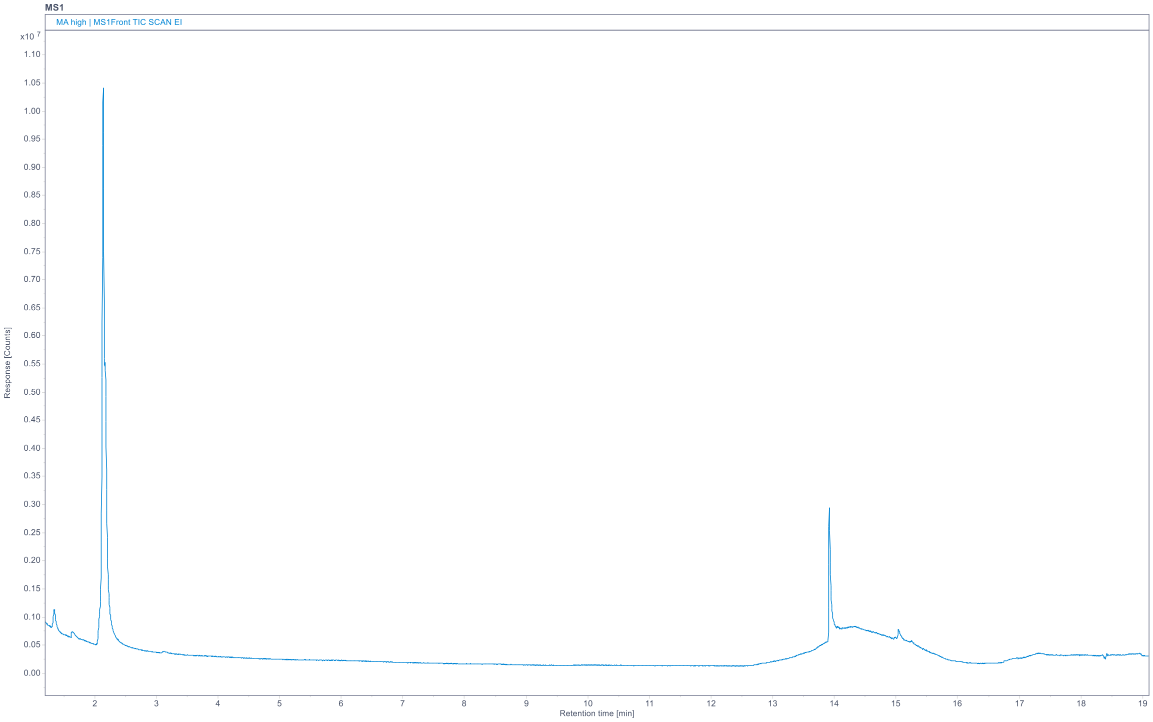

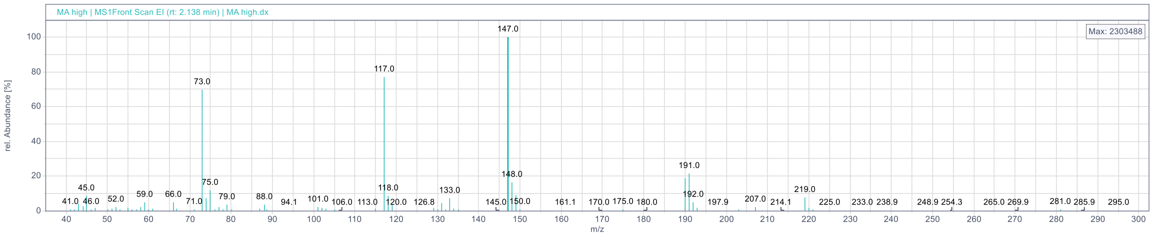

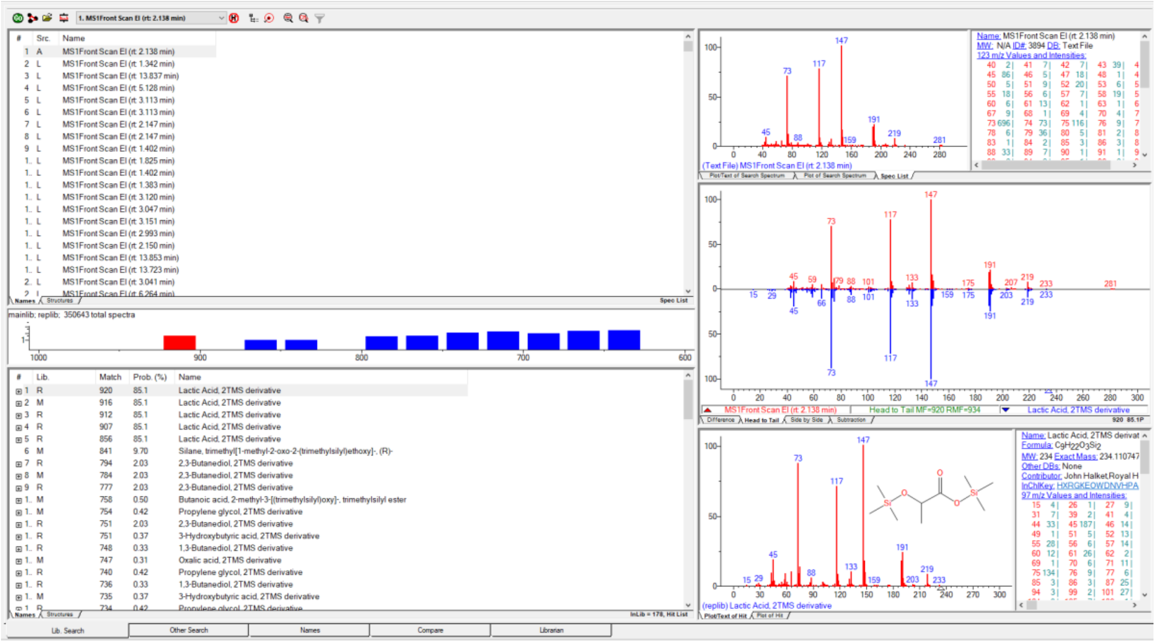

#### *Mortierella hyalina* VKM F1629

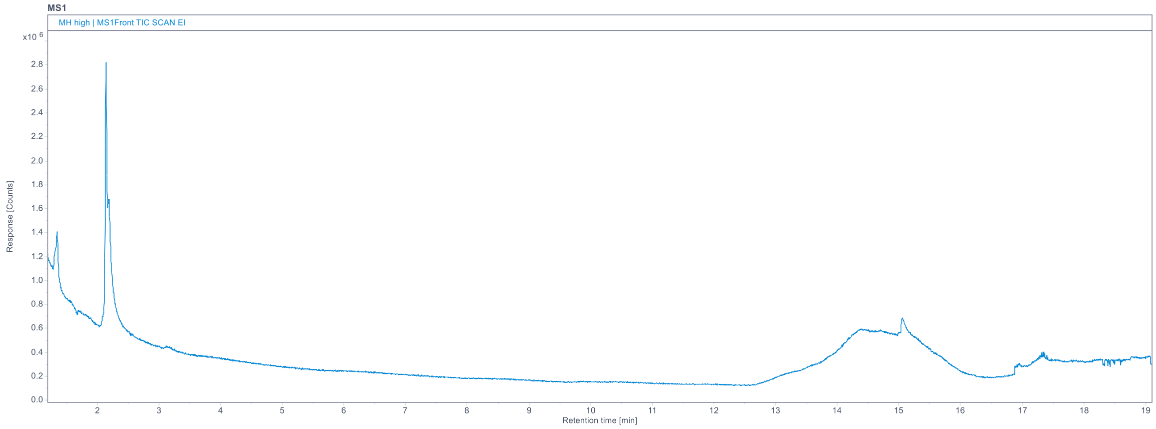

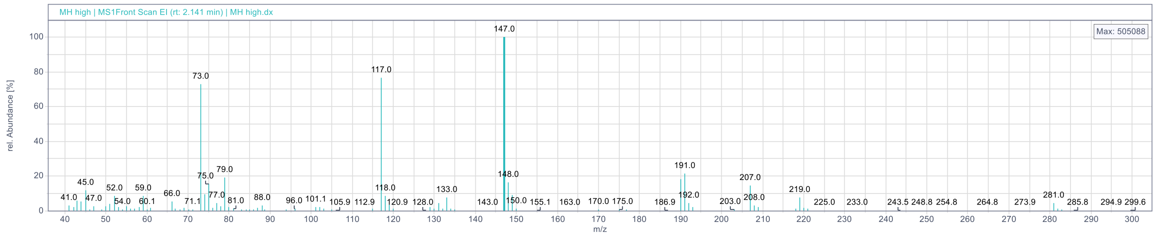

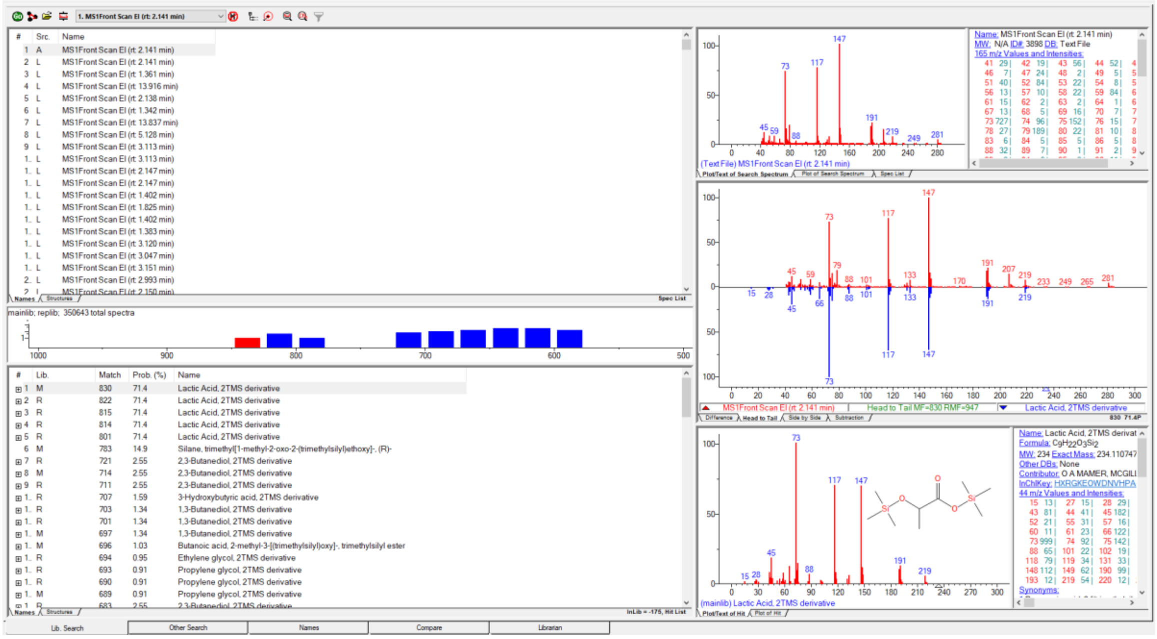

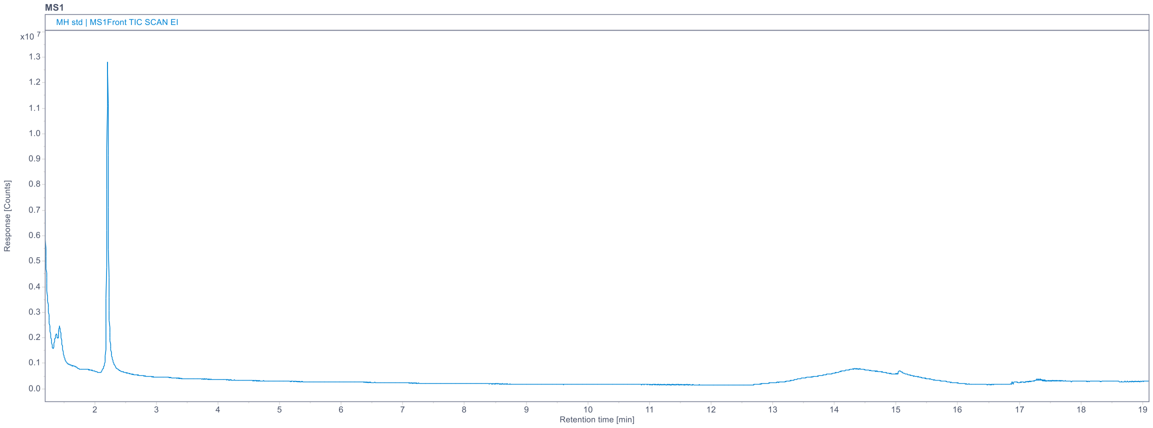
MS1
MH std | MS1Front TIC SCAN EI
Response [Counts]
Retention time [min]

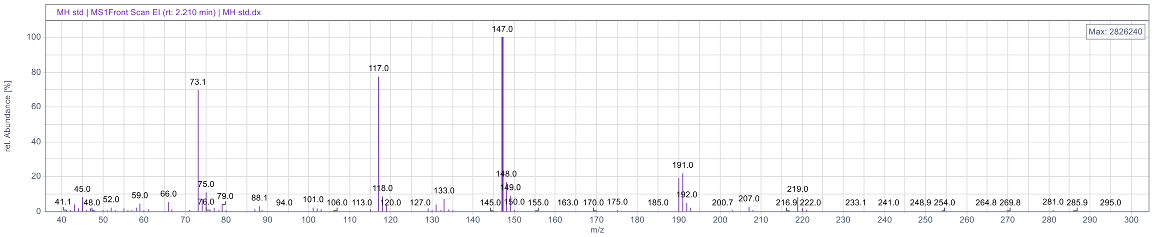
MH std | MS1Front Scan EI (rt: 2.210 min) | MH std.dx
Max: 2826240
rel. Abundance [%]
m/z
147.0
117.0
73.1
191.0

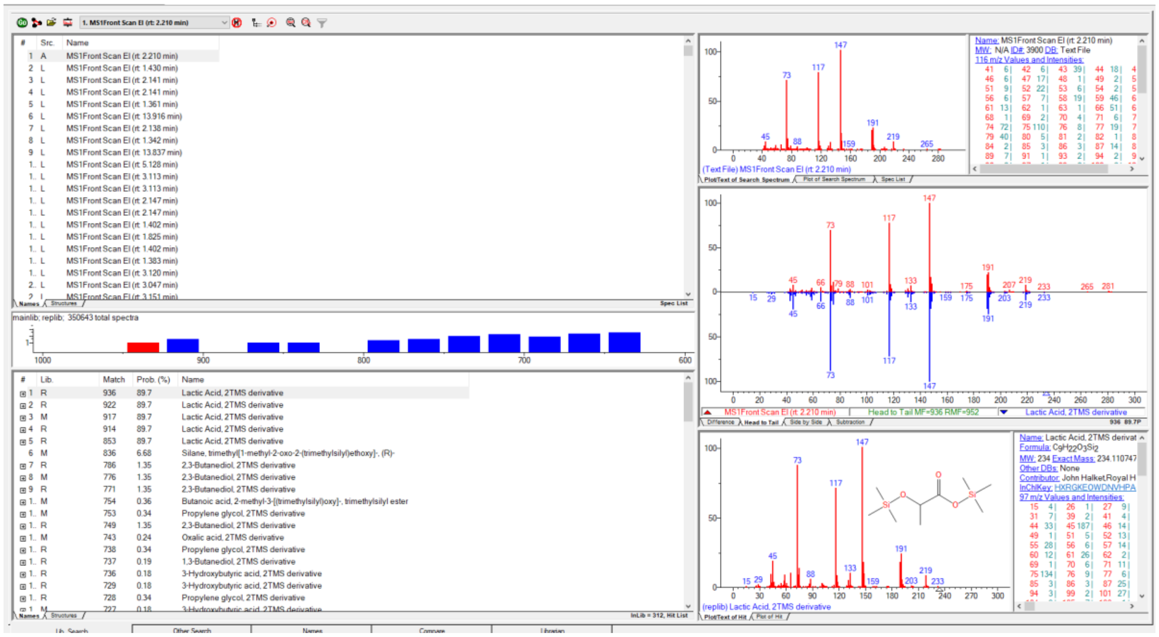
1. MS1Front Scan EI (rt: 2.210 min)
mainlib; replib; 350643 total spectra
Lactic Acid, 2TMS derivative
Head to Tail MF=936 RMF=952
(replib) Lactic Acid, 2TMS derivative

#### *Mucor circinelloides* CCM F220

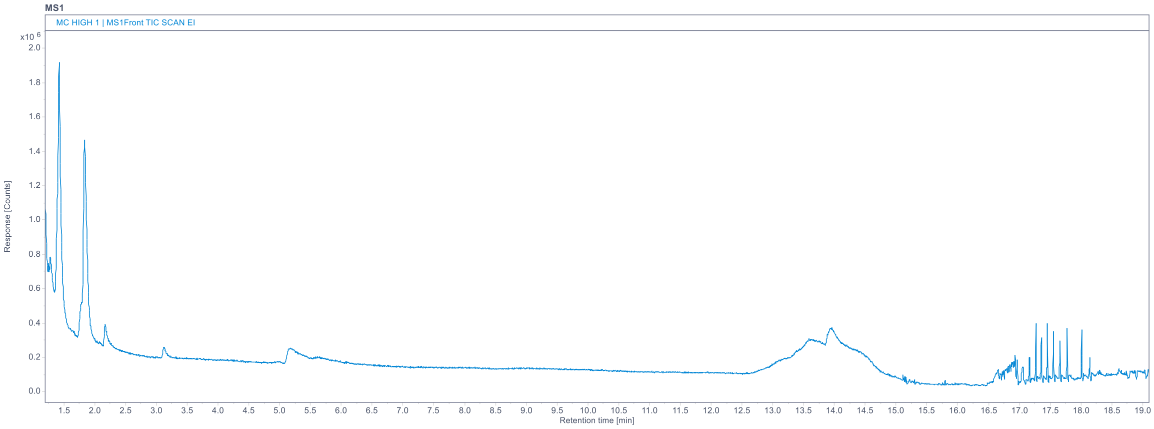

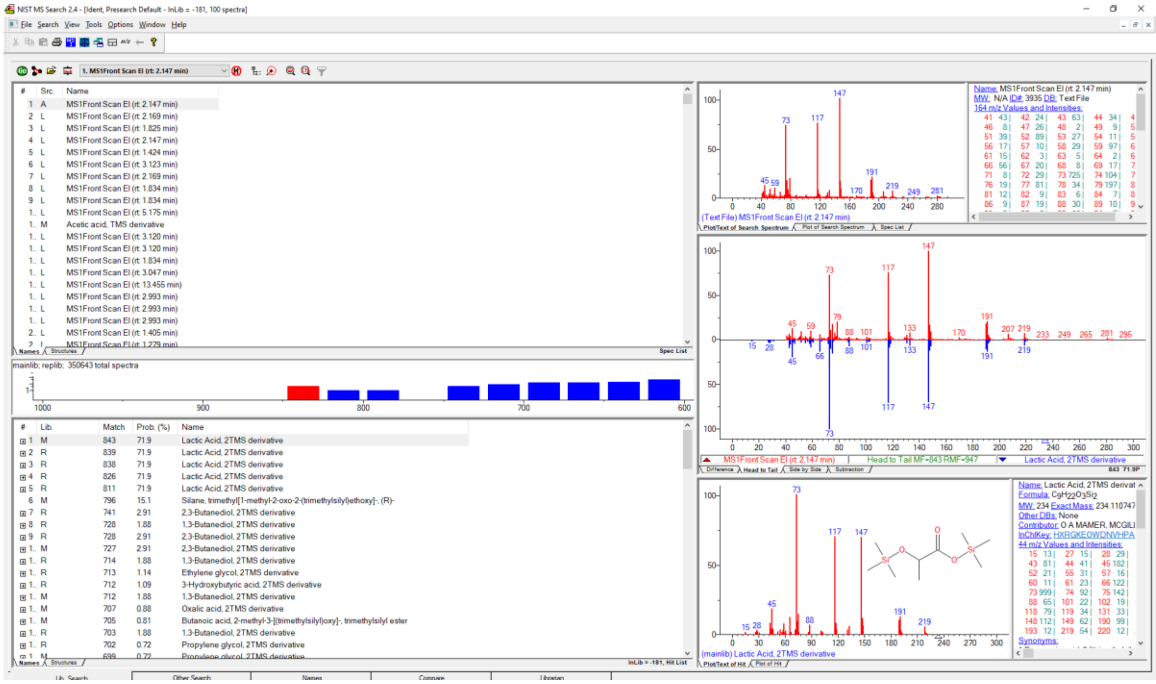

#### *Mucor circinelloides* VI04473

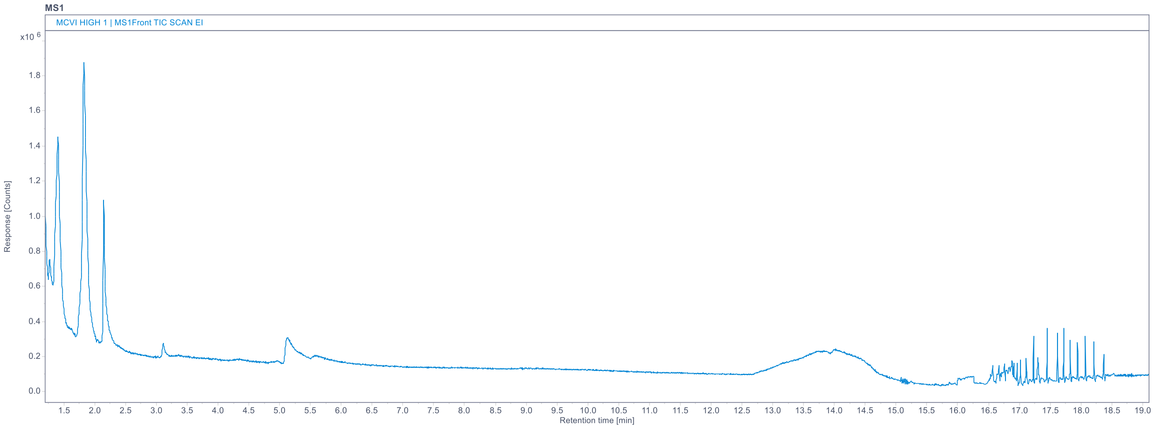

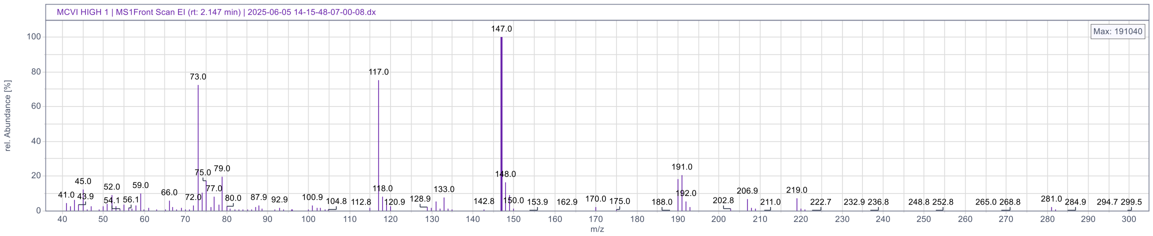

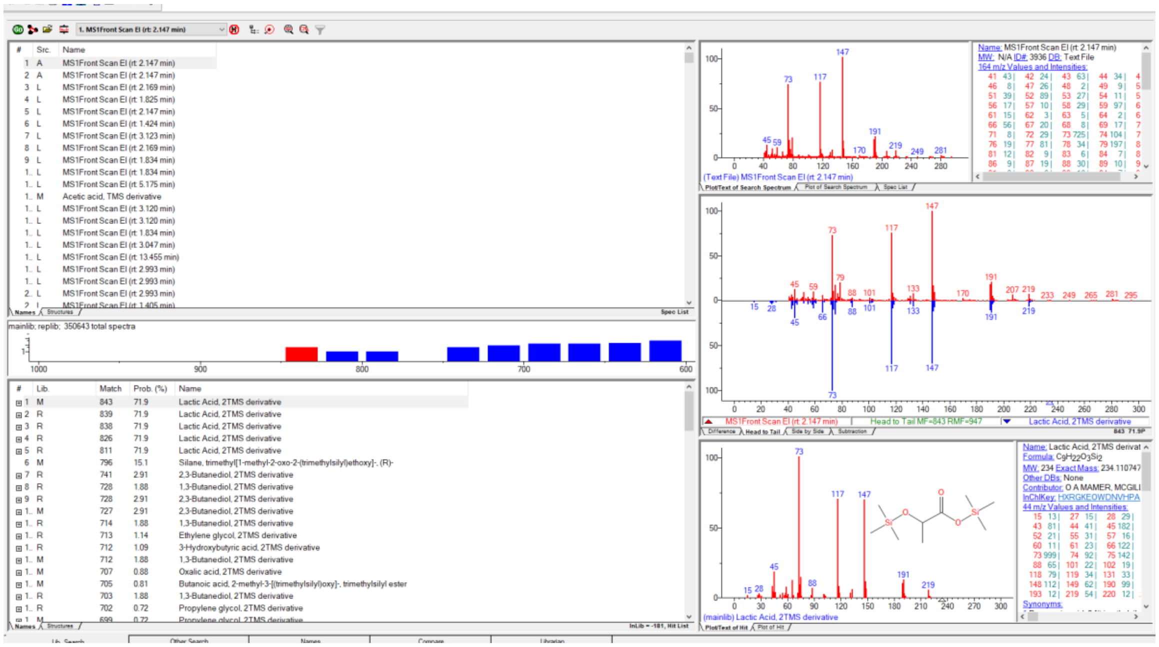

#### *Rhizopus stolonifer* VKM F400

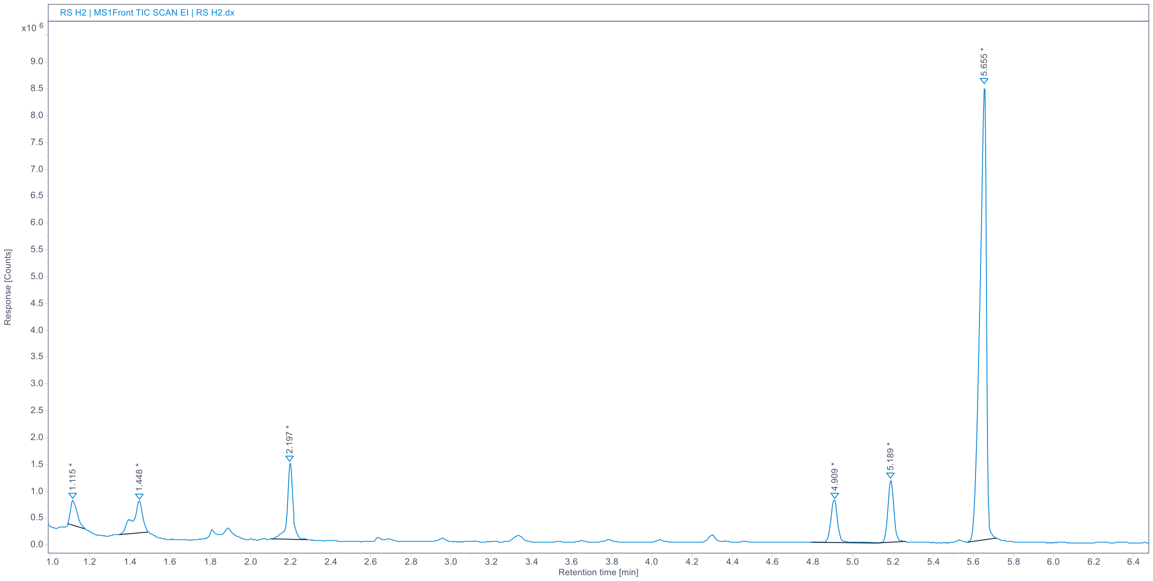

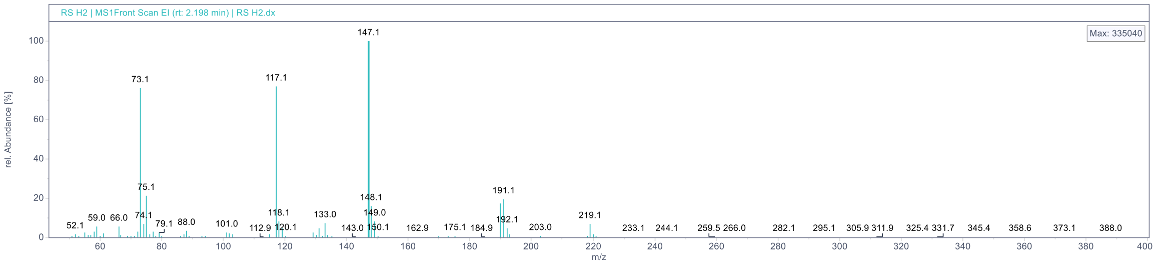

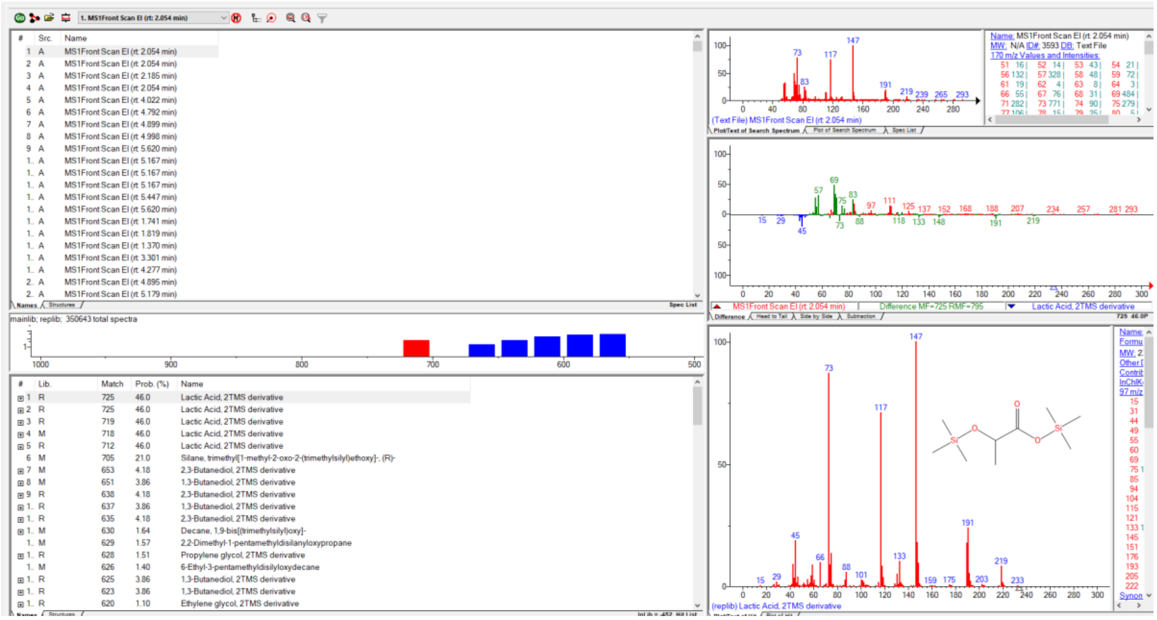

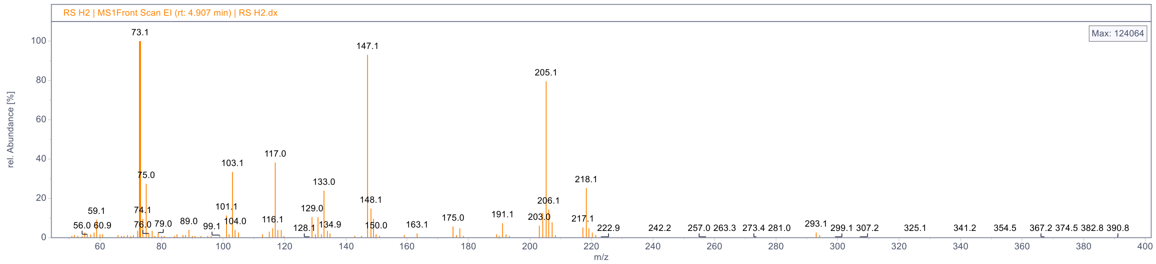
RS H2 | MS1Front Scan EI (rt: 4.907 min) | RS H2.dx
Max: 124064
rel. Abundance [%]
m/z

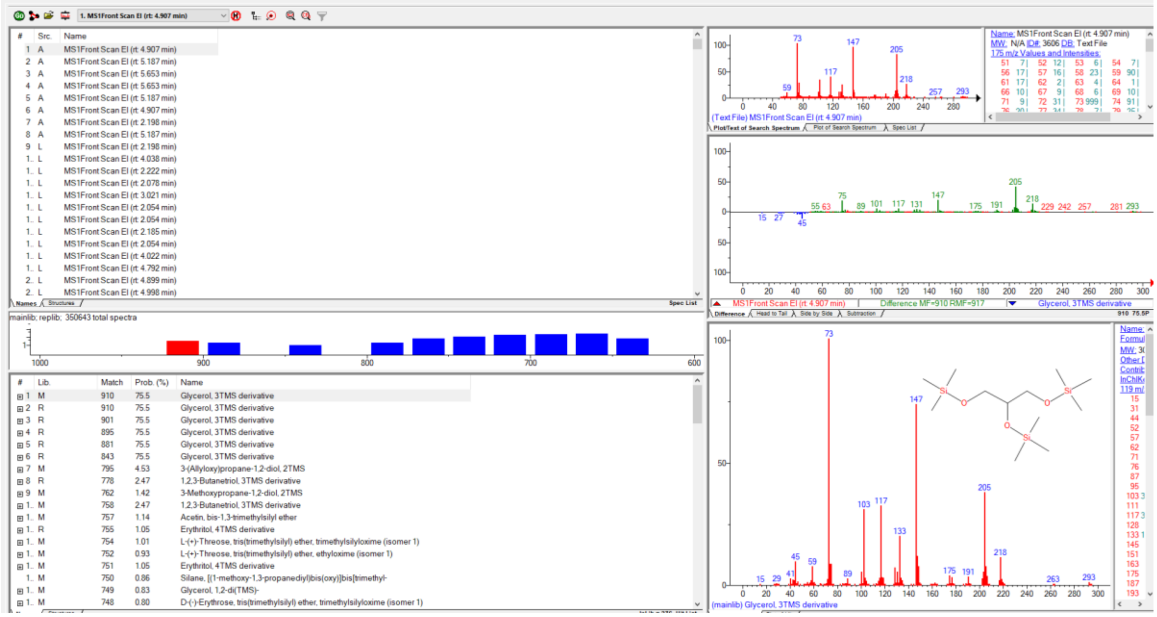
MS1Front Scan EI (rt: 4.907 min)
mainlib; replib; 350643 total spectra
Glycerol, 3TMS derivative
(mainlib) Glycerol, 3TMS derivative

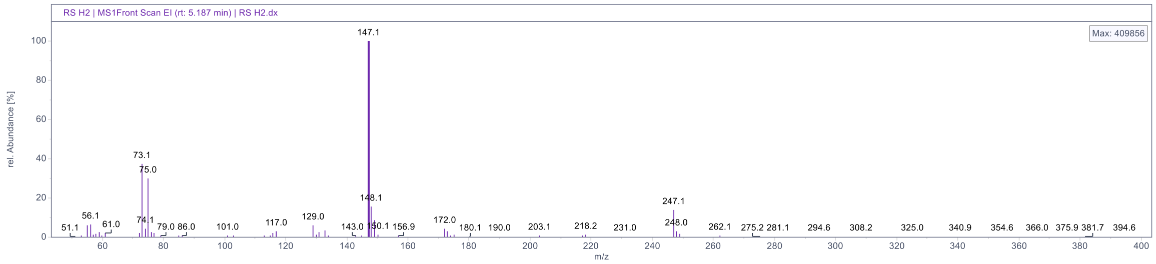
RS H2 | MS1Front Scan EI (rt: 5.187 min) | RS H2.dx
Max: 409856
rel. Abundance [%]
m/z

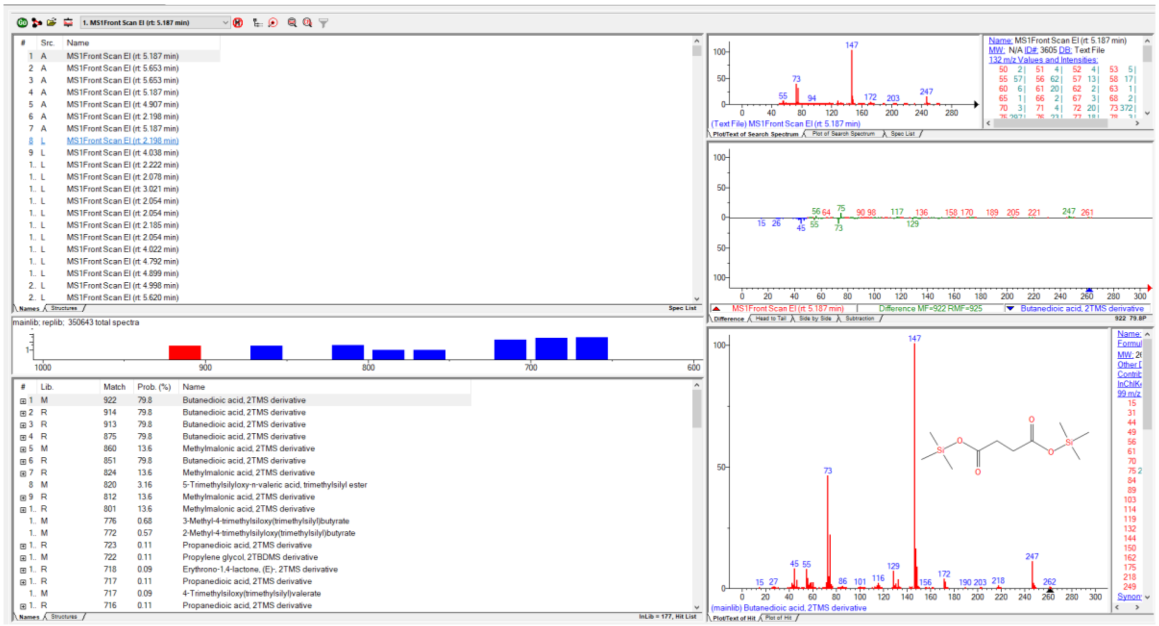
1. MS1Front Scan EI (rt: 5.187 min)
mainlib; replib; 350643 total spectra
Butanedioic acid, 2TMS derivative
(mainlib) Butanedioic acid, 2TMS derivative

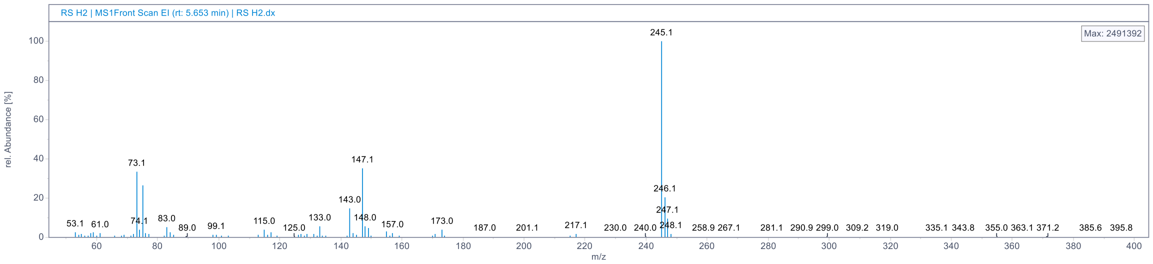
RS H2 | MS1Front Scan EI (rt: 5.653 min) | RS H2.dx
Max: 2491392
rel. Abundance [%]
m/z
245.1
147.1
73.1
143.0
246.1
247.1

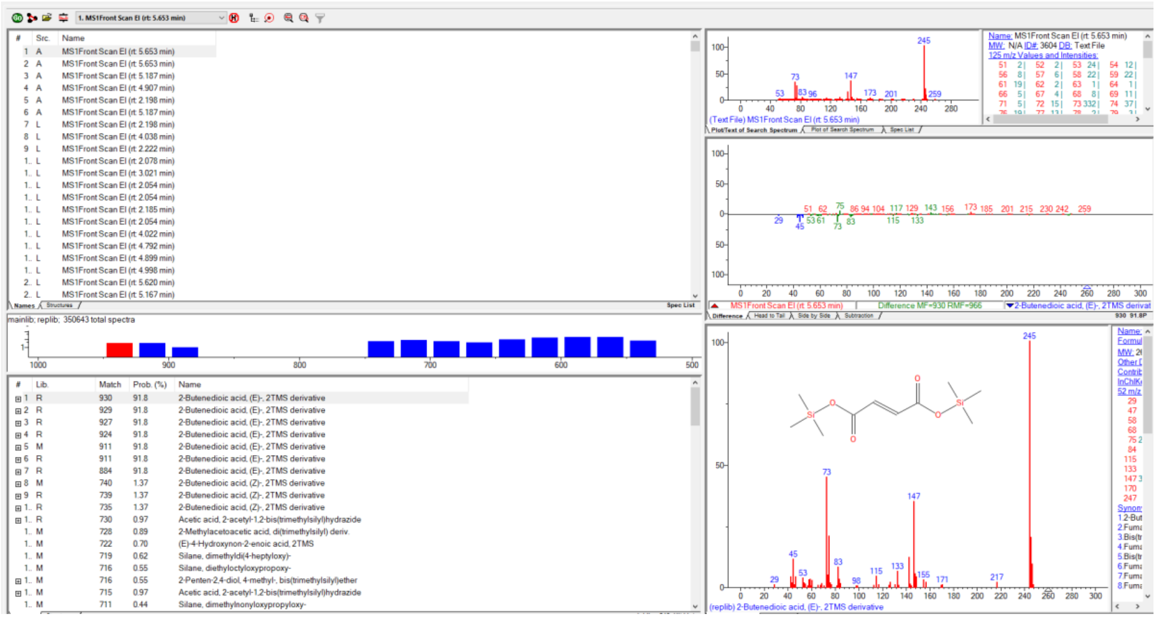
1. MS1Front Scan EI (rt: 5.653 min)
mainlib; replib; 350643 total spectra
2-Butenedioic acid, (E)-, 2TMS derivative
(replib) 2-Butenedioic acid, (E)-, 2TMS derivative

#### ***Umbelopsis vinacea* CCM F539**

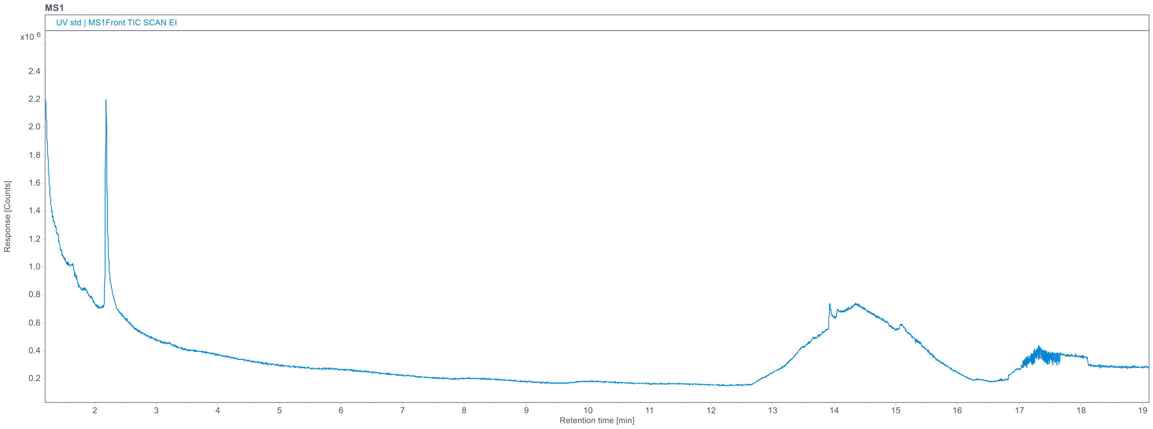

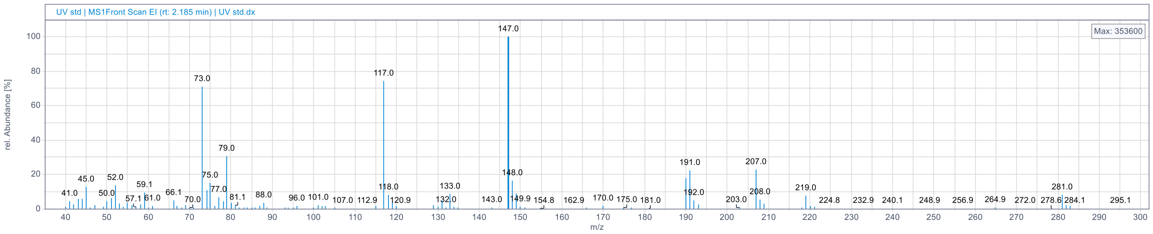

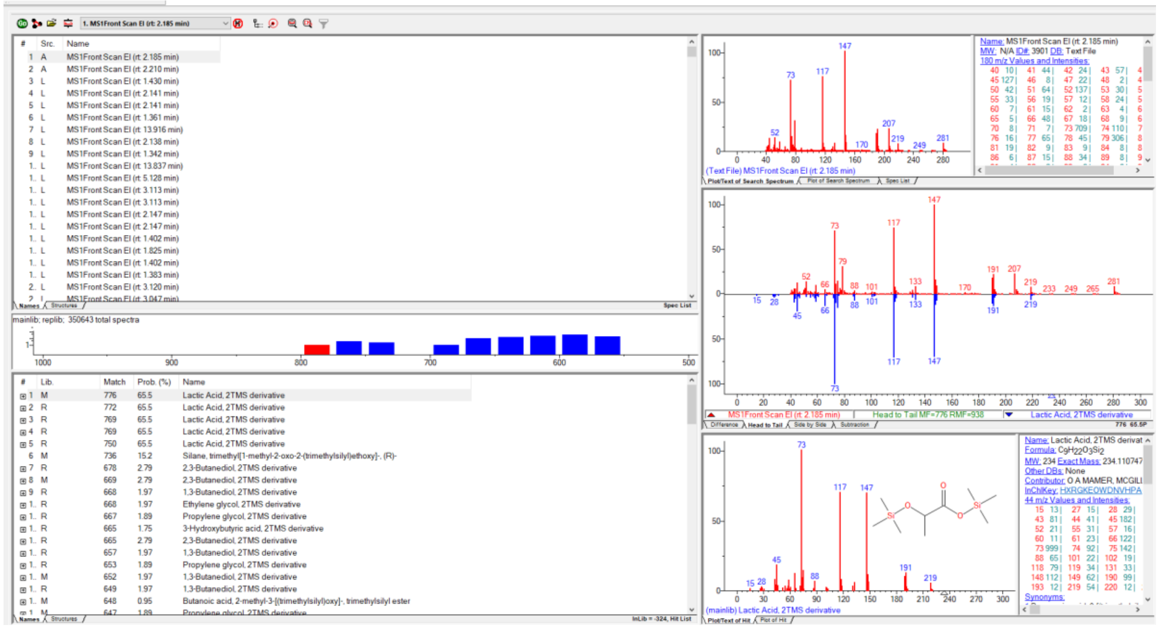
