## Supplementary file 4 for "Carbon allocation strategies across Mucoromycota and the discovery of oxalic acid secretion in *Absidia*"

### Carbon allocation strategies across Mucoromycota and the discovery of oxalic acid secretion in *Absidia*

Cristian Bolaño Losada[1], Boris Zimmermann[1], Dana Byrtusova[1], Sven Andreas Högfeldt[1], Svein Jarle Horn[2], Achim Kohler[1], Volha Shapaval[1]

[1] Faculty of Science and Technology, Norwegian University of Life Sciences (NMBU), 1432 Ås, Norway

[2] Faculty of Chemistry, Biotechnology and Food Science, Norwegian University of Life Sciences (NMBU), 1432 Ås, Norway.

**Table S1.** Dimensional characteristics of the deep well plates used for carrying out the fermentation experiments. These plates were used in combination with low evaporation sandwich covers and a clamp system from Enzyscreen (The Netherlands).

| | **Polypropylene square 96-deep-well microplates** | **Extra high polypropylene square deep-well microplates** |
|---|---|---|
| **Cat. No.** | Enzyscreen CR1496 | Enzyscreen CR1424hd |
| **No. of wells** | 96 | 24 |
| **Shape** | Square | Square |
| **Dimensions (W x D X H) mm** | 8 x 8 x 40 | 17 x 17 x 89 |
| **Well total volume (T) (mL)** | 2.2 | 25 |
| **Working volume (W) (mL)** | 0.8 | 7 |
| **W/T volume ratio** | 1/3 | 2/7 |

**Table S2.** Calcium, sodium, magnesium, potassium and phosphorus relative content in biomass of *A. glauca* CCM F451, *R. stolonifer* VKM F400, and *L. corymbifera* CCM F8077 grown in high C/N medium and 24-well plates. Samples corresponded to three biologically independent cultivations (biological replicates), each analyzed twice by the elemental composition method (technical replicates). n.d.: not detected.

| Strain | Biological replicate | Technical replicate | Ca | Na | Mg | K | P |
|---|---|---|---|---|---|---|---|
| | | | % | % | % | % | % |
| ***A. glauca* CCM F451** | 1 | 1 | 0.44 | n.d. | 0.45 | 0.71 | 2.81 |
| | | 2 | 0.23 | n.d. | 0.43 | 1.04 | 2.64 |
| | 2 | 1 | 0.31 | n.d. | 0.46 | 0.79 | 2.70 |
| | | 2 | 0.43 | n.d. | 0.50 | 0.52 | 2.95 |
| | 3 | 1 | 0.32 | n.d. | 0.42 | 0.91 | 2.62 |
| | | 2 | 0.37 | n.d. | 0.46 | 0.89 | 2.81 |
| ***R. stolonifer* VKM F400** | 1 | 1 | 0.31 | 0.29 | 0.76 | 2.68 | 3.85 |
| | | 2 | 0.29 | 0.28 | 0.83 | 2.78 | 4.04 |
| | 2 | 1 | 0.41 | n.d. | 1.11 | 1.94 | 4.67 |
| | | 2 | 0.41 | n.d. | 1.11 | 2.37 | 4.68 |
| | 3 | 1 | 0.29 | n.d. | 0.76 | 1.91 | 3.58 |
| | | 2 | 0.33 | 0.22 | 0.86 | 2.56 | 4.20 |
| ***L. corymbifera* CCM F8077** | 1 | 1 | 0.24 | 0.10 | 0.37 | 2.19 | 1.55 |
| | | 2 | 0.23 | 0.18 | 0.36 | 2.24 | 1.57 |
| | 2 | 1 | 0.21 | 0.20 | 0.37 | 2.27 | 1.59 |
| | | 2 | 0.22 | 0.18 | 0.37 | 2.21 | 1.56 |
| | 3 | 1 | 0.22 | 0.16 | 0.39 | 2.23 | 1.54 |
| | | 2 | 0.23 | 0.14 | 0.38 | 2.15 | 1.50 |

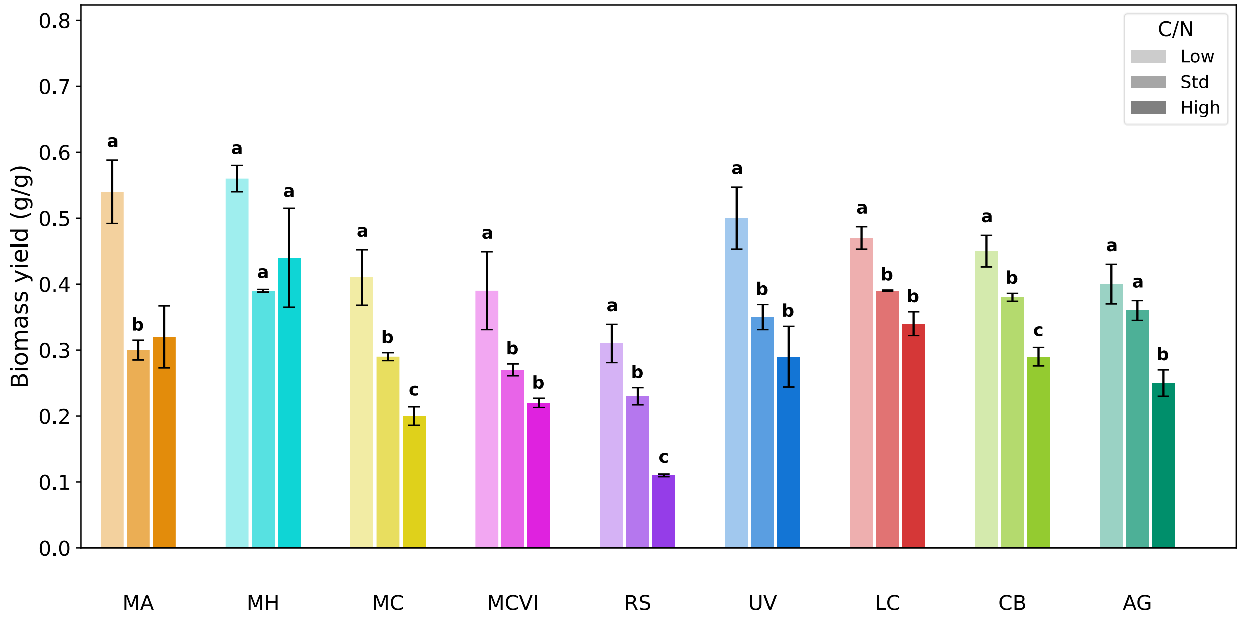


**Figure S1.** Biomass yield (biomass g/g glucose) for *Mortierella alpina* (MA), *Mortierella hyalina* (MH), *Mucor circinelloides* (MC), *Mucor circinelloides* (MCVI), *Rhizopus stolonifer* (RS), *Umbelopsis vinacea* (UV), *Lichtheimia corymbifera* (LC), *Cunninghamella blakesleeana* (CB), and *Absidia glauca* (AG) grown in low, standard and high C/N media. Error bars indicate the standard deviation from three biologically independent replicates. Letters indicate significant differences ($p < 0.05$) between C/N conditions, while their absence indicates no significant differences.

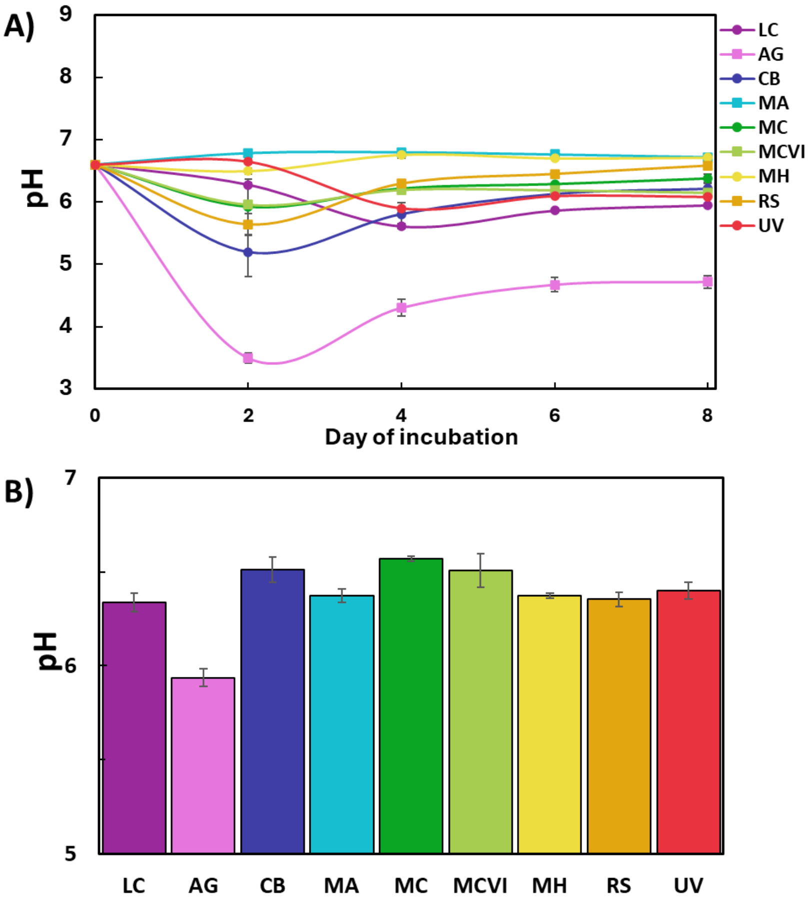


**Figure S2.** pH of supernatants after fermentation of *Lichtheimia corymbifera* CCM F8077 (LC), *Absidia glauca* CCM F451 (AG), *Cunninghamella blakesleeana* CCM F705 (CB), *Mortierella alpina* ATCC 32222 (MA), *Mucor circinelloides* CCM F220 (MC), *M. circinelloides* VI04473 (MCVI), *Mortierella hyalina* VKM F1629 (MH), *Rhizopus stolonifer* VKM F400 (RS) and *Umbelopsis vinacea* CCM F539 (UV) in A) cellobiose-based medium and B) cellulose-based. Details of the cultivation and medium conditions available in Bolaño Losada et al. (2025). Data and graphs reproduced from under rights of Creative Commons Attribution-NonCommercial-NoDerivatives 4.0 International License.

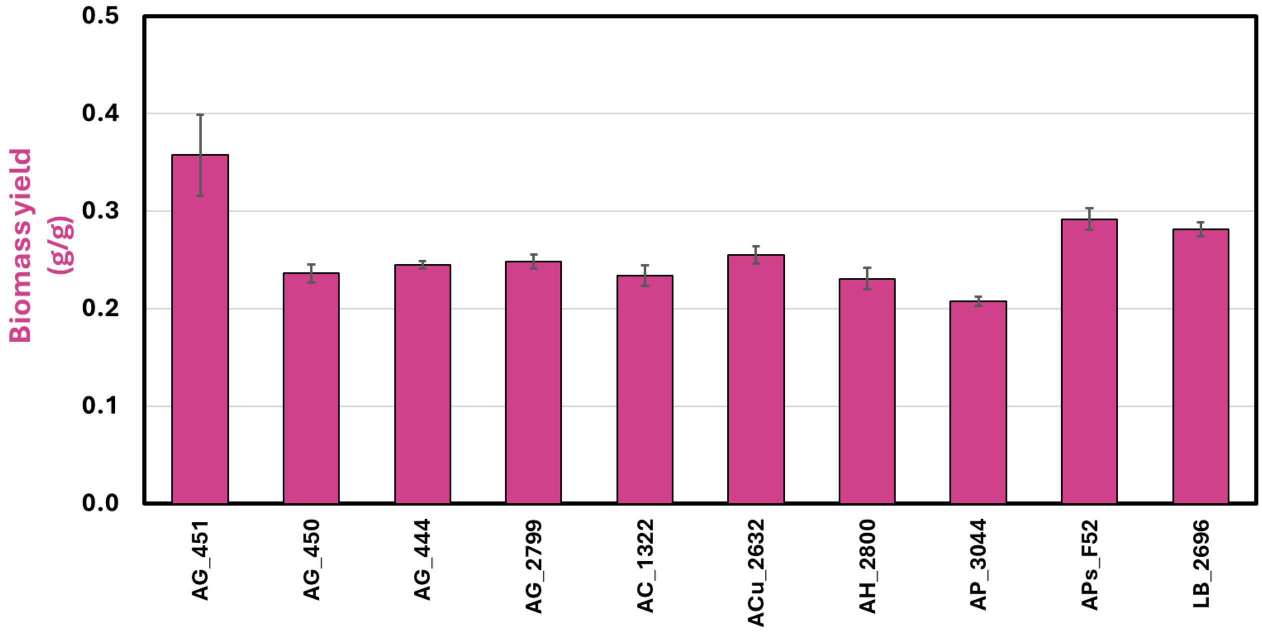


**Figure S3.** Biomass yields (biomass g/g glucose) from *Absidia glauca* (AG), *Absidia coerulea* (AC), *Absidia cuneospora* (ACu), *Absidia hyalospora* (AH), *Absidia psychrophilia* (AP), *Absidia pseudocylindrospora* (APs) and *Lichtheimia blakesleeana* (LB) strains grown in high C/N medium. Numbers indicate the culture collection number which can be fully found in table I.

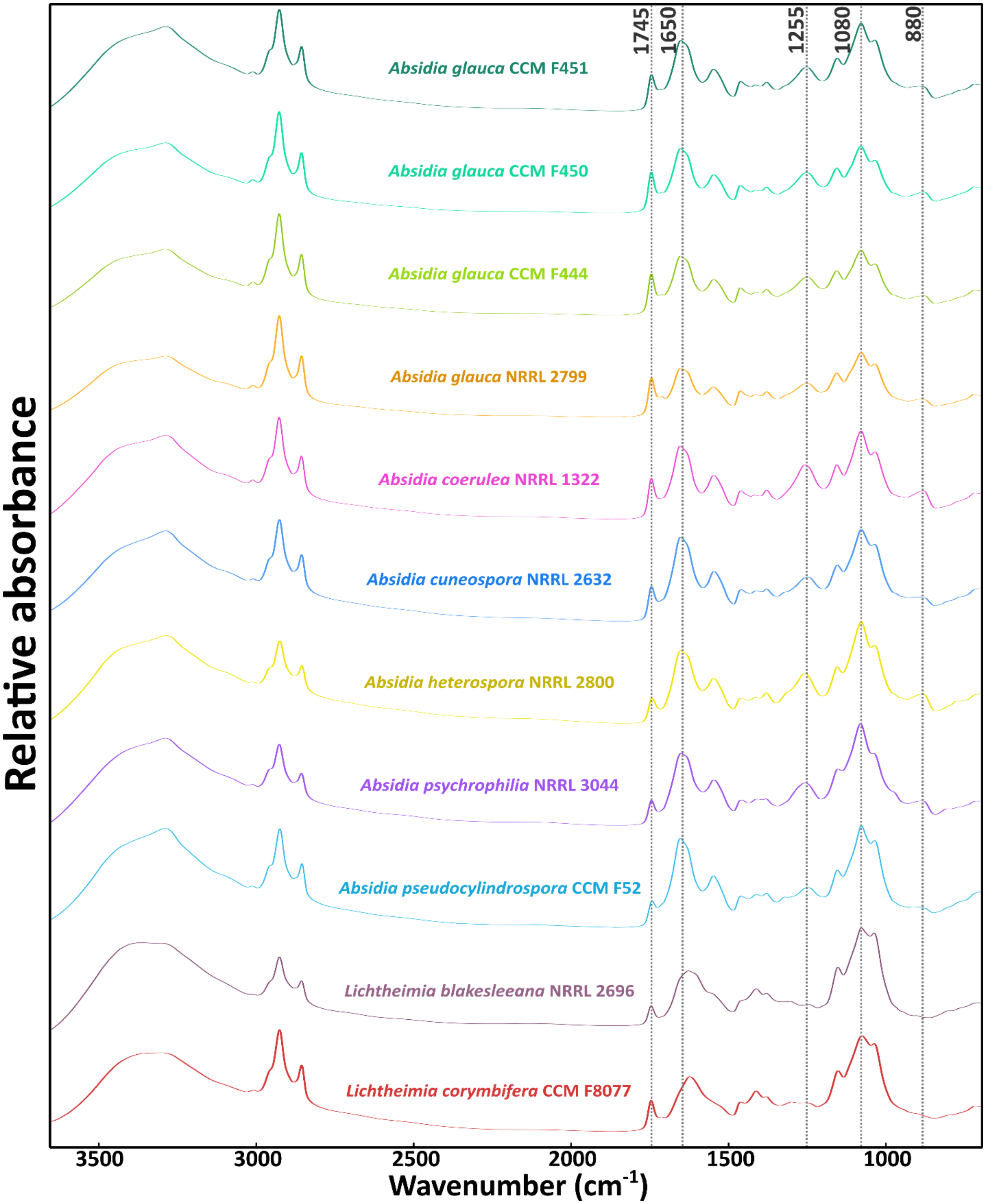


**Figure S4.** Stack graph of spectra from strains of *Lichtheimia* and *Absidia* from strain Set III grown in high C/N medium. The displayed spectra are the average of three biologically independent replicates each one measured three times. Relevant peaks related to different metabolites are added for assist the interpretation. Lipids (fatty acids) at 1745 $cm^{-1}$ (C=O stretching), proteins and chitin/chitosan at 1650 $cm^{-1}$ (amide I), (poly)phosphates at 1255 and 880 $cm^{-1}$ (P=O and P-O-P stretching, respectively), and saccharides at 1080 $cm^{-1}$ (C-O stretching).

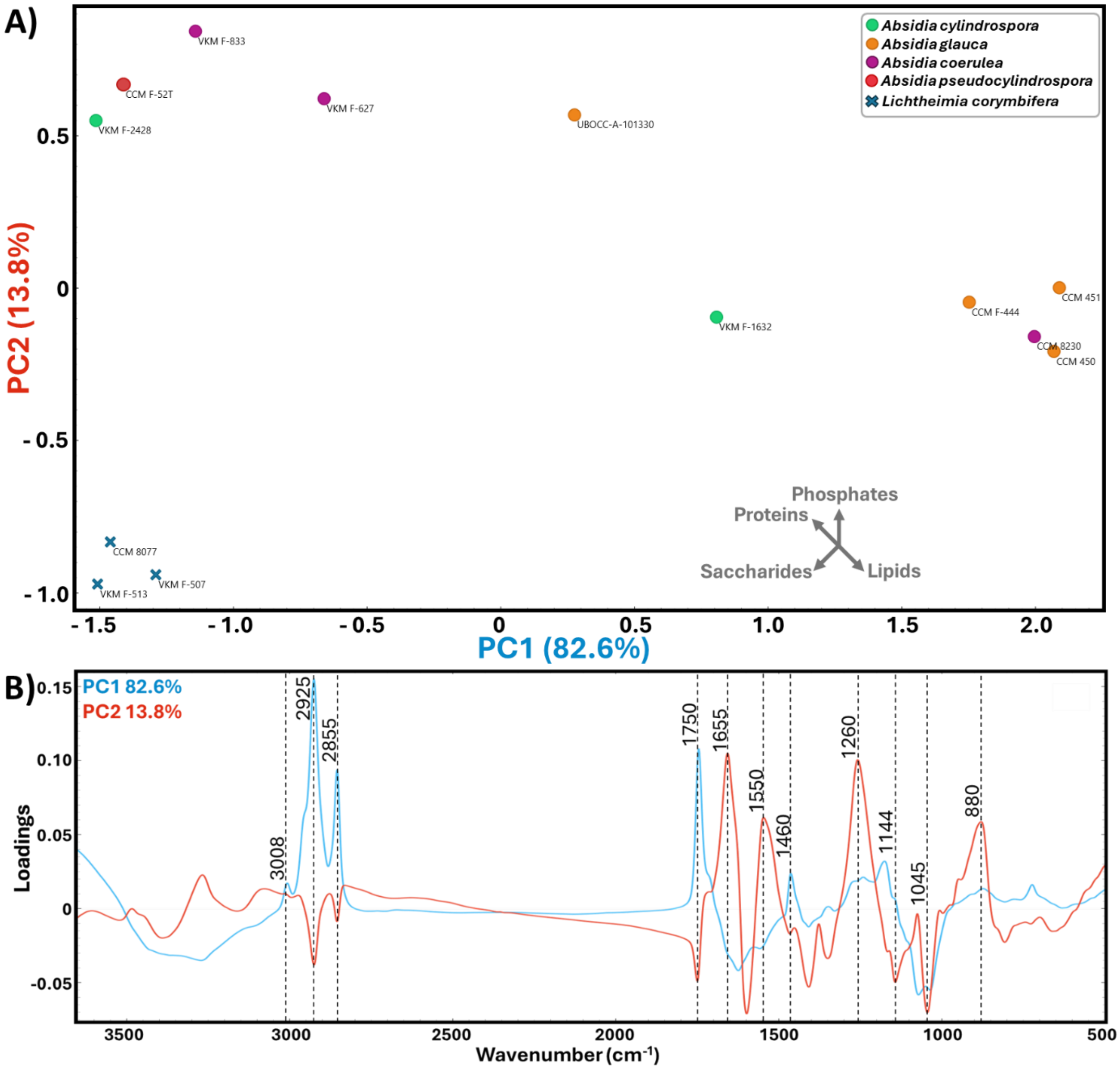


**Figure S5.** A) PCA scatter plot, and B) score plot of PC1 (blue) and PC2 (red) from the spectra of *A. cylindrospora*, *A. pseudocylindrospora*, *A. glauca*, *A. coerulea* and *L. corymbifera* strains grow at high C/N medium obtained in the study of Kosa et al. (2018). In A) an approximate axis is provided reflecting increases in relative content of metabolites according to the interpretation of the principal component loadings. Each dot corresponds to the average spectra of three biologically independent replicates. Labels indicate the strain culture collection number. In B) the assignment of the highlighted wavenumbers in PC loading plot can be found in the main text. Explained variance of the first five principal components were 82.6%, 13.8%, 1.6%, 1% and 0.5%. Data reproduced from under rights of Creative Commons Attribution 4.0 International License.

**References**

Bolaño Losada C, Slaný O, Byrtusová D, Zimmermann B, Horn SJ, Kohler A, et al. Compatible traits of oleaginous Mucoromycota fungi for lignocellulose-based simultaneous saccharification and fermentation. Biotechnology for Biofuels and Bioproducts. 2025 Dec 1;18(1). doi:10.1186/s13068-025-02621-w

Kosa G, Zimmermann B, Kohler A, Ekeberg D, Afseth NK, Mounier J, et al. High-throughput screening of Mucoromycota fungi for production of low- and high-value lipids. Biotechnol Biofuels. 2018;11(1):1–17. doi:10.1186/s13068-018-1070-7
